# Opposite and complementary roles of the two calcium thresholds for inducing LTP and LTD in models of striatal projection neurons

**DOI:** 10.1101/2025.11.17.688856

**Authors:** Daniel Trpevski

**Affiliations:** Science for Life Laboratory, Division of Computational Science and Technology, School of Electrical Engineering and Computer Science, KTH Royal Institute of Technology, Stockholm, Sweden; Brainster NEXT, Skopje, North Macedonia

## Abstract

Synaptic plasticity has been shown to occur when calcium, flowing into the synapse due to incoming stimuli, surpasses a threshold level. This threshold level is modifiable through a process called metaplasticity. Some neurons, such as the striatal projection neurons, use different sources of calcium as the signal for synaptic strengthening (long-term potentiation, LTP) or weakening (long-term depression, LTD), resulting in them having two thresholds for inducing plasticity. In this study, we show that metaplasticity enables synapses undergoing both LTP and LTD during learning to selectively express just one form of plasticity (either LTP or LTD). To show this, we use the linear and nonlinear feature binding problem (FBP and NFBP) because their input patterns share features, exposing synapses to such competing LTP and LTD processes. In particular, we identify opposite and complementary roles of metaplasticity in the two thresholds for inducing LTP and LTD: metaplasticity in one threshold (e.g. LTD) allows synaptic plasticity *of the opposite type* (e.g. LTP) to be properly expressed. This happens because metaplasticity in the LTD threshold protects strengthened synapses from weakening, thus allowing them to persistently increase during learning (and encode learned patterns). Similarly, metaplasticity in the LTP threhsold prevents weakened synapses from strengthening, thus allowing them to persistently decrease. Under more general conditions for triggering metaplasticity, reversal learning can also be solved, demonstrating metaplasticity’s importance for solving the plasticity–stability dilemma. Finally, we show that even though a single calcium threshold is sufficient for solving the FBP, NFBP and reversal learning, two calcium thresholds allow separate control over LTP and LTD.

## Introduction

Learning is thought to be implemented by several processes in the nervous system, one of which is synaptic plasticity - the modification of synapses based on incoming stimuli to the neuron (***Citri and Malenka, 2007; Kandel et al., 2014; Takeuchi et al., 2014; Abraham et al., 2019***). Most often, incoming stimuli cause calcium influx in the synapse, an important “second messenger” ion that triggers synaptic plasticity. However, not all levels of calcium can elicit synaptic plasticity - threshold levels of calcium have been reported above which plasticity is triggered (***Yang et al., 1999; Nishiyama et al., 2000; Cormier et al., 2001; Shindou et al., 2011***). These calcium thresholds for inducing plasticity are themselves modifiable (plastic) through a process called metaplasticity (***Abraham, 2008***). Metaplasticity regulates the conditions under which plasticity occurs, i.e. under which synapses are modified.

In many neurons, such as hippocampal CA1 and CA3 neurons, Purkinje neurons, cerebellar Golgi cells, and projection neurons from the core of the nucleus accumbens, the calcium signal for synaptic strengthening (long-term potentiation, LTP) and for synaptic weakening (long-term depression, LTD) comes from different sources (***Nishiyama et al., 2000; Hunt et al., 2013; Kohda et al., 1995; Binda et al., 2016; Locatelli et al., 2021; Ji and Martin, 2012***). In the direct-pathway striatal projection neurons (dSPNs), a model of which is used in this study, calcium from NMDA receptors (NMDARs) triggers LTP when followed by activation of D_1_ dopamine receptors (D_1_Rs), while calcium from voltage-gated L-type channels (Ca_v_1.3) triggers LTD when followed by inactivation of D_1_Rs (***Shen et al., 2008; Yagishita et al., 2014; Fisher et al., 2017; Fino et al., 2010; Shindou et al., 2011; Iino et al., 2020***). (Note that to trigger LTD, activation of metabotropic glutamate receptors (mGluRs) is also required (***Shen et al., 2008***). This requirement assures that LTD is only elicited in active synapses where glutamate transmission occurs; Ca_v_1.3 channels alone have an unspecific response – they can be active anywhere where the voltage is high enough.) A calcium threshold for triggering LTD in SPNs has been experimentally quantified (***Shindou et al., 2011***). Although a similar threshold for LTP in SPNs has not been reported yet, presumably such a threshold exists, because weaker stimulation protocols do not induce LTP (***Fino et al., 2005***). (However, probably due to the local striatal microcircuitry, a range of different stimulation protocols can induce a range of different plasticity outcomes, as reviewed in ***Reynolds and Wickens*** (***2002***)). A calcium threshold for LTP has been quantified in hippocampal CA1 neurons (***Cormier et al., 2001***).

By regulating when plasticity can be induced, metaplasticity has been proposed to prepare neural circuits and networks for learning, stabilize synaptic weights (prevent them from reaching very high or very low values) and to contribute to cognitive effects of disease when it is dysregulated (***Hulme et al., 2013***). Most computational metaplasticity models have only one modifiable threshold (where synaptic activity below the threshold induces LTD, and above it induces LTP) (***Yger and Gilson, 2015; Bienenstock et al., 1982; Jedlicka et al., 2015***). On the other hand, the important class of cascade models, whose general form can be seen as having two thresholds, has shown that metaplasticity can solve (several versions of) the plasticity-stability dilemma – it protects stored information from being “deleted” (overwritten) by new synaptic activity, while also allowing for flexible relearning when this information is no longer useful (***Fusi et al., 2005; Iigaya, 2016; Farashahi et al., 2017; Khorsand and Soltani, 2017; Jedlicka et al., 2022; Benna and Fusi, 2016***). The cascade model is a stochastic model of a number of synapses (not placed on any neuron) that randomly experience plasticity events. In this article we study the role of metaplasticity in learning using models closer to biological neurons: a multicompartment model of a dSPN, and a simple, biologically-grounded learning rule describing plasticity of the dSPN cortico-striatal synapse, to which we add metaplasticity in the two thresholds (Fig. 1A). Using the linear and nonlinear feature binding problem (FBP and NFBP), whose structure, as will be explained later, is particularly suitable to emphasize the role of each threshold, we show that the two thresholds have specific, opposite and complementary roles in learning. Further, using reversal learning in the FBP and NFBP, we show that metaplasticity can also solve the plasticity-stability dilemma. Lastly, we show that while a single threshold is sufficient to solve the FBP, NFBP, and reversal learning, two calcium thresholds allow separate control over synaptic strengthening and weakening.

**Figure 1.**
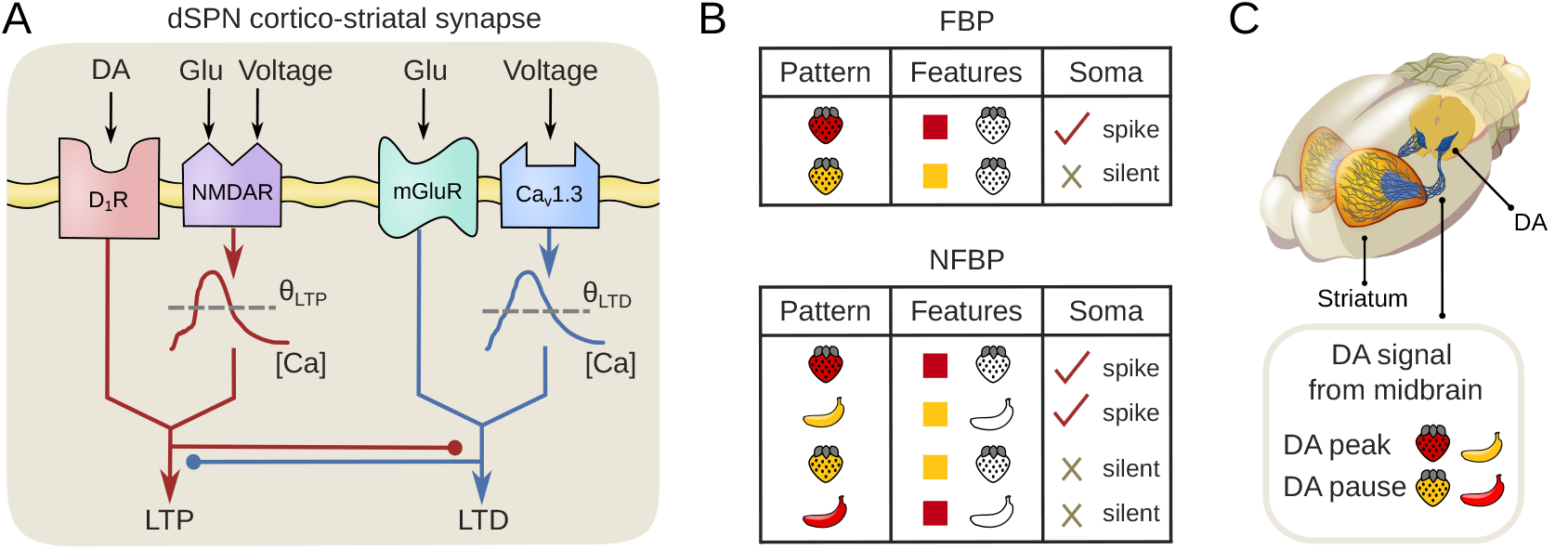
Synaptic plasticity of the cortico-striatal synapse, the tasks for learning, and the dopamine signals for the two types of patterns in the tasks. (A) Scheme describing the requirements for synaptic plasticity at the cortico-striatal synapse in dSPNs (adapted from ***Shen et al. (2008)*** by adding the calcium thresholds, which are the focus of this study). (B) The FBP and the NFBP, illustrated using features common to the visual system. SPNs might instead receive, e.g. sensory- and motor-related features. (C) Dopamine signals from the midbrain are assumed to arrive in the striatum after every pattern. Dopamine peaks are emitted after the relevant patterns, and dopamine pauses after the irrelevant patterns. (The midbrain and its projections are not explicitly modeled in this study, dopamine is programatically provided to the SPN a certain time after pattern arrival.) This figure is adapted from ***Khodadadi et al. (2025)***, licensed under CC BY 4.0 (https://creativecommons.org/licenses/by/4.0/). **Figure 1—figure supplement 1**. An example of the NFBP with features relevant for the striatum.

Before presenting the results, we introduce the FBP and NFBP. The FBP refers to combining (binding) an object’s individual features, such as color, shape, or motion, into a unified object (***Roskies, 1999; Treisman, 1999; von der Malsburg, 1999; Hardcastle, 2017)***. The binding problem arises because features are often processed in different brain regions, and possibly at different times, yet they need to be combined to represent/construct the perception of the object they correspond to. In its simplest form, the linear FBP consists of discriminating between two patterns, meaning that a neuron should spike when activated by one of them, and remain silent when activated by the other (Fig. 1B, top). Each pattern consists of two features, e.g. shape and color, and one of the features can have two values (e.g. ‘red’ and ‘yellow’ for the color feature, as opposed to just ‘strawberry’ for the shape feature in Fig. 1B, top table). Each feature excites the neuron equally, and the term “linear” refers to the computational complexity of the problem, meaning that a neuron able to perform simple linear summation (integration) of incoming stimuli, such as synaptic potentials summated (added) at the soma, can solve the problem. The NFBP consists of discriminating between four patterns: the neuron should spike for two patterns and remain silent for the other two (Fig. 1B, bottom). Each pattern also consists of two features, and in the NFBP both features can have two values (‘strawberry’ or ‘banana’ for shape, and ‘red’ or ‘yellow’ for color, shown in the bottom table in Fig. 1B). Again, each feature excites the neuron equally, and the additional layer of complexity comes from the requirement to spike for two different patterns whose features do not overlap (‘red strawberry’ and ‘yellow banana’). One of the theoretical solutions to the NFBP requires supralinear integration of inputs (***Tran-Van-Minh et al., 2015)***. SPNs exhibit such supralinearities in the form of prolonged dendritic voltage elevations called plateau potentials, demonstrated in the next section. (Supralinear integration of inputs can also solve the simpler FBP.)

As mentioned above, in addition to calcium influx, dopamine signals are also required for plasticity in SPNs. Because of this, we assume that after a pattern is presented, dopamine signals are emitted from the midbrain, and they indicate whether the pattern was “rewarding” or not (Fig. 1C). The patterns for which the SPN should learn to spike are called relevant patterns, and those for which it should be silent are called irrelevant patterns. We assume that dopamine peaks are emitted for relevant patterns, and dopamine pauses for irrelevant patterns (Fig. 1C). After stimulation of the synapse, if the required calcium signal has reached its corresponding threshold, the dopamine signal determines the direction of plasticity. Dopamine peaks are assumed to activate the D_1_Rs, enabling LTP, and dopamine pauses are assumed to deplete basal dopamine in the striatum low enough so that D_1_Rs are not activated, enabling LTD (***Yapo et al., 2017)***.

The reason why the FBP and the NFBP are particularly suitable to demonstrate the role of the calcium thresholds is that relevant and irrelevant patterns share features. In the FBP, the shared feature is the ‘strawberry’ shape (Fig. 1B, top). In the NFBP, both features of a pattern are shared, each feature being shared between one relevant and one irrelevant pattern (Fig. 1B, bottom). As a result, during learning, the synapses representing shared features will experience both dopamine signals (peaks and pauses), guiding them towards both strengthening and weakening. As will be shown, metaplasticity is crucial for “choosing” a direction, i.e. directing plasticity into either strengthening or weakening. In particular, we will see that metaplasticity in the LTD threshold is necessary for strengthening, while metaplasticity in the LTP threshold is necessary for weakening, i.e. that the threshold regulating one plasticity outcome is necessary for properly expressing the *opposite* plasticity outcome.

Lastly, as the input nucleus of the basal ganglia, the striatum integrates convergent cortical inputs from sensory, motor, limbic and associative areas, as well as thalamic and neuromodulatory inputs (***Purves et al., 2018)***. Based on these inputs it initiates and invigorates goal-directed actions, such as reaching for one of the relevant patterns in the FBP and NFBP (***Thura and Cisek, 2017)***. The cortico-striatal synapses are thought to be a major site for learning to initiate such actions, thus forming associations between environmental stimuli, internal states and rewarding outcomes (***Filippo et al., 2009)***. Because of this, despite being suitable tasks for studying metaplasticity, the FBP, NFBP, and reversal learning are also relevant tasks for the striatum’s function. Throughout this article, feature binding is illustrated with a commonly used example from the visual system, although diverse features from the other brain areas would be involved in the striatum (an example is given in Figure 1–figure supplement 1).

Before continuing to the results, we note that we recently participated in a related study with a different goal – whether SPNs can solve the NFBP (***Khodadadi et al., 2025)***. That study used the same neuron model in a regime with much less supralinear integration, and with a different, biologically more detailed learning rule, which incorporated metaplasticity in the LTP threshold only (operating according to a different mechanism and where the evoking calcium signal does not diffuse across neuronal compartments). On the other hand, the goal of this article is to study the role of metaplasticity in the two calcium thresholds in learning, for which we use a simpler, yet biologically-based learning rule where both thresholds are plastic.

## Results

### Linear and supralinear integration of inputs by SPNs and corresponding calcium elevations

We first demonstrate linear and supralinear integration in the dSPN model, which are the substrates for solving the FBP and NFBP. Figure 2 compares linear integration of synaptic inputs distributed across the dendrites with supralinear integration of clustered synaptic inputs on one dendrite (Fig. 2A). The number of synaptic inputs is varied from 1 to 30 synapses, and the evoked somatic and spine voltage, as well as the spine [Ca]_NMDA_ and [Ca]_L-type_ (calcium concentrations arising from NMDARs and L-type calcium channels, respectively) are shown in Figs. 2B and 2C. A comparison of the amplitudes of these four quantities between linear and supralinear integration is shown in Fig. 2D. With linear integration, the somatic voltage amplitude increases linearly with the number of synapses (Fig. 2B_1_ and blue line in Fig. 2D_1_). Conversely, the spine voltage amplitude remains approximately the same, because spines are distributed across the dendrites and are activated in the same way (due to the distance between them, spines cannot influence each other’s voltage, Fig. 2B_2_ and blue line in Fig. 2D_2_). The similar amplitudes of the spines’ voltage result in similar [Ca]_NMDA_ and [Ca]_L-type_ elevations (Figs. 2B_3_, B_4_ and blue lines in Figs. 2D_3_, D_4_).

**Figure 2.**
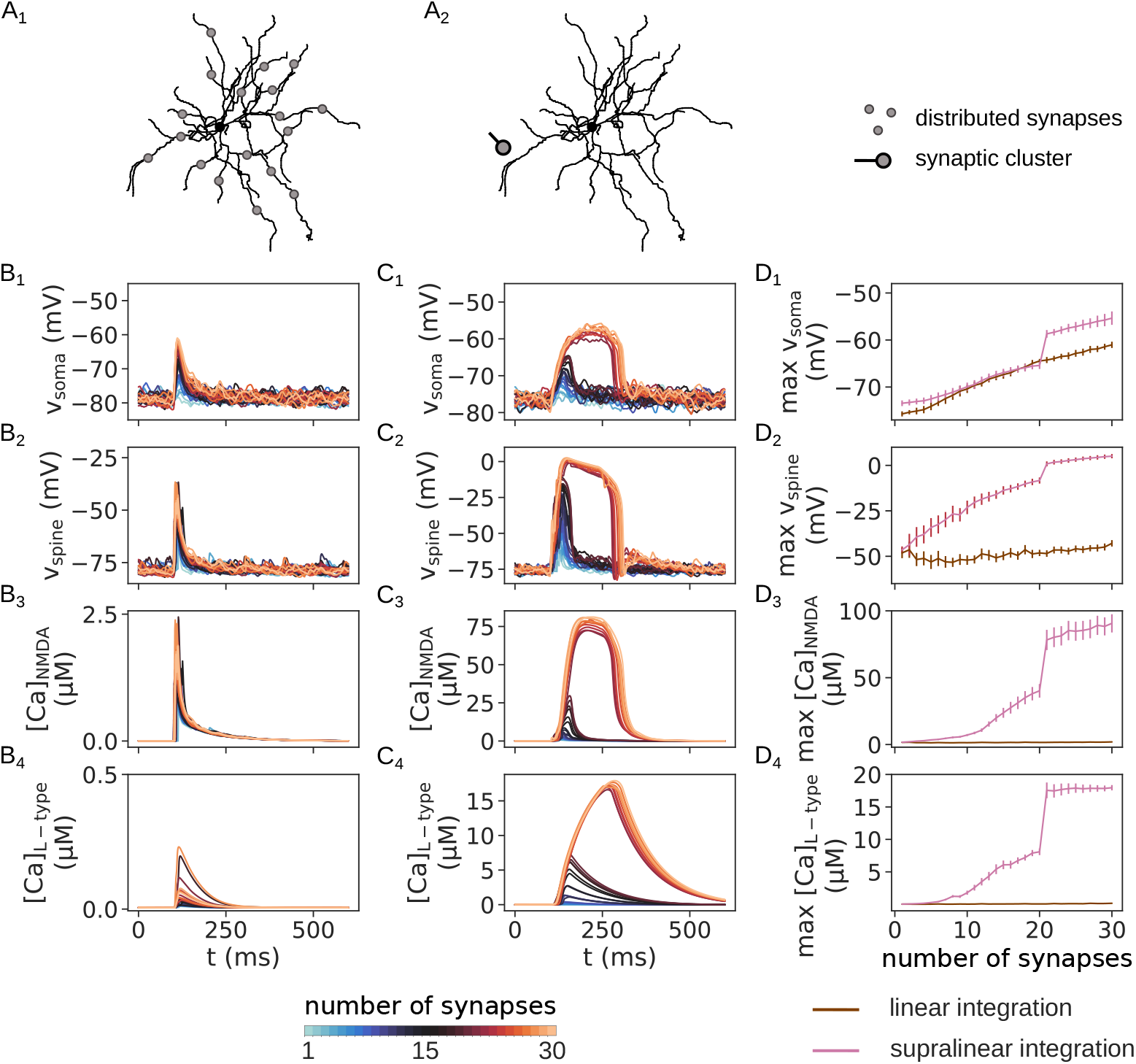
Linear versus supralinear integration of synaptic inputs by SPNs. (A) Illustrations of the two scenarios of linear integration of distributed synaptic inputs by the soma (A_1_), and supralinear integration of clustered synaptic inputs in a dendrite (A_2_). In both scenarios, each synapse is activated with 3 incoming spikes within a window of 30 milliseconds. (B_1_-B_4_) Somatic voltage, spine voltage, spine [Ca]_NMDA_ and spine [Ca]_L-type_ evoked in the linear integration scenario (randomly distributed synapses across the dendrites). Line color indicates the number of distributed synapses. In B_2_-B_4_, traces from a single spine are shown, which was randomly placed on a randomly chosen dendrite. (C_1_-C_4_) Somatic voltage, spine voltage, spine [Ca]_NMDA_ and spine [Ca]_L-type_ evoked in the supralinear integration scenario (synaptic cluster placed on one dendrite, in a 20-micrometer region approximately 120 micrometers away from the soma). Similarly, line color represents the size of the synaptic cluster, which is varied from 1 to 30 synapses. (D_1_-D_4_) The amplitude of the somatic voltage, spine voltage, spine [Ca]_NMDA_ and [Ca]_L-type_ compared between the cases of linear and supralinear integration. Results are averages over 20 trials, and in the case of synaptic clusters, over 8 different dendrites with 20 trials per dendrite (clusters located in a 20-micrometer in a dendritic region starting at approximately between 120 micrometers from the soma).

On the other hand, when synapses are clustered closely together on a dendrite, SPNs can perform supralinear integration of synaptic inputs through plateau potenitals (***Du et al., 2017; Plotkin et al., 2011)***. Plateau potentials are evoked by activating synaptic NMDARs, and experimental studies have suggested that glutamate spillover, which activates extrasynaptic NMDARs, promotes plateaus or is even necessary to evoke them (***Suzuki et al., 2008; Chalifoux and Carter, 2011; Oikonomou et al., 2012)***. We included glutamate spillover because it robustly provides an all-or-none jump in the (somatic) voltage amplitude (***Trpevski et al., 2023)***. This all-or-none supralinearity is important for solving the NFBP (***Tran-Van-Minh et al., 2015)***. Fig. 2C_1_ shows the somatic voltage elevations when varying the size of the synaptic cluster from 1 to 30 synapses, while Fig. 2C_2_ shows the spine voltage in one spine of the synaptic cluster. Figures 2C_3_, C_4_ show the spine [Ca]_NMDA_ and [Ca]_L-type_ that result from stimulating the synaptic cluster. As explained in the Methods section, calcium from these two different sources is tracked separately in the SPN model because they trigger a different plasticity outcome ([Ca]_NMDA_ is needed for LTP and [Ca]_L-type_ is needed for LTD). Also, [Ca]_NMDA_ diffuses in the neuron, while [Ca]_L-type_ does not, since it is assumed to be restricted to microdomains, a form of highly localized signaling often mediated by L-type calcium channels in neurons (***Berridge, 2006; Parekh, 2008; Chen and Sabatini, 2012)***. As the cluster size is increased, increasingly larger and longer voltage elevations are produced as a result of activating more NMDARs (Figs. 2C_1_, C_2_, traces for cluster sizes up to 20 synapses). Once glutamate spillover occurs (here at a cluster size above 20 synapses), an all-or-none plateau potential appears, causing an all-or-none jump in the calcium concentration, as well (Figs. 2C_1_-C_4_).

This all-or-none behavior is also visible as a jump in the amplitudes of the somatic and spine voltage, and the spine [Ca]_NMDA_ and [Ca]_L-type_ in Figs. 2D_1_-D_4_. The somatic and spine plateau amplitude shows roughly linear increases before and after glutamate spillover, with the largest nonlinearity being caused by the plateau (Figs. 2D_1_-D_2_, red lines). The amplitudes of [Ca]_NMDA_ and [Ca]_L-type_ before spillover reflect the voltage dependencies of NMDARs and L-type channels (Figs. 2D_3_-D_4_, red lines for up to 20 synapses), and the largest nonlinearity is again caused by the plateau. The jump in all four quantities in Fig. 2D is an indication of a “threshold supralinearity”. When a critical level of stimulation is achieved, the neuron’s somatic voltage amplitude is suddenly much larger, despite it being stimulated with only one additional synapse in the synaptic cluster. This type of supralinear behavior should in principle be sufficient to solve the NFBP (***Tran-Van-Minh et al., 2015)***.

Also, the comparison of linear and supralinear integration in Fig. 2D_1_ shows that, once a plateau appears, the same number of synapses trigger a much larger somatic voltage elevation when they are clustered compared to when they are randomly distributed across the dendrites. (In the somatic voltage there is also a small difference between linear and supralinear integration when the number of stimulated synapses is less than 10. This arises because the synaptic clusters are always placed at approximately the same distance from the soma, while the distributed synapses can be randomly placed further away.)

### Synaptic plasticity rule with metaplasticity solves both the FBP and NFBP

Before demonstrating the role of metaplasticity using the FBP and NFBP, a learning rule that can solve these two tasks is needed. The synaptic plasticity rule is simple, and is given with Eq. 1. The synaptic weight *w* is increased if, during pattern presentation, [Ca]_NMDA_ reaches a level above the LTP threshold, *θ*_LTP_, and a dopamine peak arrives afterwards. Similarly, the weight is decreased if [Ca]_L-type_ goes above the LTD threshold, *θ*_LTD_, and a dopamine pause arrives afterwards. ([Ca]_NMDA_ and [Ca]_L-type_ are tracked separately in the SPN model, since they produce a different plasticity outcome. Also, as mentioned above, [Ca]_NMDA_ diffuses in the neuron, while [Ca]_L-type_ is assumed to be restricted to microdomains, a form of highly localized signaling often mediated by L-type calcium channels in neurons (***Berridge, 2006; Parekh, 2008; Chen and Sabatini, 2012)***.) Synapses can be increased to a maximal level of *w*_max_, and decreased to a minimal level of *w*_min_, and weight updates are proportional to the current weight value. This formulation is very similar to existing plasticity rules (***Rubin et al., 2001; Kistler, 2002; Gütig et al., 2003)***. The weight updates are also proportional to a learning rate *η*, and the time step for advancing the simulation, Δ*t*. (Note that Δ*t* is neither the timing between pre- and postsynaptic spikes as in spike-timing-dependent rules – this rule does not depend on somatic spiking, nor is it the time until dopamine feedback is provided. It is simply the time step for numerical integration.) This rule is local, meaning that it depends on calcium levels locally at the synapse, and somatic spiking is not necessary for plasticity (similarly to ***Khodadadi et al. (2025)***). This is beneficial for distal synapses, as backpropagating action potentials in general do not affect the local voltage which triggers calcium influx at distant dendritic locations.

Also, at least in hippocampal CA3 pyramidal neurons, it has been reported that local dendritic depolarizations are necessary for plasticity, and somatic spiking is not, consistent with the fact that local calcium levels, driven by local voltage, are the signals triggering plasticity (***Brandalise et al., 2016)***. Similarly, in hippocampal CA1 neurons, local dendritic activity is sufficient for synaptic plasticity (***Bittner et al., 2017)***.

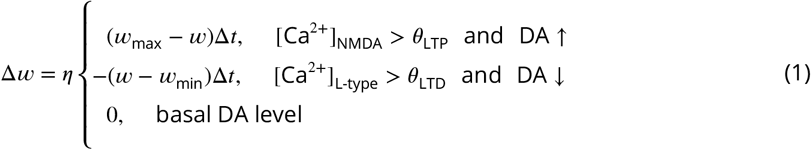

Metaplasticity is also implemented in a simple way, given with Eq. 2. In most of the article, it only occurs if synaptic plasticity also occurs. During one update step, each threshold moves closer to the amplitude (maximum) of the calcium level evoked by an arriving pattern with a rate *η*_*θ*_ (*θ*_LTP_ follows the amplitude of [Ca]_NMDA_, and *θ*_LTD_ follows the amplitude of [Ca]_L-type_). This formulation of metaplasticity is very similar to the sliding threshold in the well-studied BCM learning rule, where the threshold follows a quantity derived from synaptic activity (***Bienenstock et al., 1982)***. When a synapse is modified, both thresholds are updated.

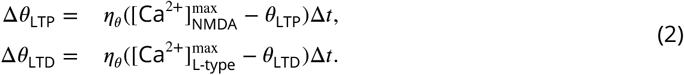

When using supralinear integration of inputs, we have added an upper threshold for LTP, Θ_LTP_ (Eq. 3). This threshold serves to stop synaptic strengthening once synapses are strong enough to evoke a plateau. With it, synapses are not strengthened to the maximal value, *w*_max_, thus avoiding strong supralinear integration for the irrelevant patterns. (This is important for solving the NFBP, as will be seen below.) The upper threshold for LTP only plays a role in synaptic plasticity, and is not taken into account in metaplasticity. This means that metaplasticity is performed in the same way as in Eq. 2, and under the conditions in Eq. 1.

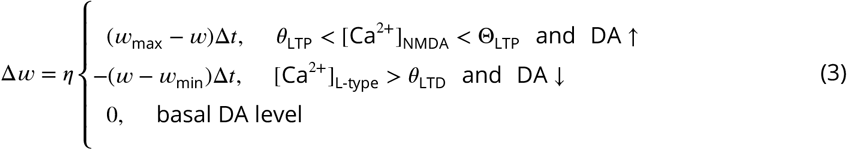

Such “thresholded” metaplasticity, which requires the same conditions to occur as synaptic plasticity, cannot solve reversal learning, a task used to demonstrate the plasticity–stability dilemma. Because of this, when simulating reversal learning, we relaxed the conditions for metaplasticity so that it can be triggered by any calcium level in activated synapses (i.e. no need for calcium to be above a threshold as in Eq. 1).

In summary, each synapse is updated based on its local calcium levels, and has its own LTP and LTD thresholds. During learning, patterns in the FBP and NFBP are presented in random order, and after some time, each pattern is followed by a dopamine feedback signal (a dopamine peak or pause), during which plasticity occurs (Fig. 3A). (The intervals until reward delivery and the next pattern were chosen to shorten simulations; behavioral timescales would likely be longer; see the Methods section for more details.) The learning rule is always active (always “on”), avoiding separate training and testing phases sometimes used in other plasticity rules.

**Figure 3.**
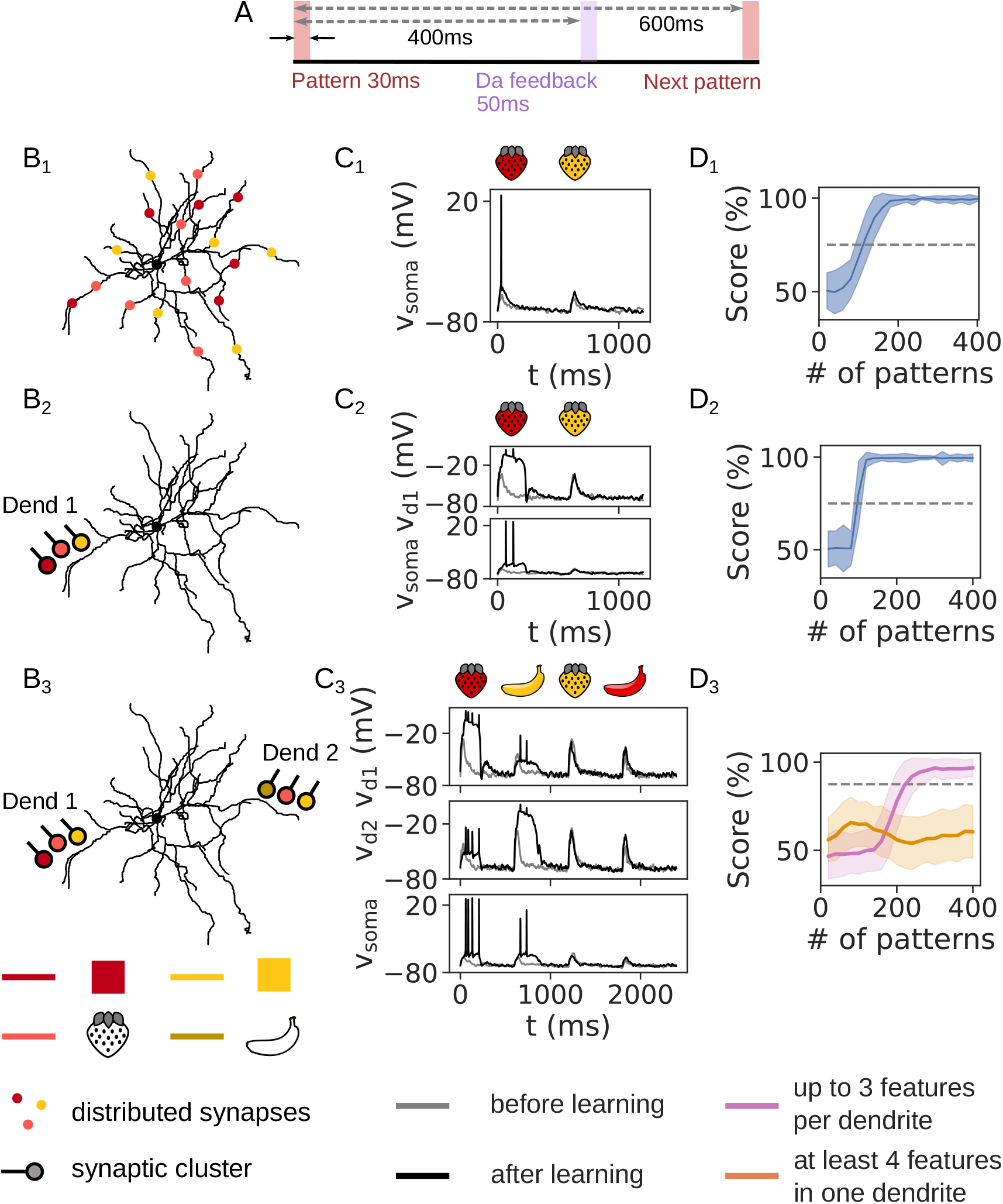
Learning outcome on the FBP and NFBP. (A) Learning protocol. Patterns are presented in random order, with each pattern consisting of synaptic inputs arriving within a 30 ms window (3 spikes per synapse). Synaptic updates occur during a 50 ms dopamine signal delivered 400 ms after pattern onset; the next pattern is presented 600 ms after pattern onset (see Methods). (B) Illustration of the setups for the FBP and NFBP. (B_1_) FBP with linear integration: each feature is represented by 15 synapses distributed across the dendrites. (B_2_) FBP with supralinear integration: 10 synapses per feature, clustered in one dendrite. (B_3_) NFBP (only supralinear integration): 10 clustered synapses per feature, and clusters are located on two different dendrites. (C) The somatic and dendritic voltages in the FBP with linear (C_1_) and supralinear (C_2_) integration, and the NFBP (C_3_), before and after learning (gray and black traces, respectively), elicited by the relevant and irrelevant pattern(s). (D) The performance on the three tasks. Lines indicate score averaged over consecutive blocks of 20 patterns; shaded areas indicate standard deviation, and dashed lines indicate the threshold scores for solving the FBP (75%) and NFBP (87.5%) (see Methods for what these threshold scores represent). In the FBP with linear (D_1_) and supralinear (D_2_) integration, scores are averaged over 50 trials. In (D_2_) the cluster location is randomized across 8 dendrites in each trial; the cluster is placed in a 20-micrometer dendritic region, approximately 120 micrometers away from the soma. In (D_3_), the score is averaged over 12 trials per input configuration, divided into two groups (Figure 3–figure supplement 1), resulting in 216 and 156 trials per group, respectively. In each trial, two dendrites are randomly selected from 8 dendrites to place the clusters, at the same distance as in (D_2_). **Figure 3—figure supplement 1**. All input configurations used in the NFBP.

Before studying the effects of each calcium threshold on learning, we first show that the synaptic plasticity and metaplasticity rule can solve the FBP and NFBP (Fig. 3). For the FBP we have tested both linear integration of randomly distributed synaptic inputs (Figs. 3B_1_-D_1_) and supralinear integration of clustered synapses (Figs. 3B_2_-D_2_), while for the NFBP we tested only supralinear integration of clustered synapses in two different dendrites (Figs. 3B_3_-D_3_), since only supralinear integration can solve the NFBP (***Tran-Van-Minh et al., 2015)***. Figs. 3C_1_-C_3_ show the voltage before and after learning for each pattern in the three scenarios. The SPN is silent before learning, and at the end of learning it has learned to spike to the relevant pattern(s). With supralinear integration, it does this by learning to evoke plateau potentials for the relevant pattern(s). For example, after learning (black traces), ‘red strawberry’ in Figs. 3C_2_ and 3C_3_ elicits a plateau in dendrite 1, which triggers somatic spiking. For the NFBP, the two relevant patterns evoke plateaus on separate dendrites (‘red strawberry’ on dendrite 1 and ‘yellow banana’ on dendrite 2, Fig. 3C_3_). Figs. 3D_1_, D_2_ show the performance on the FBP with linear and supralinear integration averaged over 50 trials, indicating that both modes of integration can solve the task. For the NFBP, in addition to the input configuration in Fig. 3B_3_, with three features on two dendrites, we also tested all possible configurations of four features on two dendrites which allow storage of the relevant patterns on separate dendrites. (These input configurations are listed in Figure 3–figure supplement 1.) Fig. 3D_3_ shows that when up to three features are present in one dendrite, the NFBP is successfully solved. When four features are present in one or both dendrites, usually only one relevant pattern is learned. Solving the NFBP in this case requires additional mechanisms such as branch plasticity or inhibition (***Legenstein and Maass, 2011; Trpevski et al., 2026)***.

Before separately examining the effects of each calcium threshold on learning, it is first necessary to understand how the learning rule successfully solves the tasks by modifying the synaptic weights and thresholds. Figure 4 shows how the synaptic weights and the two calcium thresholds change during learning: they show the same qualitative behavior for both the FBP and NFBP. For the FBP, the synapses carrying the features ‘red’ and ‘strawberry’, which together form the relevant pattern, are strengthened (Figs. 4B_1_, B_2_). The synapses for ‘yellow’ only weaken, since they are only activated as part of the irrelevant pattern (which is followed by a dopamine pause, a signal for LTD). For the NFBP, where each relevant pattern is stored in a separate dendrite, the outcome is similar: the synapses for ‘red’ and ‘strawberry’ on dendrite 1 and the synapses for ‘yellow’ and ‘banana’ on dendrite 2 are strengthened (Figs. 4B_3_, B_4_). The remaining synaptic cluster, belonging to one irrelevant pattern in each dendrite, is weakened (the synapses for ‘yellow’ on dendrite 1 and the synapses for ‘strawberry’ on dendrite 2). In all cases, weakening stops when the evoked [Ca]_L-type_ falls below the LTD threshold. For the FBP with linear integration at the soma, many of the randomly distributed synapses for the relevant pattern are strengthened almost to the maximal value *w*_max_ (Fig. 4B_1_). On the other hand, for the FBP with supralinear integration and the NFBP, the clustered synapses for the relevant pattern(s) are not strengthened to their maximal values. Instead, they stabilize once they are strong enough to evoke a plateau (Figs. 4B_2_–B_4_). This is due to the upper threshold for LTP, Θ_LTP_, shown with dashed lines in Figs. 4C_2_–C_4_. (The stabilization takes about twice as long in the NFBP simply because there are four patterns in the task, i.e. twice as many as in the FBP.) When plateaus are evoked by the relevant patterns, they drive somatic spiking (as indicated above in Figs. 3C_2_, C_3_). In contrast, an irrelevant pattern stimulates only half of the strengthened synapses on one dendrite, which are the synapses representing the shared feature with the relevant pattern (the ‘strawberry’ feature in dendrite 1 in Figs. 4B_2_, B_3_ and the ‘yellow’ feature in dendrite 2 in Fig. 4B_4_). This is not enough for a plateau potential, and even though a supralinear voltage elevation is elicited, no somatic spiking is triggered (Fig. 3C_2_, ‘yellow strawberry’ on dendrite 1 for the FBP and Fig. 3C_3_, ‘yellow strawberry’ on dendrite 1 and ‘red banana’ on dendrite 2 for the NFBP). Nevertheless, these voltage elevations produce high calcium of both types, but, as will be seen below, the shared strengthened synapses are protected from weakening as a result of metaplasticity. Finally, we also show that the NFBP cannot be solved by linear integration – with distributed synapses only one relevant pattern can be stored by the neuron (Figure 4–figure supplement 1).

**Figure 4.**
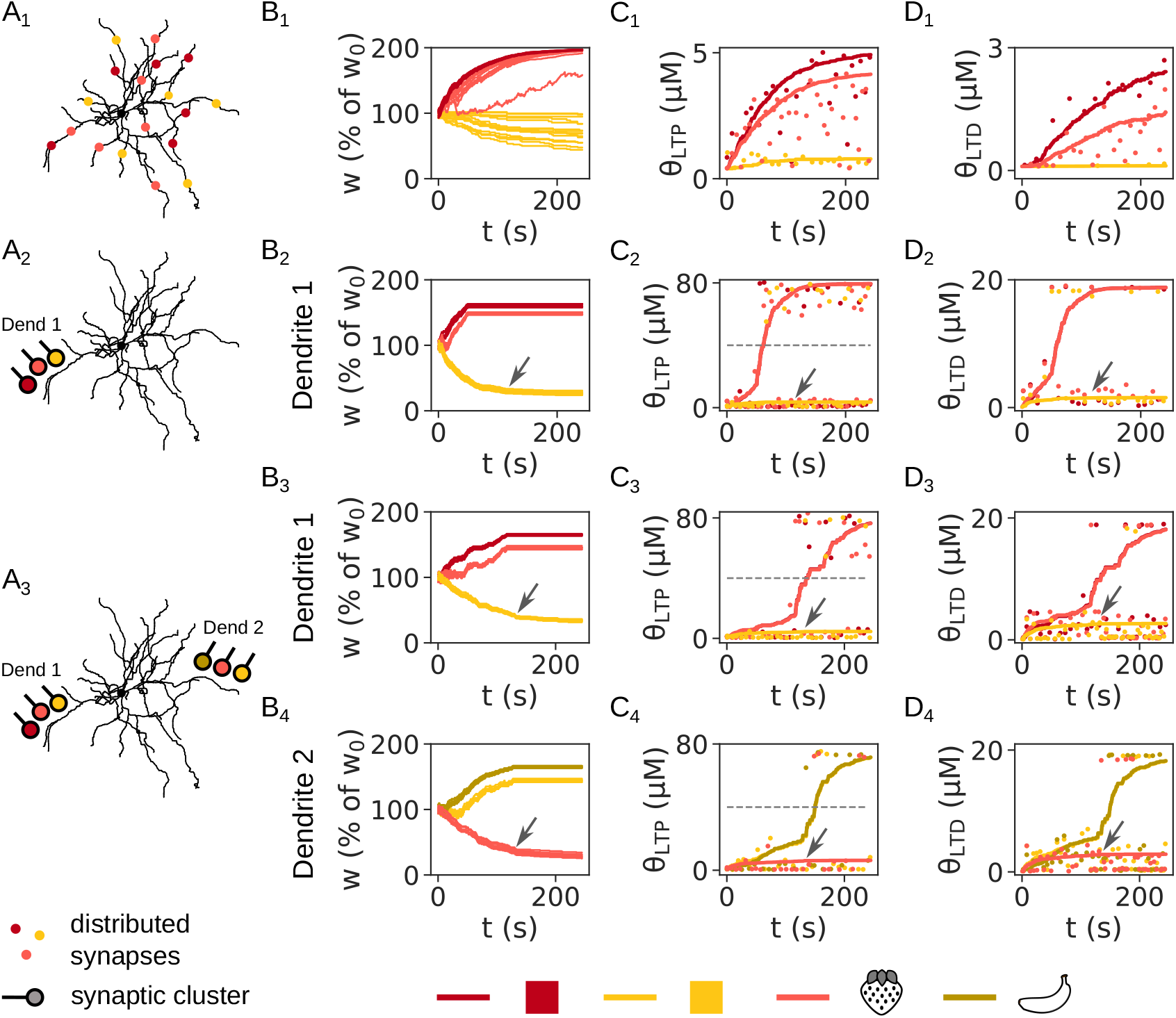
The synaptic weights and calcium thresholds during learning of the FBP and NFBP. (A) Illustrations of the setups for the FBP with linear (A_1_) and supralinear (A_2_) integration, and the NFBP (A_3_) (same as in Fig. 3). (B-D) The evolution of synaptic weights (B), LTP thresholds (C) and LTD thresholds (D) in the FBP with linear (B_1_-D_1_) and supralinear (B_2_-D_2_) integration, and the NFBP (B_3_-D_3_ and B_4_-D_4_). In (B) all synaptic weights for the features are shown, while in (C) and (D) only one synapse per feature is chosen to show its calcium threshold (solid lines). Dashed lines in (C_2_–C_4_) represent the upper threshold for plasticity Θ_LTP_. Dots in (C) and (D) represent the amplitudes of [Ca]_NMDA_ and [Ca]_L-type_, respectively, during pattern presentation. For clarity, the calcium amplitudes for some patterns are omitted. (C) and (D) are shown in larger scale in Figure 4–figure supplement 2, with much more detail in the calcium amplitudes. Arrows show the moment in time when the weakened synapses become low enough for their calcium levels to mostly stay below the calcium thresholds. **Figure 4—figure supplement 1**. The NFBP cannot be solved with linear integration at the soma. **Figure 4—figure supplement 2**. Calcium thresholds and calcium amplitudes from Fig. 4 in greater detail. **Figure 4—figure supplement 3**. The effect of having no upper LTP threshold on supralinear integration. **Figure 4—figure supplement 4**. A parameter scan of *w*_max_ using the rule without an upper LTP threshold. **Figure 4—figure supplement 5**. The maximal weight *w*_max_ prevents unbounded weight growth. **Figure 4—figure supplement 6**. The effect of the learning rate on learning. **Figure 4—figure supplement 7**. The effect of a very low learning rate on learning (*η* = 0.1). **Figure 4—figure supplement 8**. The effect of a very high learning rate on learning (*η* = 20). **Figure 4—figure supplement 9**. The effect of the metaplasticity rate on learning. **Figure 4—figure supplement 10**. The effect of a low metaplasticity rate on learning (*η*_*θ*_ = 0.2). **Figure 4—figure supplement 11**. The effect of a high metaplasticity rate on learning (*η*_*θ*_ = 20).

The LTP and LTD thresholds for just one synapse from each feature are shown in Figs. 4C and 4D, respectively (the same plots are shown with larger panels in Figure 4–figure supplement 2 for better clarity). As mentioned above, they show the same behavior for both the FBP and NFBP, the only difference being that they stabilize at different values due to the different calcium levels evoked by distributed and clustered synaptic inputs. Also, the LTP and LTD thresholds show the same behavior when compared to each other. For example, looking at the strengthened synapses first, they increase to follow the calcium amplitudes evoked by each pattern presentation, and stabilize at these values towards the end of the simulation (thresholds for ‘red’ and ‘strawberry’ tend to the dots of the same color above them in Figs. 4C_1_-C_3_ and 4D_1_-D_3_, and the thresholds for ‘yellow’ and ‘banana’ tend to the dots of the same color above them in Figs. 4C_4_, D_4_; this is better visible in Figure 4–figure supplement 2 and also highlighted with the light blue regions in Figure 4–figure supplement 2B_3_). In the case of linear integration they stabilize at low values, because distributed synapses do not have cooperative voltage effects (due to the large distances between them), and low calcium levels are evoked as a result (Figs. 4C_1_, D_1_). Instead, with supralinear integration, the plateaus evoked by the clustered synapses cause much larger voltage and calcium elevations, so the thresholds rise to much higher levels (Figs. 4C_2_-C_4_ and 4D_2_-D_4_). Note that the LTP threshold *θ*_LTP_ can also go above the upper plasticity threshold Θ_LTP_ (*θ*_LTP_ and Θ_LTP_ are independent), reflecting the assumption that different molecular circuitry implements these two thresholds. (For example, experiments in hippocampal CA1 pyramidal neurons have demonstrated that the two different voltage thresholds for LTD and LTP can reverse their relative positions as a result of metaplasticity: the voltage threshold for LTD can rise to -20 mV, going above the voltage threshold for LTP, which reduced to -30 mV (***Ngezahayo et al., 2000)***).

We next look at the thresholds for the weakened synapses (‘yellow’ synapse in Figs. 4C_1_-C_3_ and 4D_1_-D_3_ and ‘strawberry’ synapse in Figs. 4C_4_, D_4_). At the beginning, the thresholds increase. Later, once the synapses are weakened enough so that the evoked [Ca]_L-type_ falls below the LTD threshold, they reach a stable level. For example, in Figs. 4C_2_, C_3_ and 4D_2_, D_3_, at the beginning of learning, yellow dots can be seen above the ‘yellow’ thresholds, while later they almost always remain below the thresholds. Similarly, in Figs. 4C_4_, D_4_, light red dots are at first seen above the ‘strawberry’ thresholds, while later they are found below them. This transition of the [Ca]_L-type_ amplitudes from above to below the LTD thresholds is marked with the arrows in Figs. 4B_2-4_-D_2-4_, corresponding to the moment in time where synapses are weakened enough. (This is more clearly visible in Figure 4–figure supplement 2, and it is also highlighted with the green and brown regions in Figure 4–figure supplement 2B_3_, and the pink region where yellow dots are absent.) We mention that in the FBP, where the weakened synapses are only activated by the irrelevant pattern, both of their thresholds are only updated during LTD. On the other hand, in the NFBP the weakened synapses and their thresholds can be updated during both LTP and LTD, because these synapses are also activated by a relevant pattern (i.e. are shared with a relevant pattern). (However, in Fig. 4 this only happens at the very beginning, when the weakened synapses also undergo LTP. As they are being weakened, their LTP threshold is raised, so [Ca]_NMDA_ amplitudes later do not go above the LTP threshold.) In summary, once a calcium amplitude falls below its threshold, plasticity and metaplasticity cannot be triggered, and the thresholds, as well as the synapses, are not updated anymore. In other words, once the thresholds reach the evoked calcium amplitudes, synapses are stabilized. In effect, metaplasticity acts as a lock on the synapses: synapses are flexible for learning in the time it takes for the thresholds to reach the calcium amplitudes, and are stabilized once that occurs. Accordingly, we have assumed that at the beginning of learning thresholds start at a low value, so that synapses are initially flexible to store information. All of this will be much more evident in the next section, where we study the role of metaplasticity.

We stress once again that in the FBP with supralinear integration, one of the strengthened synaptic clusters belongs to the shared feature (the ‘strawberry’ cluster in Figs. 4A_2_, B_2_), and in the NFBP both strengthened synaptic clusters belong to shared features (the ‘red’ and ‘strawberry’ clusters in dendrite 1 in Figs. 4A_3_, B_3_, and the the ‘yellow’ and ‘banana’ clusters in dendrite 2 in Figs. 4A_3_, B_4_). Importantly, the LTD threshold for these synapses is raised because of the plateaus (Figs. 4D_2_–D_4_). Because of this, the high [Ca]_L-type_ levels in these synapses evoked from an irrelevant pattern are below the LTD threshold and are not enough to weaken them (e.g. light red dots in Figs. 4D_2_, D_3_ just above the ‘yellow’ threshold after 100 seconds are far below their ‘strawberry’ threshold, and dark red dots are far below their ‘red’ threshold in Fig. 4D_3_; similarly yellow and dark yellow dots in Fig. 4D_4_ located just above the ‘strawberry’ threshold after 100 seconds are far below their ‘yellow’ and ‘banana’ thresholds, respectively; dots are also highlighted in the pink region of Figure 4–figure supplement 2B_3_). In this way, these strengthened synapses belonging to a shared feature are protected from weakening, allowing them to store one part of the relevant pattern. This effect will also be much more evident in the next sections, where we look at the effects of metaplasticity on learning.

#### The effects of parameters in the learning rule

We have also examined the effects of parameters in the learning rule. In supralinear integration, the upper plasticity threshold Θ_LTP_ prevents the strengthened synapses from saturating at their maximal levels. Without it, the FBP is still solved, even though weights attain their maximal values (Figure 4–figure supplements 3B_1_ and 3E_1_). It is successfully solved because the features innervate only one dendrite, and the saturated weights do not cause somatic spiking when activated by the irrelevant pattern (Figure 4–figure supplement 3A_1_). However, the NFBP is not solved without an upper threshold Θ_LTP_, because the features innervate two dendrites, and the maximally strengthened synapses cause supralinear voltage elevations in the dendrites for the irrelevant patterns, which trigger somatic spiking (Figure 4–figure supplements 3A_2_, 3B_2_, 3B_3_ and 3E_2_). The maximally strengthened synapses also drive more weakening in the weakened synapses, compared to when an upper threshold is present (Figure 4–figure supplement 3B, cf. weakened weights in Fig. 4B_2_– B_4_). This is due to the elevated [Ca]_L-type_ evoked when an irrelevant pattern activates the maximally strengthened synapses (shared “strawberry” synapses in Figure 4–figure supplements 3B_1_, B_2_ and shared “yellow” synapses in Figure 4–figure supplement 3B_3_). The elevated [Ca]_L-type_ for the irrelevant patterns is also visible in the elevated LTD thresholds for the weakened synapses (“yellow” synapse in Figure 4–figure supplement 3D_1_, D_2_ and “strawberry” synapse in Figure 4–figure supplement 3D_3_, cf. Figs. 4D_2_–D_4_). (The LTP thresholds for the weakened synapses are also elevated, due to higher [Ca]_NMDA_ produced by the maximally strengthened synapses. This is visible in Figure 4–figure supplement 3C when compared to Figs. 4C_2_–C_4_).

The upper threshold Θ_LTP_ is necessary to solve the FBP and NFBP because there is no single value of the maximal weight *w*_max_ with which both tasks are solved simultaneously, as can be seen from the parameter scan for *w*_max_ in Figure 4–figure supplement 4 done in the absence of Θ_LTP_. The FBP with supralinear integration and the NFBP are first solved for 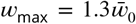 (due to the *w*_max_ threshold for glutamate spillover being between 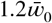 and 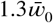 in this study, and spillover evoking plateaus), but this value is not high enough to solve the FBP with linear integration (Figure 4–figure supplement 4, second column). A value of 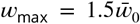 is needed to solve the FBP with linear integration, which, on the other hand, is too high to solve the NFBP (the performance on the NFBP drops to the threshold score for solving it in Figure 4–figure supplement 4, fourth column). All other values for 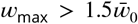 increase the performance on the FBP with linear integration, and cause faster learning of the FBP with supralinear integration, but lower the performance on the NFBP due to spiking for the irrelevant patterns, as shown previously in Figure 4–figure supplement 3A_2_. This suggests that if a range of *w*_max_ values exists that solves all three tasks, it is very narrow, and such a solution is not robust to variations in *w*_max_.

The maximal weight *w*_max_ prevents unbounded weight growth, ensuring stability of the neuron’s firing rate in the FBP with linear integration (Figure 4–figure supplement 5). Without *w*_max_, the weights grow until [Ca]_NMDA_ saturates in the spines (which does not occur for most synapses within the simulation time in Figure 4–figure supplement 5B; also the weights grow to unphysiological values, which of course does not happen in reality). Thus, in the FBP with linear integration, the neuron enters depolarization block when stimulated by a pattern after learning (Figure 4–figure supplement 5A_1_). On the other hand, in the FBP with supralinear integration, firing rate stability is ensured by the plateau potentials, since a plateau’s voltage is limited by the NMDAR reversal potential, regardless of how high the synaptic weights are (Figure 4–figure supplement 5A_2_). This is part of the plateaus’ function to provide dynamic range compression, as shown experimentally in Figs. 3E, 4D and 12 in ***Oikonomou et al. (2012)***.

The learning rate *η* has an effect on the speed of learning – more pattern presentations are required with lower learning rates, and vice versa (Figure 4–figure supplement 6). However, if the learning rate *η* is very low or very high, the NFBP cannot be solved: a very low *η* causes spiking for the irrelevant patterns because synapses are not weakened enough (Figure 4–figure supplement 7), and a very high *η* causes large updates to the synaptic weights, disrupting learning due to the similarly large changes in calcium, which can easily fall below the thresholds (Figure 4–figure supplement 8). Within a certain range, the rate of threshold adaptation (metaplasticity rate) *η*_*θ*_ has no significant effect on learning to solve the FBP and NFBP (Figure 4–figure supplement 9). If the metaplasticity rate is too low, it takes a long time for the thresholds to reach the calcium amplitudes, and as a result, synapses take a longer time to stabilize in the FBP with linear integration and in the NFBP (Figure 4–figure supplement 10B_1_, 10B_3_, 10B_4_). Synapses stabilize quickly in the FBP with supralinear integration because it is sufficient that one of the clusters is strengthened enough to trigger plateaus (Figure 4–figure supplement 10B_2_). Learning is slowed down in the NFBP – more patterns are needed before the same performance level is reached (Figure 4–figure supplement 10E_3_, cf. Fig. 3D_3_), while learning in the FBP is unaffected (Figure 4–figure supplements 10E_1_, E_2_, cf. Figs. 3D_1_, D_2_). On the other hand, if the metaplasticity rate is too high, the thresholds reach the respective calcium levels quickly, shortening the period where synapses are flexible and preventing them from being changed very much, i.e. they are quickly stabilized (Figure 4–figure supplement 11B–D). This results in more variable performance on the FBP with linear integration because weights are not strengthened enough (Figure 4–figure supplement 11E_1_), and in reduced performance on the NFBP because weakened clusters are not weakened enough, causing spiking for the irrelevant patterns (Figure 4–figure supplement 11E_3_).

### Metaplasticity in the LTD threshold prevents weakening of strengthened synapses, protecting learned patterns

The role of each threshold is easily examined with this plasticity and metaplasticity learning rule by turning metaplasticity off, i.e. fixing a threshold’s value during learning. As mentioned in the introduction, the FBP and NFBP are particularly useful for demonstrating the role of each threshold because the patterns in the tasks share features. In the FBP, the relevant and irrelevant pattern (the red and yellow strawberry) share the shape feature (the feature “strawberry”). This means that during learning, the synapses representing this feature experience both strengthening and weakening (e.g. visible at the beginning of learning in Figs. 4B_1_, B_2_), and that strengthening is somehow “chosen”. Fixing the LTD threshold (to the initial low value in the simulation) shows that the shared synapses cannot strengthen properly, i.e. cannot stabilize (Figs. 5B_1_, B_2_). This happens because there is no metaplasticity in the LTD threshold – it is fixed, i.e. it never rises to follow [Ca]_L-type_. As a result, whenever an irrelevant pattern arrives, it always causes higher [Ca]_L-type_ than the fixed LTD threshold, and can always weaken the shared synapses, despite them also being strengthened by the relevant pattern (Figs. 5D_1_, D_2_ – LTD thresholds stay fixed during learning, and [Ca]_L-type_ is always above *θ*_LTD_). The other two features are not shared – the feature “red” is only activated by the relevant pattern and, the feature “yellow” only by the irrelevant pattern. As a result, they can only ever experience plasticity in one direction: the feature “red” is strengthened, and the feature “yellow” is weakened. The fixed LTD threshold has no influence on the strengthened feature (“red”). However, in the FBP with supralinear integration, where the cooperative voltage effects of the clustered synapses produce higher calcium elevations, it does influence the weakened feature (“yellow”): because the LTD threshold is fixed to a low value, the evoked [Ca]_L-type_ is high enough to decrease the weakened synapses much more compared to when metaplasticity in the LTD threshold is active (Figs. 5B_1_, B_2_, cf. “yellow” synapses in Figs. 4B_1_, B_2_).

**Figure 5.**
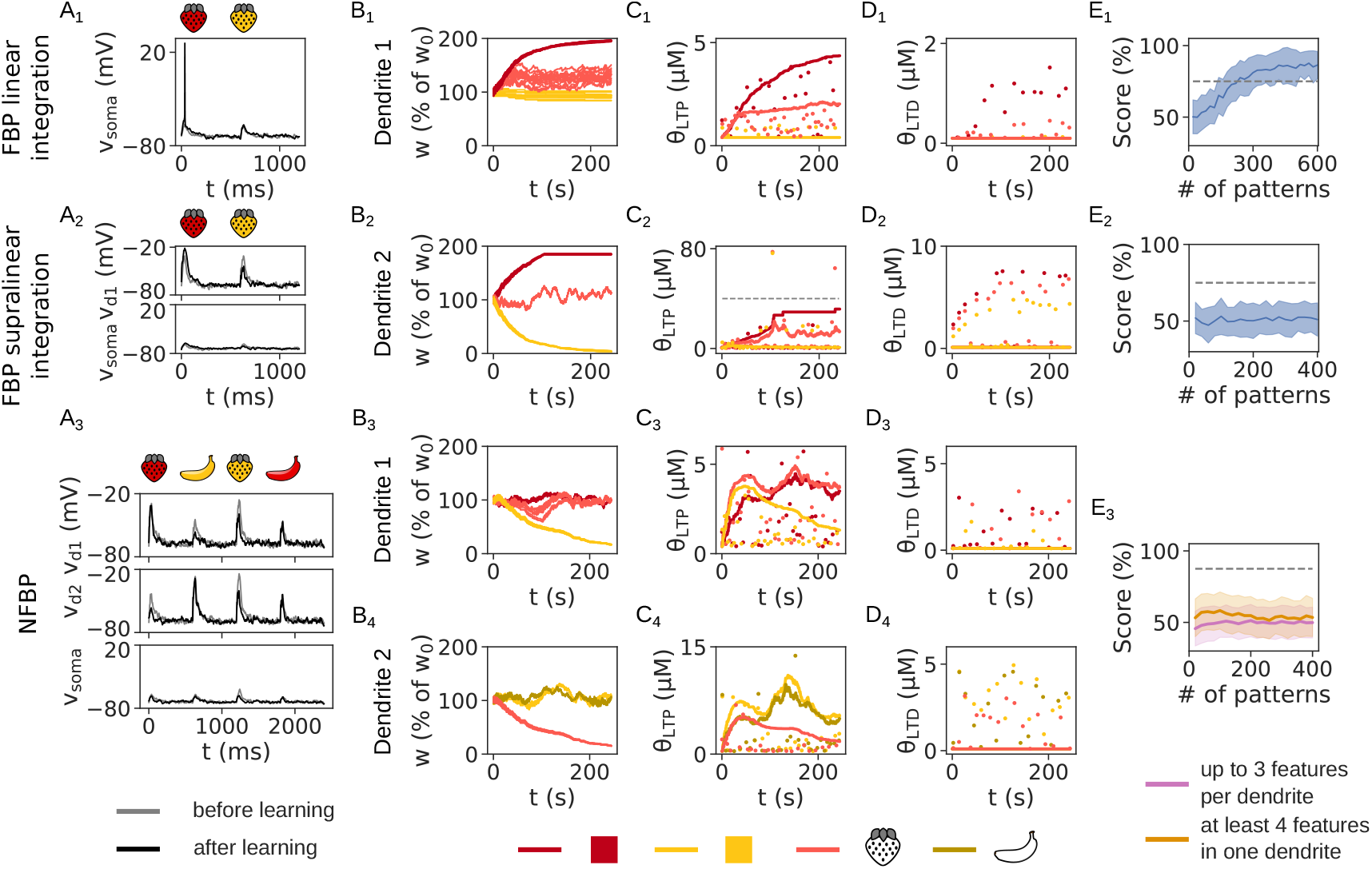
Learning outcome on the FBP and NFBP without metaplasticity in the LTD threshold. (A) The dendritic and somatic voltage before (gray traces) and after learning (black traces) for the FBP with linear (A_1_) and supralinear integration (A_2_) and the NFBP (A_3_). (B–D) The evolution of synaptic weights (B), LTP thresholds (C) and LTD thresholds (D). (D) shows the fixed LTD thresholds. In (B) all synapses are shown, and in (C, D) only one synapse per feature is used to show its calcium threshold. Dots represent the amplitudes of [Ca]_NMDA_ (C) and [Ca]_L-type_ (D) during pattern presentation, omitting some patterns for clarity. Dashed line in (C_2_) shows the upper threshold for LTP, Θ_LTP_. (E) The performance on the FBP with linear (E_1_) and supralinear integration (E_2_) and the NFBP (E_3_) without metaplasticity in the LTD threshold. Performance on the FBP is averaged over 50 trials, and in the NFBP over 12 trials per input configuration (divided into the two groups shown in Figure 3–figure supplement 1), with 216 and 156 trials, respectively).

Regarding the performance, in the FBP with linear integration, despite the shared synapses not stabilizing, the strengthening is enough to drive somatic spiking (Fig. 5A_1_), and the task is solved but with a lower performance (Fig. 5E_1_). On the other hand, in the FBP with supralinear integration, because the shared “strawberry” synapses do not stabilize, the clustered synapses are not strong enough to evoke plateaus (Fig. 5A_2_), and the FBP is not learned (Fig. 5E_2_).

The effect of the fixed LTD threshold on the NFBP is analogous, but even more prominent because in the NFBP, a relevant pattern shares its features with both irrelevant patterns (e.g. “red strawberry” shares the color feature with “red banana” and the shape feature with “yellow strawberry”). This means that during learning, both features of a relevant pattern experience strengthening and weakening. Hence, when the LTD threshold is fixed, none of their synapses stabilize, and the relevant patterns are not stored in the dendrites (neither “red” and “strawberry” in dendrite 1, Fig. 5B_3_, nor “yellow” and “banana” in dendrite 2, Fig. 5B_4_). As a result, they cannot evoke plateaus, and the NFBP is not learned (Figs. 5A_3_, E_3_). The LTP threshold follows the level of [Ca]_NMDA_ and also does not stabilize (Figs. 5C_3_, C_4_). As with the FBP, this happens because there is no metaplasticity in the LTD threshold, and irrelevant patterns always cause higher [Ca]_L-type_ than the fixed LTD threshold (Figs. 5D_3_, D_4_). This means that despite being strengthened, these synapses will always get weakened. Conversely, when metaplasticity in the LTD threshold is active, it follows the increased levels of [Ca]_L-type_ elicited by a plateau (evoked due to prior synaptic strengthening, as shown previously in Figs. 4D_2_–D_4_). This elevated LTD threshold protects the strengthened synapses from weakening when they are activated by an irrelevant pattern. As mentioned above, they are protected because an irrelevant pattern, activating only half of the strengthened synapses, does not evoke a plateau, resulting in [Ca]_L-type_ lower than the elevated LTD threshold in Figs. 4D_2_–D_4_ (e.g. light red dots in pink region in Figure 4–figure supplement 2B_3_ are below the elevated “strawberry” threshold).

Without metaplasticity in the LTD threshold, strengthened synapses are not protected from weakening. Also, because the LTD threshold is fixed to a low value, the synapses belonging to the weakened feature in each dendrite are almost weakened to the minimal value *w*_min_. In summary, by protecting synapses from weakening, the *threshold for weakening* allows proper synaptic *strengthening*.

### Metaplasticity in the LTP threshold prevents strengthening of weakened synapses, promoting the weakening of unnecessary synapses

On the other hand, the LTP threshold has an opposite effect on learning. This effect can be seen in a shared feature which ultimately needs to be weakened, but which experiences a drive to both strengthen and weaken during learning. There is no such feature in the FBP (the only shared feature, “strawberry”, needs to be strengthened), so fixing the LTP threshold has no effect on solving this task (Figs. 6A_1_–E_1_ and 6A_2_–E_2_, cf. Figs. 3A_1,2_–C_1,2_, 4A_1,2_–D_1,2_). Nevertheless, there are such features in the NFBP, which is why it is a suitable task to demonstrate the role of the LTP threshold. In fact, each irrelevant pattern in the NFBP also shares its two features with the relevant patterns, and, as a result, they also experience both strengthening and weakening during learning (e.g. the features “yellow” in dendrite 1 and and “strawberry” in dendrite 2 in Fig. 4A_3_).

**Figure 6.**
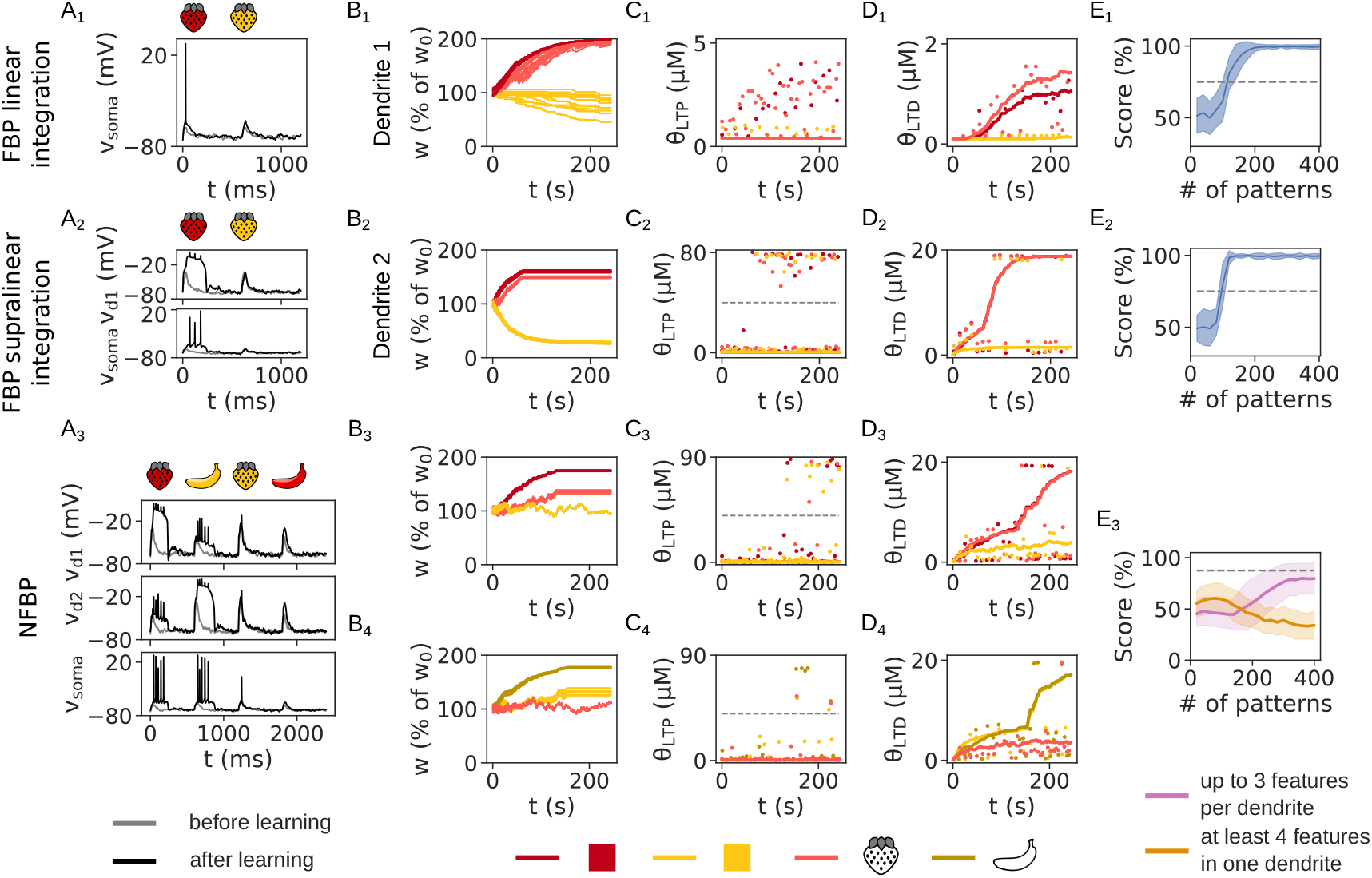
Learning outcome on the FBP and NFBP without metaplasticity in the LTP threshold. (A) The dendritic and somatic voltage before (gray traces) and after learning (black traces) for the FBP with linear (A_1_) and supralinear integration (A_2_) and the NFBP (A_3_). (B–D) The evolution of synaptic weights (B), LTP thresholds (C) and LTD thresholds (D) for the synapses in each dendrite. (C) shows the fixed LTP thresholds, and the dashed lines show the upper LTP threshold, Θ_LTP_. In (B) all synapses are shown, and in (C, D) only one synapse per feature is used to show its calcium threshold. Dots represent the amplitudes of [Ca]_NMDA_ (C) and [Ca]_L-type_ (D) during pattern presentation, omitting some patterns for clarity. Dashed lines in (C_2_–C_4_) show the upper threshold for LTP, Θ_LTP_. (E) The performance on the FBP with linear (E_1_) and supralinear integration (E_2_) and the NFBP (E_3_) without metaplasticity in the LTP threshold. Performance on the FBP is averaged over 50 trials, and in the NFBP over 12 trials per input configuration (divided into the two groups shown in Figure 3–figure supplement 1, with 216 and 156 trials, respectively).

If the LTP threshold is fixed, the features belonging to the irrelevant pattern in each dendrite cannot be weakened properly (“yellow” synapses in dendrite 1 in Fig. 6B_3_ and “strawberry” synapses in dendrite 2 in Fig. 6B_4_). Because of the low value of the LTP threshold, whenever an irrelevant pattern activates these synapses, the evoked [Ca]_NMDA_ is still greater than the threshold (Figs. 6C_3_, C_4_), triggering synaptic strengthening even if synapses have previously been weakened. This means that when metaplasticity in the LTP threshold is active, by following the levels of [Ca]_NMDA_ (e.g. LTP threshold for “yellow” in Fig. 4C_3_ and for “strawberry” in Fig. 4C_4_), it prevents these synapses from strengthening when they are activated by the relevant pattern (which can only be stored fully in the other dendrite). As a result, the synapses for “yellow” in Fig. 4B_3_ and for “strawberry” in Fig. 4B_4_ are successfully weakened, while in Figs. 6B_3_, B_4_ they are not. (The LTD threshold behaves as usual for the strengthened synapses, protecting them from weakening, as shown in Figs. 6B_3_, B_4_ and 6D_3_, D_4_. However, for the weakened synapses, it does not stabilize.) Without metaplasticity in the LTP threshold, the NFBP is on average not solved (Fig. 6E_3_). Despite both relevant patterns being stored by each dendrite, the performance is lowered because the SPN sometimes also spikes for the irrelevant patterns (Fig. 6A_3_). Also, in the input configurations containing four features in one dendrite, the average performance even drops below 50 % (green trace in Fig. 6E_3_). In summary, by preventing synapses from strengthening, the *threshold for strengthening* allows proper synaptic *weakening*.

### Both thresholds need to be updated during one synaptic modification in order to solve the NFBP

The rule is formulated so that during one synaptic modification (e.g. LTP) the thresholds for both LTP and LTD are updated. Alternatively, it could be formulated so that during LTP, only the threshold for LTP is updated, and during LTD, only that for LTD is updated. We test this variant in this section, and the results from the previous two sections already give an intuition how this variant would work. This time we start with the NFBP, since that is the task where the role of both thresholds can be demonstrated. Updating only the corresponding threshold during one synaptic modification means that the other threshold is, in effect, kept fixed during that plasticity update, making metaplasticity only partially active. The effects are seen in Figs. 7C_1_, C_2_. Synapses are neither properly strengthened, nor weakened, at least within the length of the simulation. For example, one of the relevant patterns (“red strawberry”) shows a trend of strengthening, but is also being weakened (Fig. 7C_1_). This is because as the LTD thresholds for “red” and “strawberry” move higher (Fig. 7E_1_), LTD in these synapses is (sometimes) prevented, and a trend of synaptic strengthening appears. Also, as the LTP threshold for “yellow” increases (Fig. 7D_1_), it prevents the “yellow” synapses from strengthening, and as a result, they weaken to a degree.

**Figure 7.**
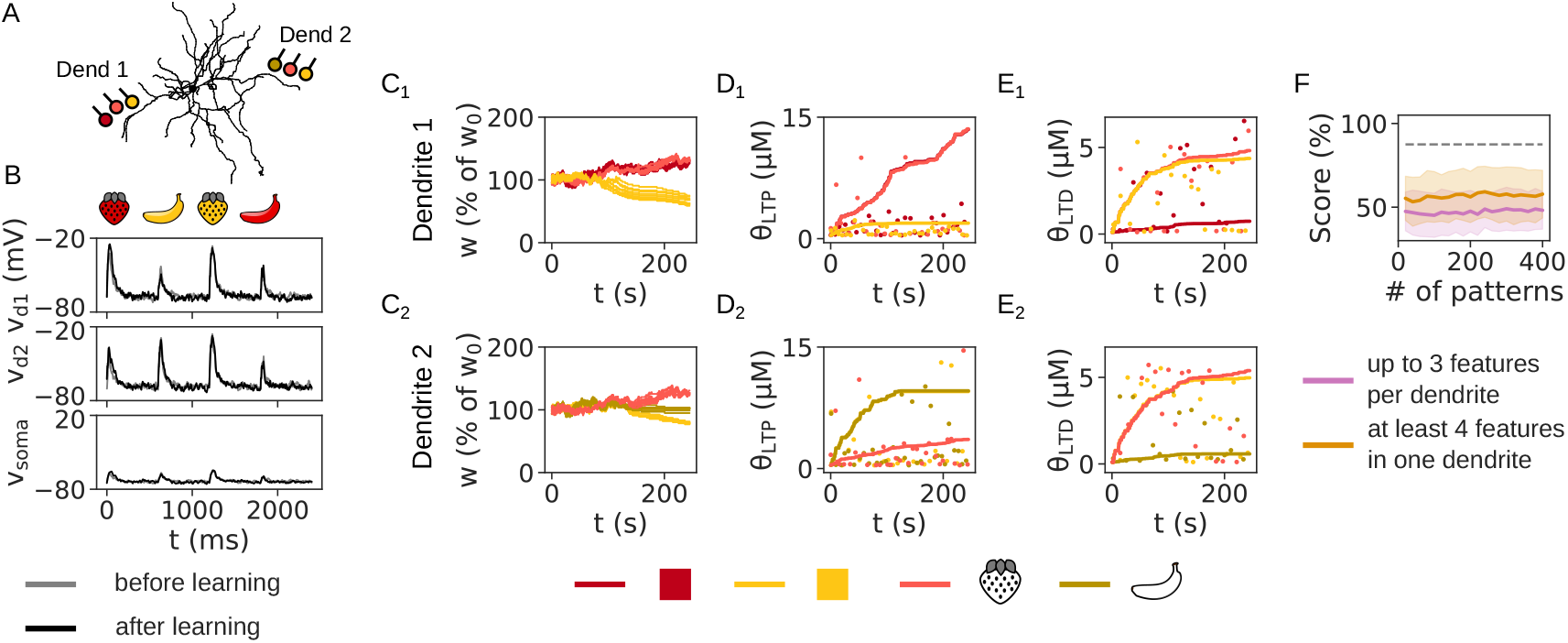
Learning outcome on the NFBP with partial metaplasticity, where the LTP threshold is updated only during LTP, and the LTD threshold is updated only during LTD. (A) An illustration of the setup for the NFBP (same as in Fig. 3B_3_). (B) The dendritic and somatic voltage before (gray traces) and after learning (black traces) for all four patterns. (C-E) The evolution of synaptic weights (C), LTP thresholds (D) and LTD thresholds (E) for the synapses in each dendrite. In (C) all synapses are shown, and in (D, E) only one synapse per feature is used to show its calcium threshold. Dots represent the amplitudes of [Ca]_NMDA_ (D) and [Ca]_L-type_ (E) during pattern presentation, omitting some patterns for clarity. Dashed lines in (C_2_–C_4_) show the upper threshold for LTP, Θ_LTP_. (F) The performance on the NFBP with partial metaplasticity, averaged over 12 trials per input configuration (divided into the two groups shown in Figure 3–figure supplement 1, with 216 and 156 trials, respectively). **Figure 7—figure supplement 1**. The effect of partial metaplasticity on solving the FBP.

However, since the LTD threshold is not updated during LTP, it does not follow the [Ca]_L-type_ levels generated by the relevant patterns. It is updated only during LTD, following the [Ca]_L-type_ levels from irrelevant patterns. For example, in dendrite 1, the “red banana” activates only the cluster representing “red”, evoking little [Ca]_L-type_ and, as a result, the LTD threshold for the “red” synapse is lower (Fig. 7E_1_). On the other hand, the “yellow strawberry” activates two synaptic clusters (Fig. 7A_1_), causing a larger voltage elevation and consequently larger [Ca]_L-type_, so the LTD thresholds for these synapses move higher (Fig. 7E_1_). However, because of the trend of synaptic strengthening in the “red” and “strawberry” synapses, the irrelevant patterns sometimes still evoke [Ca]_L-type_ higher than the LTD thresholds. As a result, the synapses for “red” and “strawberry” in dendrite 1 are not protected from weakening (when activated by the irrelevant patterns). Also, as the LTD threshold for “yellow” increases in the first 70 s of the simulation (Fig. 7E_1_), the synapses for “yellow” show a trend of increasing in this period, as well, before being decreased afterwards (when the “yellow” LTP threshold is increased). On the other hand, in dendrite 2 the relevant pattern is not strengthened, and it is not clear whether more learning trials will cause both relevant patterns to be stored in the two dendrites. On average, the NFBP is not solved in this case (Fig. 7F).

Finally, this variant is similar to not having metaplasticity at all (Fig. 8). Without metaplasticity, no synapses stabilize, i.e. they are neither strengthened nor weakened (Fig. 8C_1_, C_2_).

**Figure 8.**
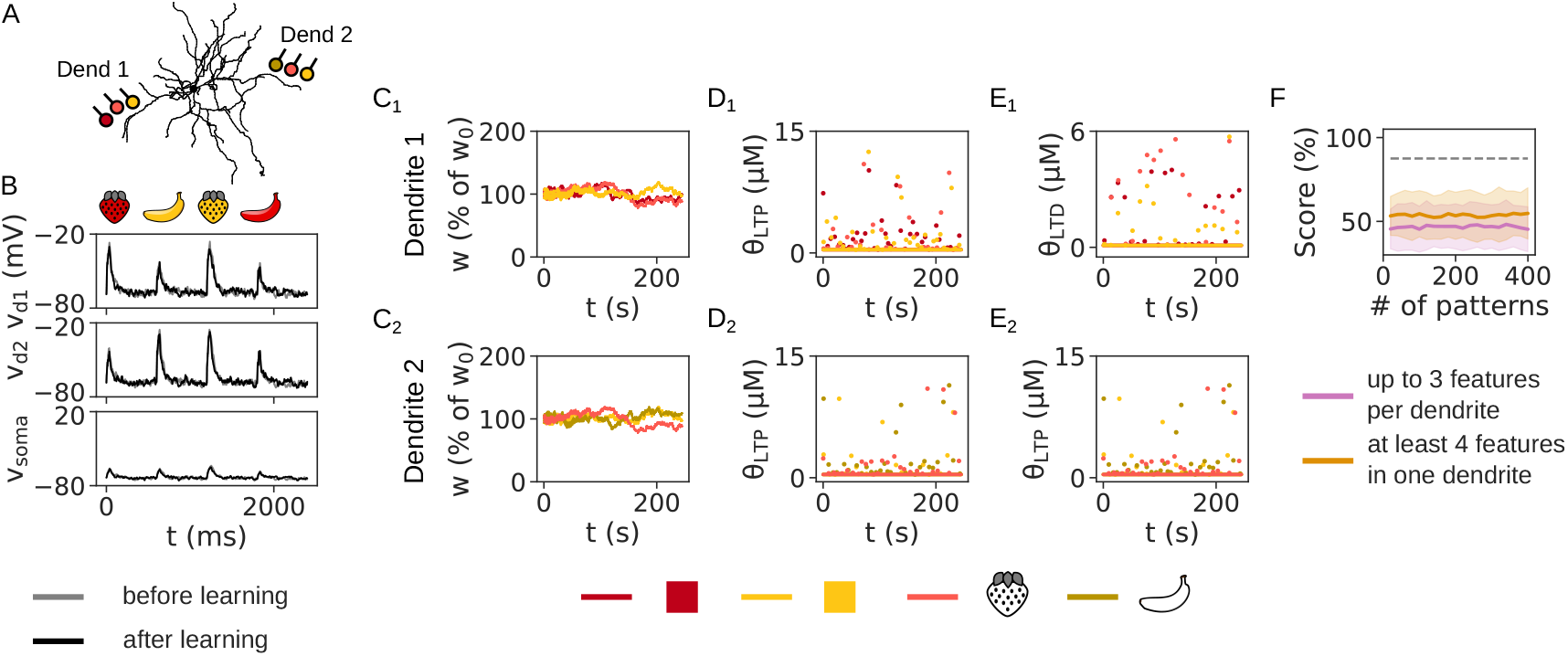
Learning outcome on the NFBP without metaplasticity (the LTP and the LTD thresholds are fixed). (A) An illustration of the setup for the NFBP (same as in Fig. 3B_3_). (B) The dendritic and somatic voltage before (gray traces) and after learning (black traces) for all four patterns. (C) The evolution of synaptic weights in both dendrites. (D, E) The fixed LTP thresholds (D) and LTD thresholds (E) in each dendrite, with calcium amplitudes during pattern presentation shown with dots (omitting some patterns for clarity). (F) The performance on the NFBP without metaplasticity (same number of trials as in Fig. 7F. **Figure 8—figure supplement 1**. The effect of having no metaplasticity on solving the FBP.

For the FBP, the results in the previous two sections again give an intuition for how partial metaplasticity would work. As shown in Figs. 6A_1_–E_1_ and 6A_2_–E_2_, even when the LTP threshold is fixed, it does not have an effect on solving the FBP (because there are no shared features in the FBP which ultimately need to be weakened). Similarly, neither does it have an effect when it is partially active: the LTP threshold is merely less elevated than compared to when metaplasticity is always active (Figure 7–figure supplement 1C, cf. Figs. 4C_1_, C_2_). On the other hand, partial metaplasticity in the LTD threshold is not enough to protect the shared strengthened synapses from weakening. The LTD threshold does not follow [Ca]_L-type_ fast enough (Figure 7–figure supplement 1D), and as a result the shared “strawberry” synapses are not strengthened properly (even though, for example, their weights stabilize when using supralinear integration, Figure 7–figure supplement 1B). Similarly to Figs. 5A_1_–E_1_ and 5A_2_–E_2_, this does not affect learning to solve the FBP with linear integration (Figure 7–figure supplement 1A_1_ and 1E_1_), but it does affect learning of the FBP with supralinear integration (Figure 7–figure supplement 1A_2_ and 1E_2_).

The effects of having no metaplaticity on the FBP with linear integration are similar to those of having partial metaplasticity (Figure 8–figure supplement 1). Again, the synapses encoding the shared feature cannot stabilize (synapses for “strawberry”, Figure 8–figure supplement 1B), only moderately affecting the performance (Figure 8–figure supplement 1A_1_ and 1E_1_). No metaplasticity in the FBP with supralinear integration also causes synapses encoding the shared to not stabilize (Figure 8–figure supplement 1A_2_). However, compared to partial metaplasticity, the unstable shared synapses sometimes evoke plateaus as they oscillate, on average solving the task (Figure 8–figure supplement 1E_2_). In summary, this shows that in the FBP, metaplasticity is important for robust, consistently high performance on the task.

### Reversal learning can be solved when relaxing the conditions for metaplasticity or introducing a signal that resets the thresholds

We have so far demonstrated a situation where the calcium thresholds are initially set to a low value, making the synapses “flexible” for learning in the time it takes for the thresholds to reach the [Ca]_NMDA_ and [Ca]_L-type_ amplitudes. Once the thresholds have elevated and reach the calcium amplitudes, the synapses are effectively “locked”, providing stable information storage. Conversely, fixing the calcium thresholds to study metaplasticity (in Figs. 5 and 6) made the synapses permanently flexible for learning and no information could be stored. This illustrates the role of metaplasticity as a lock on the plasticity process, and that it plays a role in solving the plasticity–stability dilemma, which means that synapses should “both be stable enough to store information and be plastic enough to accommodate new learning” (***Jedlicka et al., 2022)***. Reversal learning is a straightforward example of the plasticity–stability dilemma applied to the FBP and NFBP: after the tasks have been learned, we switch the reward policy so that relevant patterns become irrelevant (and are followed by dopamine pauses), and vice versa, irrelevant patterns become relevant (and are followed by dopamine peaks). To solve this task in the (N)FBP, the neuron should keep the learned changes in (one of the) the shared strengthened feature(s) (e.g. the shared “strawberry” is necessary for both the “red strawberry” and “yellow strawberry” patterns), and change the weights of the other features (e.g. “red” should be weakened and “yellow” should be strenghtened to reverse the neuron’s response from “red strawberry” to “yellow strawberry”).

We first show that “thresholded” metaplasticity cannot solve reversal learning (Fig. 9). The reward policy is reversed in the middle of the simulation, and all three tasks are solved before the reversal (blue voltage traces in Fig. 9A, and score in Fig. 9E before reversal). After reversal, because metaplasticity is only activated if a calcium signal is above its threshold (and followed by dopamine feedback), this means that the strengthened features cannot be weakened properly once their calcium levels fall below their LTD thresholds (strengthened synapses in Figs. 9B_1_–B_4_ weaken only slightly because their [Ca]_L-type_ levels fall below the LTD thresholds after weakening, shown in Figs. 9D_1_–D_4_; this fall in [Ca]_L-type_ is very prominent with supralinear integration because after weakening, no plateaus can be evoked). Neither can the weakened features be strengthened in most of the cases, because the previosuly irrelevant patterns, which are now rewarded and relevant, rarely cause [Ca]_NMDA_ above the LTP thresholds (calcium dots in Figs. 9C_1_–C_4_ largely stay below the LTP threshold of the weakened synapses). The FBP and NFBP are successfully learned before the point of reward policy reversal, while after reversal, the synapses cannot adapt to the new situation. As a result, in the end there is no spiking for the initially irrelevant patterns, nor is there spiking for the initially relevant patterns that were learned before reversal (Figs. 9A_1_–A_3_, black traces, and Figs. 9E_1_–E_3_). As expected, “thresholded” metaplasticity permanently locks the patterns in the dendrites, making the rule inflexible to learn new pattern-outcome associations if the reward policy changes.

**Figure 9.**
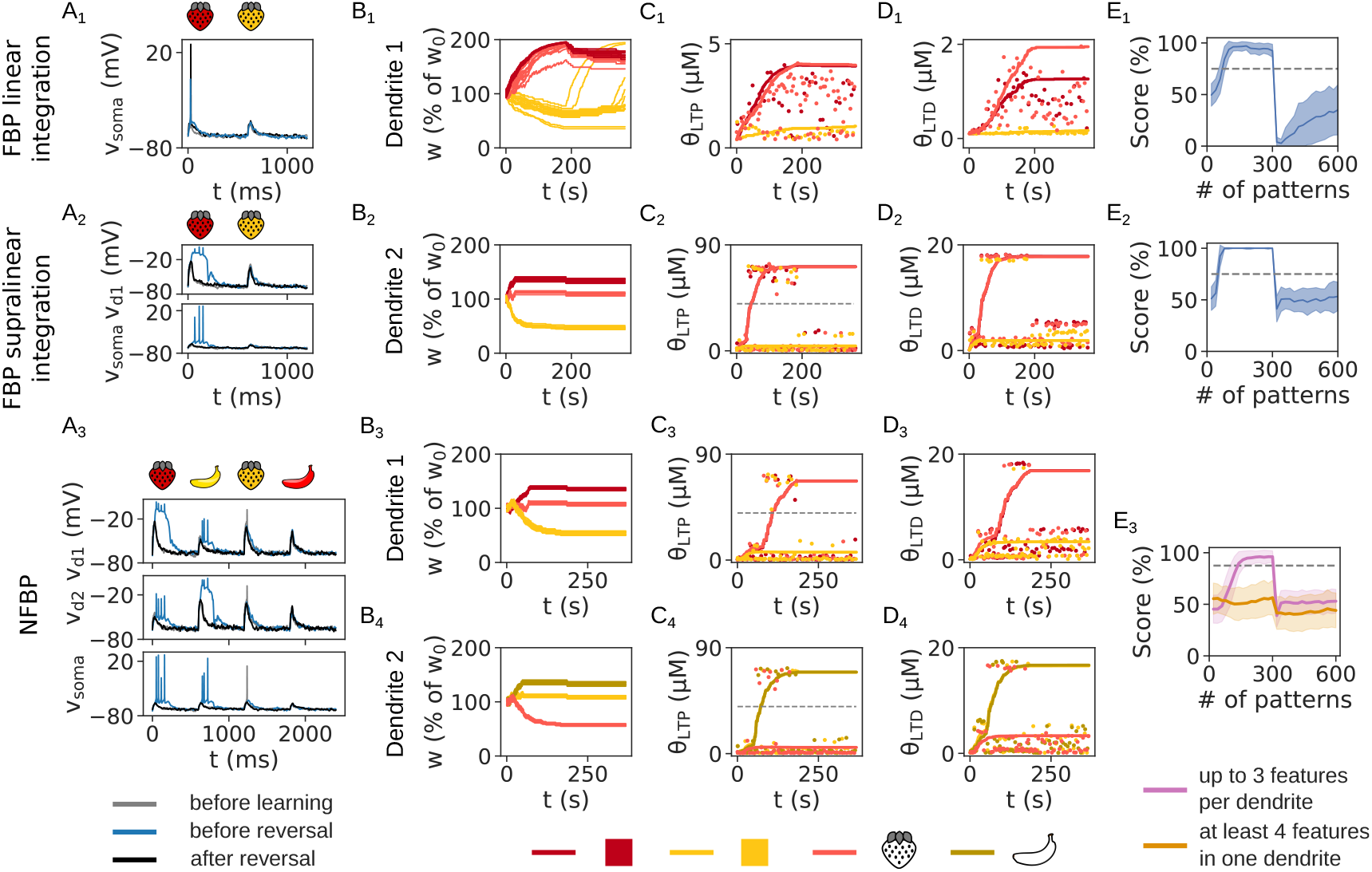
Reversal learning in the FBP and NFBP cannot be solved with “thresholded” metaplasticity. (A) The dendritic and somatic voltage before learning (gray traces), before reversal (blue traces) and after reversal (black traces) for the FBP with linear (A_1_) and supralinear integration (A_2_) and the NFBP (A_3_). (B–D) The evolution of synaptic weights (B), LTP thresholds (C) and LTD thresholds (D) for the synapses in each dendrite. In the middle of the simulation, the reward policy is reversed. Dashed lines in (C) show the upper LTP threshold, Θ_LTP_. In (B) all synapses are shown, and in (C, D) only one synapse per feature is used to show its calcium threshold. Dots represent the amplitudes of [Ca]_NMDA_ (C) and [Ca]_L-type_ (D) during pattern presentation, omitting some patterns for clarity. Dashed lines in (C_2_–C_4_) show the upper threshold for LTP, Θ_LTP_. (E) The performance on reversal learning in the FBP with linear (E_1_) and supralinear integration (E_2_) and the NFBP (E_3_). Performance on the FBP is averaged over 50 trials, and in the NFBP over 12 trials per input configuration (divided into the two groups shown in Figure 3–figure supplement 1), with 216 and 156 trials, respectively). **Figure 9—figure supplement 1**. “Relaxed” metaplasticity also solves the FBP and NFBP. **Figure 9—figure supplement 2**. “Relaxed” metaplasticity can overcome initially high calcium thresholds, unlocking synapses and allowing learning.

To solve reversal learning, we demonstrate two different approaches. One is to allow the thresholds to be lowered again, i.e. prevent them from stabilizing at an elevated level forever. To do this, we simply relax the conditions for metaplasticity so that any calcium levels in the active synapses can cause metaplasticity, i.e. calcium no longer needs to be above a threshold level for metaplasticity to occur (note that those conditions still hold for synaptic plasticity, as in Eq. 1). This approach has been taken in an extension of the cascade model, which in its original form cannot solve reversal learning. (In the generalized model the threshold is allowed to move in both directions (***Farashahi et al., 2017)***.) We first show that such “relaxed” metaplasticity can solve the FBP and NFBP (Figure 9–figure supplement 1). In this case, the thresholds always follow any calcium levels evoked when a synapse is activated by presynaptic input, allowing them to lower in value. When the variability in the calcium amplitudes is small, the fluctuations in the thresholds are also small, as occurs in the FBP with linear integration (Figure 9–figure supplement 1C_1_, D_1_), the strengthened “red” synaptic cluster in the FBP with supralinear integration (Figure 9–figure supplement 1C_2_, D_2_, dark red traces) and the weakened synaptic clusters in supralinear integration (Figure 9–figure supplement 1C_3, 4_, D_3,4_). Conversely, when the variability in the calcium amplitudes is large, as occurs in the shared synapses that need to be strengthened when using supralinear integration, the thresholds show large fluctuations (Figure 9–figure supplement 1B_2–4_–D_2–4_). Strictly speaking, with “relaxed” metaplasticity the thresholds never stabilize, which is particularly evident for the shared strengthened clustered synapses with large variability in the calcium amplitudes (Figure 9–figure supplement 1C_2–4_, D_2–4_). Despite the fluctuations, the thresholds perform their function equally well: the LTD thresholds are elevated enough to protect the strengthened synapses from weakening, and the LTP thresholds follow [Ca]_NMDA_ in the the weakened synapses to prevent them from strengthening. Note that because of the relaxed metaplasticity conditions, the LTD thresholds for the weakened synapses are not elevated as with “thresholded” metaplasticity (Figure 9–figure supplement 1D_2_–D_4_, cf. Fig. 5D_2_–D_4_). Hence, the weakened synapses decrease almost to the minimal level *w*_min_ (Figure 9–figure supplement 1B_2_–B_4_). Finally, “relaxed” metaplasticity allows to overcome initially high calcium thresholds set before learning (Figure 9–figure supplement 2). With “thresholded” metaplasticity, if the thresholds are initialized to values higher than the calcium signals, the synapses will remain locked and no learning will occur (Figure 9–figure supplement 2B_1_, B_2_). “Relaxed” metaplasticity can instead adapt the thresholds, thus unlocking the synapses and allowing learning (Figure 9–figure supplement 2B_3_, B_4_).

Relaxing the conditions for metaplasticity allows the rule to solve reversal learning in both the FBP and the NFBP (Fig. 10). The key to solving it with this approach is that calcium thresholds which have been raised while learning the FBP and NFBP are free to be lowered again after reversing the reward policy. In the FBP, after reversal, the LTD thresholds for the features “red” and “strawberry” are lowered, so these two features are weakened (Figs. 10B_1_, B_2_ and 10D_1_, D_2_, these weights and thresholds are weakened immediately after reversal). Simultaneously, “yellow strawberry” evokes higher [Ca]_NMDA_ than the “yellow” LTP threshold because in the initial stages of reversal the feature “strawberry” is still strong, despite being weakened (Figs. 10C_1_, C_2_, LTP threshold for “yellow” rises to follow the evoked [Ca]_NMDA_). When the feature “yellow” is strengthened enough, it elicits strong [Ca]_NMDA_ together with the “strawberry” feature, causing the latter to strengthen again (Figs. 10B_1_, B_2_, “yellow” synapses increase after reversal, and “strawberry” synapses follow after some time). The LTD thresholds for “yellow” and “strawberry” become elevated and protect the two strengthened features from weakening (Figs. 10D_1_–D_3_). Conversely, the LTP thresholds for “red” follow [Ca]_NMDA_ and prevent this feature from strengthening. In this way, in the FBP, the initially relevant pattern “red strawberry” stops evoking spiking (through linear integration at the soma or via a plateau), and the neuron starts responding to the initially irrelevant pattern “yellow strawberry”, thus solving the task (blue voltage traces before reversal learning and black traces after reversal learning in Fig. 10A_1_, A_2_, and performance scores in Fig. 10E_1_, E_2_).

**Figure 10.**
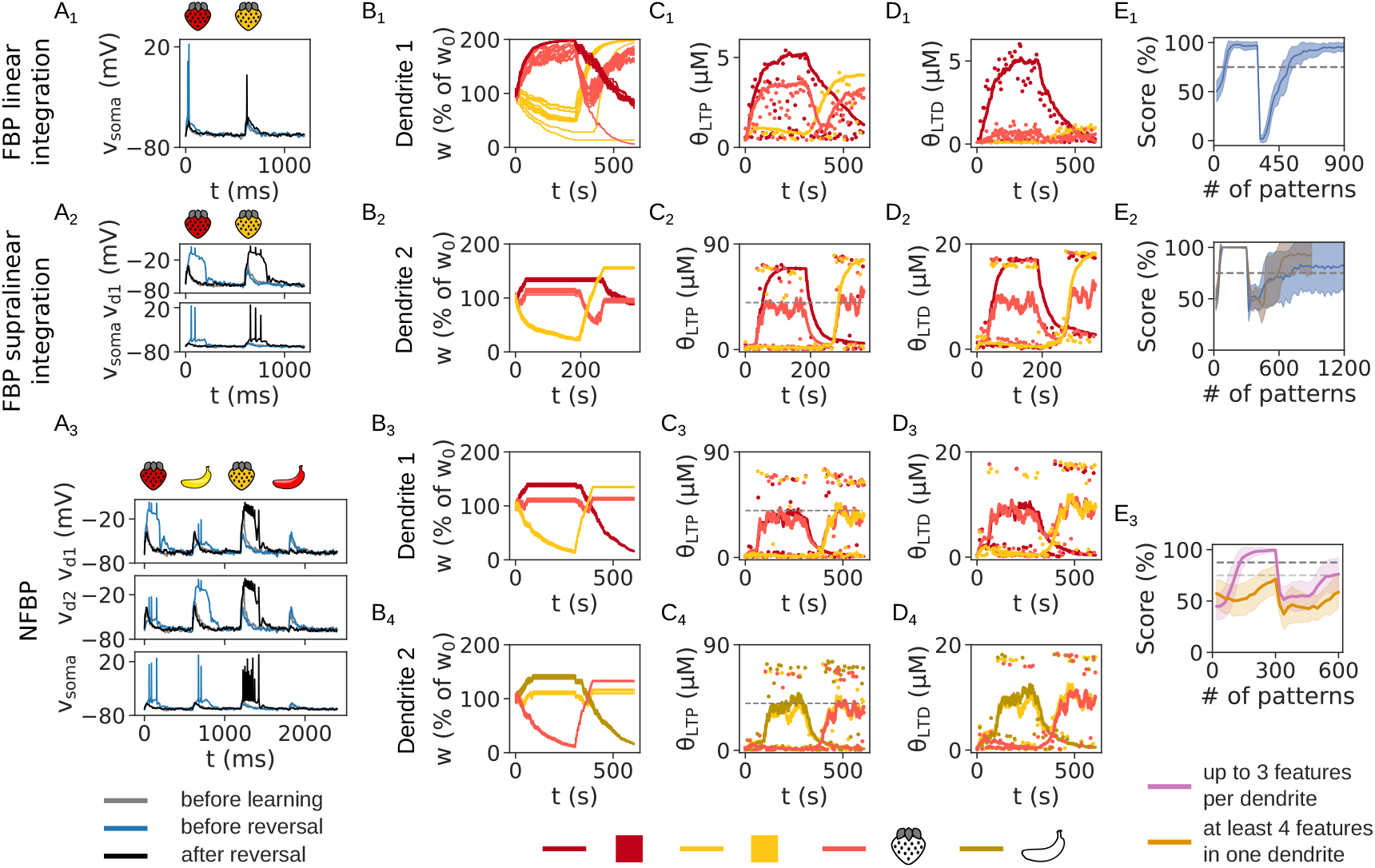
Reversal learning in the FBP and NFBP is solved when the conditions for metaplasticity are relaxed. (A) The dendritic and somatic voltage before learning (gray traces), before reversal (blue traces) and after reversal (black traces) for the FBP with linear (A_1_) and supralinear integration (A_2_) and the NFBP (A_3_). (B–D) The evolution of synaptic weights (B), LTP thresholds (C) and LTD thresholds (D) for the synapses in each dendrite. In the middle of the simulation, the reward policy is reversed. Dashed lines in (C) show the upper LTP threshold, Θ_LTP_. In (B) all synapses are shown, and in (C, D) only one synapse per feature is used to show its calcium threshold. Dots represent the amplitudes of [Ca]_NMDA_ (C) and [Ca]_L-type_ (D) during pattern presentation, omitting some patterns for clarity. (E) The performance on reversal learning in the FBP with linear (E_1_) and supralinear integration (E_2_) and the NFBP (E_3_). Performance on the FBP is averaged over 50 trials. In (E_2_), due to the thresholds changing slowly, it often occurs that after reward policy reversal, calcium never goes above the thresholds to trigger new learning, thus lowering the performance (blue trace). Increasing the metaplasticity rate to *η*_*θ*_ = 4 can solve this (brown trace). For the NFBP, 12 trials per input configuration are used (divided into the two groups shown in Figure 3–figure supplement 1, with 216 and 156 trials, respectively). **Figure 10—figure supplement 1**. All input configurations in reversal learning with the NFBP that can store one initially relevant pattern and both irrelevant patterns. **Figure 10—figure supplement 2**. Relaxed metaplasticity also solves the NFBP in input configurations that can store both (initially) irrelevant patterns. **Figure 10—figure supplement 3**. Reversal learning in the NFBP on one input configuration that can store both relevant and both irrelevant patterns.

The same occurs in the NFBP – the neuron stops responding to the two initially relevant patterns (Fig. 10A_3_, blue voltage traces before reversal show a plateau potential in each dendrite evoked by a relevant pattern, and black traces after reversal show no plateaus and no somatic spiking for these two patterns). However, since there is no structural plasticity, the available synaptic connectivity in this example (Fig. 3A_3_) allows for only one of the initially relevant patterns to be stored (in both dendrites, as seen from the synaptic weights in Figs. 10B_3_, B_4_; also, black voltage traces after reversal show plateaus in both dendrites for “yellow strawberry” in Fig. 10A_3_). This is also reflected in the performance, which shows that before reversing the reward policy, the NFBP is solved, while after the reversal the neuron responds to only one pattern (Figs. 10E_3_). To show that both initially irrelevant patterns can be learned in reversal learning, we also simulated the task on input configurations that are innervated by only one initially relevant pattern, but both irrelevant patterns (Figure 10–figure supplements 1 and 2). An example of such a configuration is given in Figure 10– figure supplement 2A, where only one initially relevant pattern (“red strawberry”) can be stored in both dendrites, but both initially irrelevant patterns (“yellow strawberry” and “red banana”) innervate dendrites 1 and 2, respectively. During learning, the neuron first learns to respond to “red strawberry”, and after reward policy reversal spikes to both “yellow strawberry” and “red banana”, elicited via plateau potentials in dendrites 1 and 2, respectively (Figure 10–figure supplement 2B, blue traces before reversal, and black traces after reversal, and Figure 10–figure supplement 2E, score of 75% before reversal, and close to 100% after reversal). Finally, there are a total of four input configurations in group 1 in Figure 3–figure supplement 1 and Figure 10–figure supplement 2 (marked with a star) which can store both relevant patterns and both irrelevant patterns. Reversal learning using (one of) these is shown in Figure 10–figure supplement 3, demonstrating that given the appropriate structural connectivity, reversal learning from both relevant to both irrelevant patterns can be done.

Another approach to solve reversal learning is to assume that another part of the brain detects the change in the reward policy and sends a signal to temporarily “reset” the permanently stabilized thresholds to a low value, thus “unlocking” the synapses, i.e. making the weights flexible again to relearn the new pattern-reward associations (Fig. 11). (This approach has also been taken in another extension of the original cascade model (***Iigaya, 2016)***.) The noradrenergic system, particularly the locus coeruleus, has been implicated in detecting unexpected changes in environmental conditions and facilitating behavioral adaptation, including during reversal learning (***Jordan, 2024; Sara, 2009; Pajkossy et al., 2023)***. The striatum receives only limited projections from the locus coeruleus, but SPNs express adrenergic receptors which modulate phosphorylation of DARPP-32, an important protein regulating cortico-striatal synaptic plasticity (***Hara et al., 2010)***, and hence could be affected by such signals about environmental changes.

**Figure 11.**
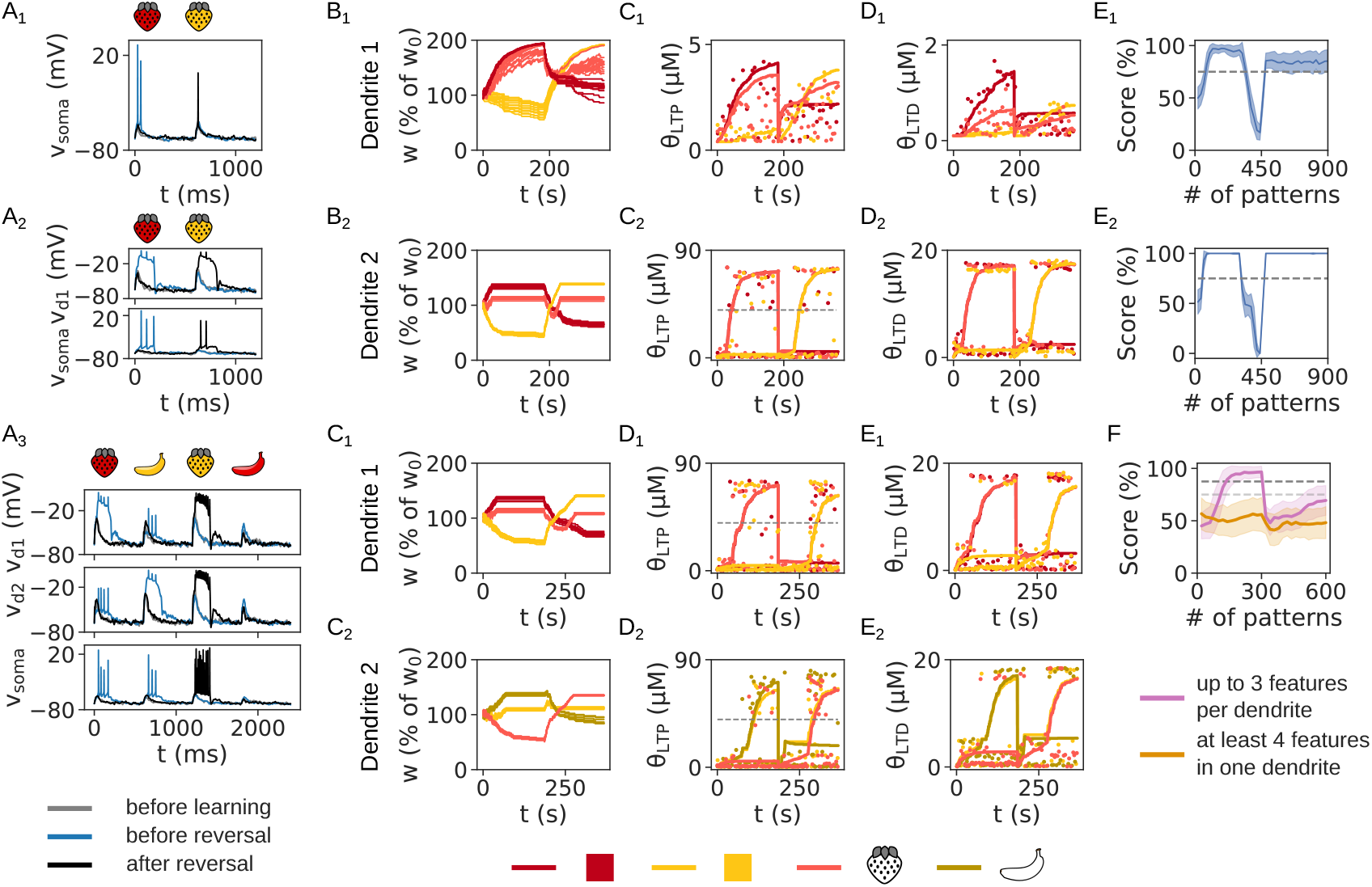
Threshold resetting solves reversal learning. A signal which resets the thresholds is assumed to arrive from a different brain region detecting the change in reward policy. (A) The dendritic and somatic voltage before learning (gray traces), before reversal (blue traces) and after reversal (black traces) for the FBP with linear (A_1_) and supralinear integration (A_2_) and the NFBP (A_3_). (B–D) The evolution of synaptic weights (B), LTP thresholds (C) and LTD thresholds (D) for the synapses in each dendrite. In the middle of the simulation, the reward policy is reversed. Dashed lines in (C) show the upper LTP threshold, Θ_LTP_. In (B) all synapses are shown, and in (C, D) only one synapse per feature is used to show its calcium threshold. Dots represent the amplitudes of [Ca]_NMDA_ (C) and [Ca]_L-type_ (D) during pattern presentation, omitting some patterns for clarity. (E) The performance on reversal learning in the FBP with linear (E_1_) and supralinear integration (E_2_) and the NFBP (E_3_). Performance on the FBP is averaged over 50 trials, and in the NFBP over 12 trials per input configuration (divided into the two groups shown in Figure 3–figure supplement 1, with 216 and 156 trials, respectively).

### Two calcium thresholds provide separate control over LTP and LTD

In this section we ask the question what is the use of two separate calcium thresholds for LTP and LTD? The FBP, NFBP and reversal learning can be solved just as well with a single plasticity threshold (Fig. 12 and Figure 12–figure supplements 1 and 2). To show this we simulated a scenario where calcium from both NMDA and L-type channels is tracked with a single variable, [Ca] (rather than two separate ones), and diffuses axially across the spines and dendrites, and the same [Ca] signal is responsible for LTP and LTD when it crosses a single calcium threshold (Eqs. 4 and 5). This is not biologically realistic in dSPNs, but allows us to compare the functionality of two thresholds versus just one. As before, the dopamine signal determines the direction of synaptic plasticity:

**Figure 12.**
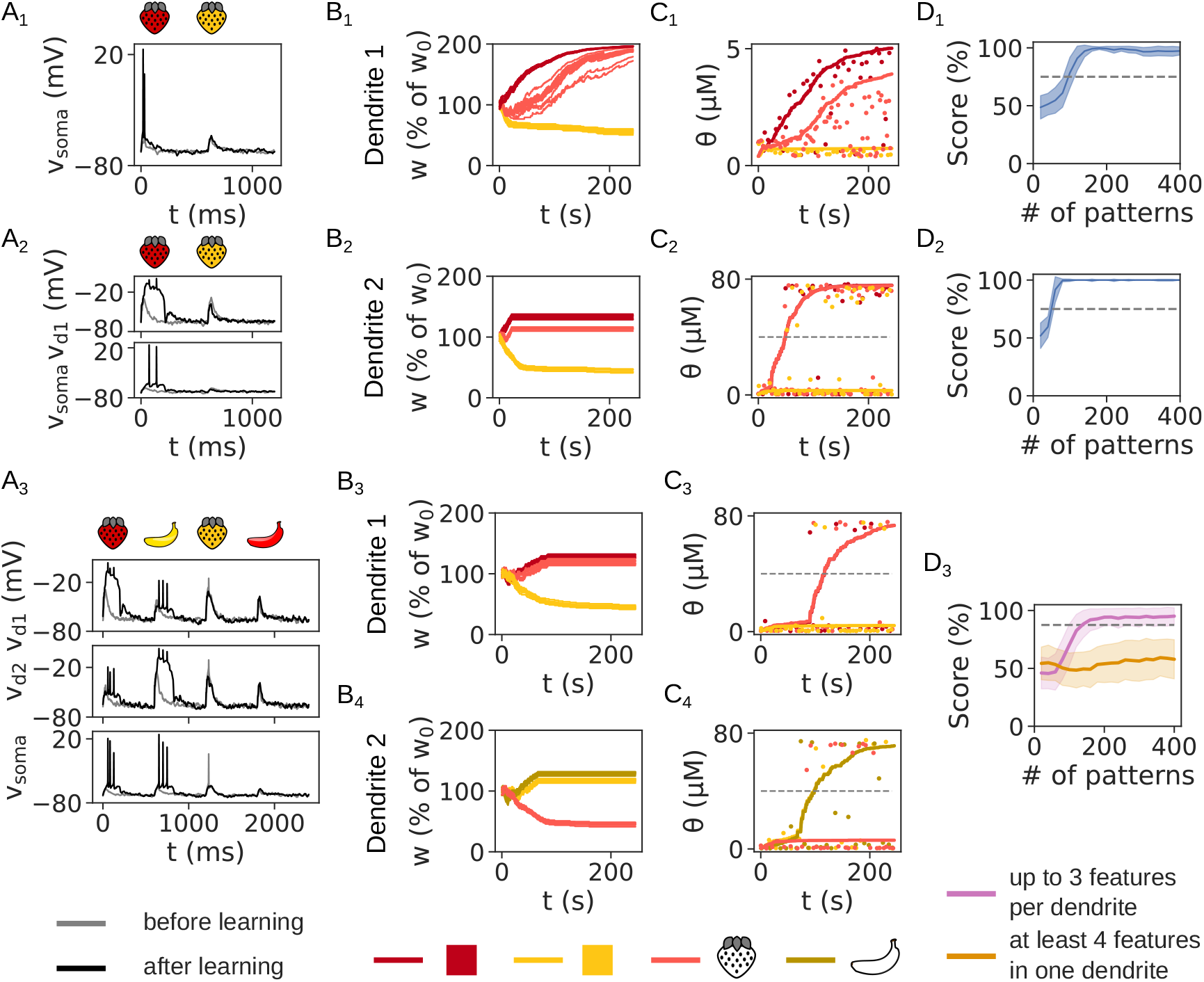
A single plasticity threshold is sufficient to solve the FBP and NFBP. “Thresholded” metaplasticity is used here. (A) The dendritic and somatic voltage before (gray traces) and after learning (black traces) for the FBP with linear (A_1_) and supralinear integration (A_2_) and the NFBP (A_3_). (B, C) The evolution of synaptic weights (B) and the single calcium threshold (C) for the synapses in each dendrite. Dashed lines in (C) show the upper LTP threshold, Θ_LTP_. In (B) all synapses are shown, and in (C) only one synapse per feature is used to show its calcium threshold. Dots represent the [Ca] amplitudes during pattern presentation, omitting some patterns for clarity. Dashed lines in (C_2_–C_4_) show the upper threshold for LTP, Θ_LTP_. (D) The performance on the FBP with linear (D_1_) and supralinear integration (D_2_) and the NFBP (D_3_). Performance on the FBP is averaged over 50 trials, and in the NFBP over 12 trials per input configuration (divided into the two groups shown in Figure 3–figure supplement 1, with 216 and 156 trials, respectively). **Figure 12—figure supplement 1**. A single plasticity threshold is sufficient to solve the FBP and NFBP. when the conditions for metaplasticity are relaxed. **Figure 12—figure supplement 2**. A single plasticity threshold is sufficient to solve reversal learning when the conditions for metaplasticity are relaxed. **Figure 12—figure supplement 3**. Learning outcome without metaplasticity in the single threshold scenario (the threshold is fixed). **Figure 12—figure supplement 4**. [Ca]_NMDA_ and [Ca]_L-type_ are highly correlated and carry essentially the same information.

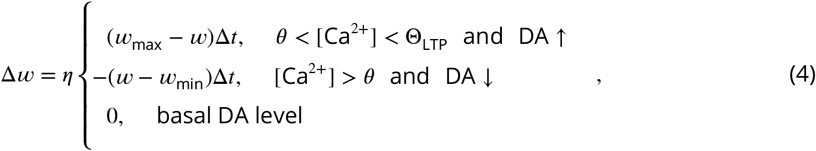

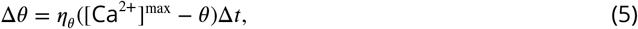

and we use both “thresholded” metaplasticity (*θ* is updated only when [Ca] goes above it, Fig. 12) and the relaxed condition for metaplasticity (*θ* is updated for any [Ca] level in active synapses, Figure 12–figure supplement 1). In both cases, the dynamics of the weights and thresholds are analogous to those when two separate calcium thresholds are used. As before, no metaplasticity (fixing the threshold to a low value) results in the synapses representing shared features not stabilizing (Figure 12–figure supplement 3). With one calcium threshold, in the case of the NFBP none of the synapses stabilize (as occurs when both thresholds are fixed in Fig. 8). Comparing Figure 12–figure supplement 3B_3_, B_4_ to Figs. 5B_3_, B_4_ and 6B_3_, B_4_ where just one of the two calcium thresholds is fixed, we see that having two separate thresholds allows for separate control over synaptic strengthening and weakening.

The reason why a single plasticity threshold works as well as two separate ones on the FBP, NFBP and reversal learning is that [Ca]_NMDA_ and [Ca]_L-type_ are driven by the same variable synaptic voltage. Both their amplitudes, which are used to compare against the thresholds and the full [Ca]_NMDA_ and [Ca]_L-type_ time-courses are highly correlated (Figure 12 – figure supplement 4), meaning that the two signals carry the same information, and it is not necessary to have two separate calcium variables to track that information. In this study, we have triggered metaplasticity in both thresholds using the same signal (a calcium source should be above its threshold, followed by a dopamine signal; or simply a dopamine signal with “relaxed” metaplasticity). If metaplasticity in the LTP and LTD thresholds is triggered under different conditions for each, then two separate thresholds could be useful.

## Discussion

In this article we studied the effects of metaplasticity in two separate calcium thresholds for inducing synaptic plasticity, one of which is a threshold for LTP, and the other for LTD. Plasticity occurs when calcium levels from different sources, implicated in synaptic plasticity in SPNs, reach levels above the thresholds. We used the FBP and NFBP because the patterns in these tasks share their features, and the role of each threshold is clearly demonstrated on the synapses representing these shared features. Metaplasticity slides the thresholds towards the calcium amplitudes evoked by arriving patterns. Two different metaplasticity rules were tested: i) one where calcium levels need to be above the thresholds to trigger metaplasticity (“thresholded” metaplasticity), and ii) one where any calcium levels trigger metaplasticity (relaxed conditions). The results showed that:

1. The FBP and NFBP can be solved with this simple plasticity and metaplasticity rule, which is required in order to demonstrate the role of each calcium threshold. With this rule, metaplasticity is necessary for solving the NFBP, whereas in the FBP it ensures robust and consistently high task performance.
2. Metaplasticity enables synapses undergoing both LTP and LTD during learning to selectively express (converge to) only one form of plasticity (either LTP or LTD).
3. Metaplasticity in the LTD threshold protects learned patterns from being weakened, allowing the LTP process to properly strengthen (encode) the patterns. In effect, *the threshold for weakening allows proper synaptic strengthening*.
4. Metaplasticity in the LTP threshold prevents unnecessary features from being strengthened, allowing the LTD process to persistently weaken them. In effect, *the threshold for strengthening allows proper synaptic weakening*.
5. For successful learning of the NFBP, metaplasticity in *both* thresholds needs to occur during a synaptic modification, irrespective of whether the modification is LTP or LTD.
6. Relaxing the conditions for metaplasticity so that it is triggered by any calcium levels allows reversal learning in the FBP and NFBP to be solved, demonstrating metaplasticity’s importance for solving the plasticity–stability dilemma. Alternatively, assuming a signal exists that can reset the calcium thresholds, reversal learning can be solved without relaxing the conditions for metaplasticity.
7. Using a single calcium threshold instead of two separate ones also solves the FBP, NFBP and reversal learning, but does not allow for separate control over synaptic strengthening and weakening.

### The role of metaplastcity in this and other studies

In the original BCM rule, metaplasticity prevents runaway growth or collapse of the synaptic weights, which is one of its most commonly cited roles (***Bienenstock et al., 1982; Abraham, 2008; Hulme et al., 2013)***. This is achieved by using a nonlinear function in the update rule for the threshold. Conversely, here we used a linear function in the metaplasticity rule for sliding the thresholds towards the calcium amplitudes, so metaplasticity by itself cannot prevent the weights from unbounded growth (unless that growth produces saturation in the evoked calcium amplitude). Instead, metaplasticity has a different role in this study – it enables synapses exposed to competing LTP and LTD drives to converge to one stable plasticity state, i.e. either LTP or LTD – without metaplasticity, the synapses never stabilize. Although not explicitly demonstrated, the same role of metaplasticity can be inferred from the dynamics of the cascade model (***Fusi et al., 2005; Farashahi et al., 2017)***. In fact, the calcium thresholds in the current rule are a possible mechanistic implementation of the abstract hidden metaplastic states in the cascade model: threshold sliding is analogous to moving up and down the cascade of states, making synaptic plasticity easier or more difficult to occur. (Hence the similar role of metaplasticity in both studies.)

More specifically, this study shows that the two calcium thresholds have opposite and complementary roles in regulating the strengthening and weakening of synapses. The ultimate effects of metaplasticity are unexpected, at least at first glance, since the threshold for one plasticity type (e.g. LTD) allows proper expression of the other plasticity type (e.g. LTP). In addition to separately controlling the two plasticity processes, two thresholds allow implementing a different update rule for each threshold (a case which we have not studied here). Generally speaking, two thresholds would be useful if the different calcium concentrations that they track are uncorrelated, i.e. exhibit separate dynamics, which does not seem to be the case for the NMDA and L-type channels in the dSPNs. They could also be useful if metaplasticity in each threshold is triggered by a different signal (a case which we also did not study here, but only touched upon with partial metaplasticity). Having two thresholds might also contribute to increased robustness of the machinery for synaptic plasticity across different conditions, a common feature of biological systems (***Kitano, 2004; Wollman, 2018)***.

Finally, in a range of studies using the same threshold for all synapses, derived from the neuron’s activity (similarly to ***Bienenstock et al. (1982)***), metaplasticity provides a homeostatic, heterosynaptic effect which helps to keep a neuron’s firing rate stable (***Jedlicka et al., 2015; Zenke et al., 2013)***. The learning rule in this article does not incorporate such effects. As a result, without an upper bound on the synaptic weight, synapses would increase indefinitely in the case of FBP with linear integration, causing a depolarization block in the neuron. Similarly, without the upper threshold for LTP, they would also increase indefinitely when using supralinear integration. However, in the latter case the plateau potential provides stability of the neuronal firing rate, since the dendritic voltage is limited by the NMDAR reversal potential.

### Directions for future studies

This study also provides a basis for exploring other metaplasticity rules. Here we only considered metaplasticity evoked in the same synapses activated by presynaptic spikes (homosynaptic metaplasticity). However, heterosynaptic metaplasticity mechanisms also exist, where the evoking stimuli originate from different synapses or different neurons (***Abraham, 2008; Abraham et al., 2019; Hulme et al., 2014)***. Furthermore, as detailed calcium models for other neurons are developed, the contributions of different calcium sources in triggering synaptic plasticity can be studied using this simple learning rule as a basis. Regarding SPNs specifically, more realistic metaplasticity rules can also be developed once future experimental findings appear.

Some examples of different metaplasticity mechanisms are known from other brain areas. In the Schaffer collateral synapses to the CA1 hippocampal area, prior activation of NMDARs (which does not induce long-term plasticity) temporarily increases the LTP threshold and prevents LTP, while prior activation of mGluRs facilitates LTP (***Huang et al., 1992; Cohen and Abraham, 1996)***. Additionally, weakening these synapses lowers the LTP threshold and raises the LTD threshold, while weakening does the opposite, i.e. lowers the LTD threshold and likely raises the LTP threshold (***Ngezahayo et al., 2000)***. In the synapses from the perforant path to the dentate gyrus, prior stimulation at theta frequency (typical for the hippocampus) lowers the threshold for both LTP and LTD (***Christie et al., 1995; Christie and Abraham, 1992)***. Additionally, prior stimulation at a high frequency also lowers the LTD threshold in both unchanged and in previously strengthened synapses (***Holland and Wagner, 1998)***. Adapting the learning rule to incorporate the metaplasticity mechanisms in different neurons will allow to study whether metaplasticity supports other specific functions in various brain regions.

In addition to signals arising from the local neuronal microcircuitry, it is interesting to see if global signals reaching many brain areas might also have a role in triggering metaplasticity. The noradrenergic neuromodulatory system is involved in detecting unexpected changes in learned pattern-outcome associations, and has been proposed to transiently interrupt or reset ongoing neural processing (***Jordan, 2024; Sara, 2009; Pajkossy et al., 2023; Yu and Dayan, 2005; Dayan and Yu, 2006)***. Noradrenaline is known to modulate synaptic plasticity through adrenergic receptors, such as the phosphorylation level of DARPP-32 (***Hara et al., 2010)***, an important protein for corticostriatal synaptic plasticity. Hence, it is interesting to ask whether noradrenergic signaling can act as a global metaplastic signal that transiently modifies plasticity thresholds.

An important direction for future work is to develop a learning rule that also models structural plasticity. Since NMDAR-dependent supralinear integration relies on synaptic clustering within a short dendritic segment, or more widely, within a whole branch (***Losonczy and Magee, 2006; Branco et al., 2010)***, a learning rule would ideally implement mechanisms that can give rise to such clusters. Synaptic clustering occurs already in development, and learning in the motor cortex, for example, builds on existing functional clusters by growing new spines into them (***Winnubst and Lohmann, 2012; Hedrick et al., 2022)***. Motor learning also strengthens the synaptic connections from the motor cortex to the striatum, which are themselves clustered (***Hwang et al., 2022)***. A rule incorporating structural plasticity could then also include thresholds specific to structural plasticity (***Ueda et al., 2022)***.

### Assumptions in the learning rule

We have made two assumptions when formulating the learning rule. The first is that metaplasticity operates in cortico-striatal synapses, which is not a strong assumption, since metaplasticity has been reported for many synapses. The second – a major assumption – is the upper threshold for synaptic strengthening, which means that high [Ca]_NMDA_ during LTP conditions (a dopamine peak) prevents synaptic plasticity (blocks LTP). It is an open question whether this assumption is true. Nevertheless, there are some molecular mechanisms which could form the basis for such an upper threshold. In the neuromodulatory branch of the signaling network in Fig. 1 (the branch activated by dopamine), the enzyme adenylyl cyclase 5, which transduces the dopamine signal from the D_1_R, is inhibited by calcium. This indicates that high [Ca]_NMDA_ might prevent synaptic strengthening already very early in the signaling network. Further downstream in the signaling network, calcium activates PP2B through calmodulin, and PP2B inactivates PKA, an enzyme necessary for synaptic strengthening (***Church et al., 2021; Yagishita et al., 2014)***. This is another route through which high [Ca]_NMDA_ could prevent LTP. Due to the complexity of the signaling network, the influence of high [Ca]_NMDA_ on synaptic plasticity requires further study, and kinetic models of the cortico-striatal synaptic signaling network which incorporate the many molecular interactions might provide some answers (***Lindskog et al., 2006; Nakano et al., 2010; Nair et al., 2015)***. Finally the astrocytically secreted S100B protein could be involved in limiting the amount of LTP within certain bounds (***Jones, 2015)***.

### Insights into nonlinear computation by neurons

Finally, this study gives insights into nonlinear computation by single neurons, which have a variety of mechanisms for nonlinear input processing, giving them computational abilities similar to those of multilayer artificial neural networks (***Gidon et al., 2020; Poirazi et al., 2003; Jones and Kording, 2021; Major et al., 2013; Antic et al., 2010; Tran-Van-Minh et al., 2015)***. Metaplasticity allows dSPNs to perform the NFBP, a computationally difficult task that relies on supralinear integration by the clustered synapses. In the simpler FBP, which can be solved with either linear or supralinear integration of synaptic inputs, metaplasticity ensures robust, consistently high performance on the task.

We have recently participated in another study with a different goal – whether SPNs can solve the NFBP (***Khodadadi et al., 2025)***. (The NFBP is related to the XOR problem, the canonical minimal example of a linearly non-separable task requiring nonlinear computation beyond a single-layer perceptron.) ***Khodadadi et al. (2025)*** uses a different learning rule that incorporates more biological complexity/detail and focused on dendritic integration which was much less supralinear, i.e. the dendrites were in a regime where they produced so-called “boosting” nonlinearities, that are likely common physiologically. Completely solving the NFBP required both excitatory and inhibitory synaptic plasticity, regardless of whether the synaptic clusters were closer or further away from the soma. Although here we study the role of metaplasticity, the results also add some information related to the solution of the NFBP. The all-or-none plateau potentials in this study achieve a very high score on the NFBP with excitatory plasticity alone. This indicates that with a threshold nonlinearity instead, the SPNs might be able to completely solve the task without requiring inhibitory synapses and even for very distally located synaptic clusters. Whether this is true will be explored in another study, as theoretical considerations suggest that the NFBP can be solved with excitatory synapses only (***Tran-Van-Minh et al., 2015)***.

In summary, at least when it comes to solving the NFBP, the proposed learning rule highlights the importance of deciding when to stop updating the synaptic weights. The two processes (mechanisms) that regulate when synaptic plasticity occurs are the upper plasticity threshold for LTP and metaplasticity. Without the upper plasticity threshold, all strengthened synapses would reach their maximally attainable synaptic weight. Without metaplasticity, the synaptic weights would not stabilize.

## Methods

### Plasticity rule

We highlight again that, similar to ***Khodadadi et al. (2025)***, the learning rule is:

- local: only signals at the local synaptic site are used to trigger plasticity,
- calcium- and reward-based: local calcium levels trigger plasticity, the type of which is decided by the dopamine reward signal, and
- always “on”: no separate training and testing phases are needed because learning is regulated by metaplasticity.

Differently from ***Khodadadi et al. (2025)***, the learning rule has metaplasticity in both calcium thresholds, and is also (much) simpler, making it suitable for studying the thresholds’ roles in synaptic plasticity.

The parameters used in the plasticity rule are given in Table 1. The values of *w*_max_ and *w*_min_ are not varied throughout the simulations. They describe the maximal and minimal values that the synaptic weight can take on in the model. Whether these values are reached depends on the metaplasticity mechanisms (the values of the calcium thresholds), and, in the case of supralinear integration, the upper plasticity threshold Θ_LTP_, which is a fixed parameter that can be set at any value in the region of the supralinearity (the jump) in [Ca]_NMDA_ in Fig. 2D_3_ (we used 40 *μ*M). The learning rate *η* and the metaplasticity rate *η*_*θ*_ are also kept fixed throughout all simulations, except in Figure 4–figure supplements 6–11, where they are varied to study their effect on learning. Finally, the LTP and LTD thresholds, *θ*_LTP_ and *θ*_LTD_, are initialized to low values, *θ*_LTP, 0_ and *θ*_LTD, 0_, which are also the minimal values that these thresholds can take on. They are initialized to low values so that synapses are flexible (“unlocked”) for learning. Otherwise, if they are set to high values, they may act to protect synapses from weakening (high *θ*_LTD, 0_) or prevent them from strengthening (high *θ*_LTP, 0_), as is shown with Figs. 5, 6 and Figure 9 – figure supplement 2. The synaptic weights are initialized uniformly at random from the interval [0.3, 0.35].

**Table 1.**
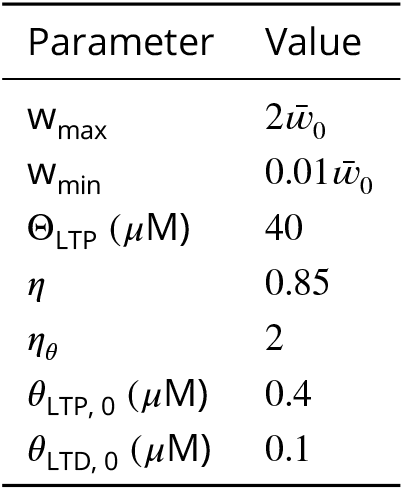
The parameters in the plasticity rule.

Fianlly, the rule uses the amplitudes of [Ca]_NMDA_ and [Ca]_L-type_ to trigger plasticity (and metaplasticity, in the scenarios with “thresholded” metaplasticity), implicitly assuming that such values can be remembered by the synapse’s molecular circuitry while dopamine feedback arrives. At least within the stimulation protocol in Fig. 3A, this assumption is justified, since dopamine feedback up to 2 seconds after synaptic stimulation can trigger LTP in experiments (***Yagishita et al., 2014)***.

### Neuron model

We use a morphologically realistic, multicompartmental model of a direct-pathway striatal projection neuron from the collection of 71 such models in ***Lindroos and Hellgren Kotaleski*** (***2020)*** (index 34 in the collection). As detailed in ***Lindroos and Hellgren Kotaleski*** (***2020)***, the ionic conductances in all models in the collection have been selected with a search process to produce frequency-current (F-I) responses and dendritic calcium elevations arising from backpropagating action potentials that match experimental data, and in addition produced a suitable number of spikes to injected current, no voltage oscillations leading up to the spikes, proper afterhyperpolarization behavior, and an F-I slope within three standard deviations of the experimentally determined mean.

The main contributor to supralinear integration in the SPN model is the activation of NMDARs. This has been shown, for example, in Fig. S7 in ***Du et al. (2017)***, where supralinear integration with clustered synapses is similar in an active and a passive dendrite in the SPN. This means that the NMDA synapses, and not the SPN’s ion channels, are causing it. SPN–specific ion channels do have a large effect on the SPN’s elecrophysiological behavior – the lowered resting potential of around -85 mV is a result of the strongly inwardly rectifying K^+^ channels (***Nisenbaum and Wilson***, ***1995; Wilson***, ***2005)***. Because of this, to raise their somatic voltage, the SPNs require strong synaptic drive from distributed synaptic inputs across the dendrites, or a localized activation of clustered synapses which generates a plateau potential. With this behavior, SPNs are said to have two states: a “down state”, which requires strong synaptic drive to reach the firing threshold, and an “up state”, where they are more sensitive to incoming synaptic inputs (***Plotkin et al., 2011)***. It has also been suggested that plateau potentials themselves cause the transition from a “down” state to an “up” state (***Oikonomou et al., 2014)***. Plateau potentials in SPNs normally do not cause somatic spiking without additional synaptic inputs, which is why in the simulations we have used elevated background noise.

In this study, synapses are placed on spines, which are not present in the original model in ***Lindroos and Hellgren Kotaleski*** (***2020)***. The spines consist of a neck and head with neck and head with lengths and diameters *l*_neck_ = 0.5*μ*m, *l*_head_ = 0.5*μ*m, and *d*_neck_ = 0.125*μ*m, *d*_head_ = 0.5*μ*m, respectively. Axial resistance in all compartments of the model is 150 Ω ⋅ cm, except for the spine neck, where it is 1130 Ω ⋅ cm (***Dorman et al., 2018)***. The spines contain the inwardly rectifying K^+^ channel, and the same voltage-gated calcium channels as in the dendrites. The spine calcium channel conductances have been manually tuned to match the relative proportions determined in ***Carter and Sabatini*** (***2004)*** and ***Higley and Sabatini*** (***2010)***, as well as the Ca^2+^ concentration amplitudes arising from backpropagating action potential (bAP) stimulation as in Fig. 2 of ***Shindou et al. (2011)***.

### Calcium dynamics and diffusion model

Because calcium from two different sources triggers LTP and LTD in SPNs, we tracked calcium from the two sources separately. Calcium from NMDA channels, when followed by a peak in dopamine, triggers LTP (***Shen et al., 2008; Yagishita et al., 2014; Fino et al., 2010; Fisher et al., 2017)***. Calcium from Ca_v_1.3 channels, when paired with a pause in dopamine, has been shown to be necessary for LTD (***Wang et al., 2006; Fino et al., 2010; Shindou et al., 2011; Iino et al., 2020)***.

Without implementing calcium diffusion explicitly, the accumulation of [Ca]_NMDA_ in a spine shows a strange effect when the synaptic cluster size is gradually increased, which is that the largest plateaus do not correspond to the largest calcium amplitude (Figs. 13A, B). This is simply a result of less NMDAR membrane current flowing in as the plateau’s voltage approaches the NMDAR reversal potential in Eq. 8. Larger plateaus giving rise to lower [Ca]_NMDA_ amplitudes could pose a problem to the learning rule, since the decision when to trigger plasticity and metaplasticity is made by comparing the [Ca]_NMDA_ amplitude against the LTP threshold. In a synaptic cluster that is being strengthened, but starts to evoke a smaller [Ca]_NMDA_ amplitude after some strengthening due to the effect in Figs. 13A, B, learning might be prematurely stopped as the [Ca]_NMDA_ amplitude falls below the increased LTP threshold.

**Figure 13.**
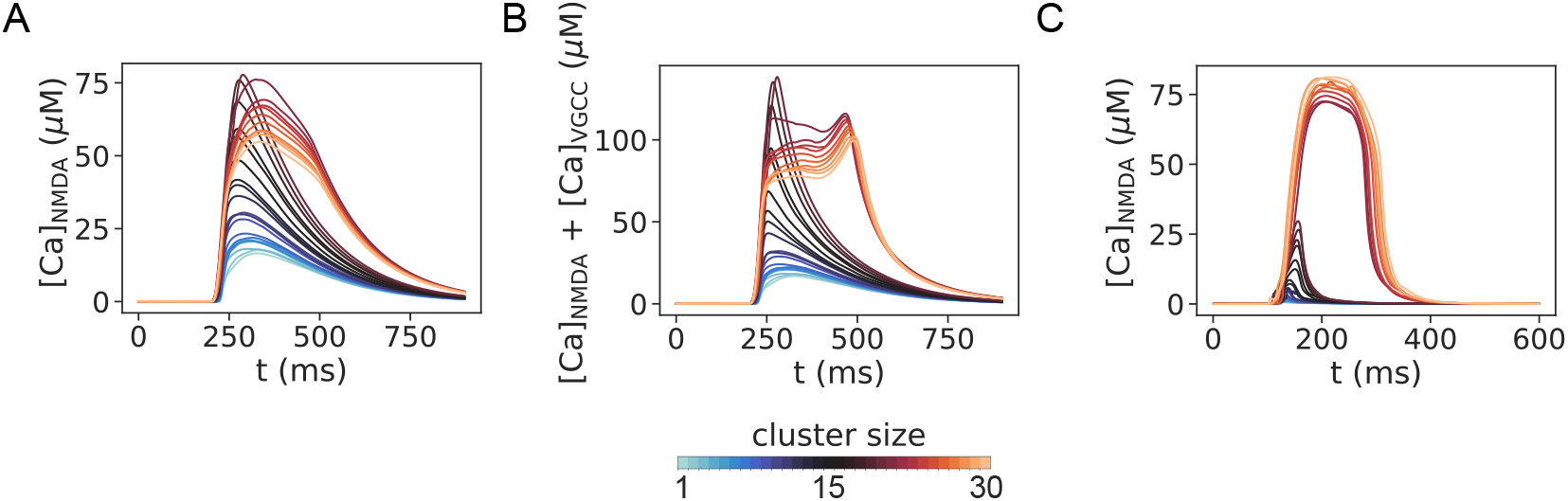
Alternative models of calcium accumulation. (A) Spine [Ca]_NMDA_ without axial diffusion. [Ca]_NMDA_ is described with a pool model similar to that in Eq. 6 (except that instead of [Ca]_L-type_, this pool accumulates [Ca]_NMDA_ arising from the NMDA current in Eq. 8). (B) Spine [Ca]_NMDA_ + [Ca]_VGCC_ without axial diffusion. Calcium from the NMDA current and all voltage-gated calcium channels (except L-type) accumulates in a pool model. (C) Spine [Ca]_NMDA_ with axial diffusion. Compared to Fig. 2C_3_, in this plot only calcium from NMDARs diffuses in the neuron, without calcium from other voltage-gated calcium channels. **Figure 13—figure supplement 1**. Axial diffusion of calcium from NMDARs only.

Because of this, we sought to obtain a monotonic increase of calcium amplitude with increasing plateau voltage (as in Fig. 2C_3_), which seems to also occur in experiments where plateaus are evoked (Fig. 2C_2_, D_2_ in ***Oikonomou et al. (2012)***). Increasing cluster size when evoking NMDA spikes also produces monotonic increases in calcium amplitude, at least for the voltages achieved by NMDA spikes (Figs. 5 and 8 in ***Losonczy and Magee*** (***2006)***). To obtain such monotonic increase, it was necessary to implement axial diffusion and buffering for [Ca]_NMDA_. Because tuning a detailed calcium diffusion model to data is a project in its own right, we used the most detailed calcium diffusion model for SPNs available to date (by ***Dorman et al. (2018)***) as a reference. We reuse the same buffer molecules, their amounts and their calcium binding and unbinding rates from ***Dorman et al. (2018)*** with small changes, given in Table 2. The rationale for reusing them is that the idealized morphology used in ***Dorman et al. (2018)*** should on average represent any other SPN, so the parameters obtained from the optimization procedure in that study should on average also apply to other multicompartment SPN models. The model in ***Dorman et al. (2018)*** also includes a plasmamembrane Ca pump (PMCA) in the soma, dendrites and spines, which we added in this model, and a sodium/calcium exchanger in the spines, which we ommited, because with the parameters from ***Dorman et al. (2018)***, it did not noticeably alter the spine [Ca]_NMDA_ concentration. In the calcium diffusion model in ***Dorman et al. (2018)***, calcium originating from all sources, which in that study are NMDARs and voltage–gated calcium channels (VGCCs), and a minor, 0.1% contribution of the AMPAR current, diffuses throughout the SPN model. Because the calcium buffer parameters are optimized to such a scenario, we also included calcium from the Ca_v_2.1 (Q-type), Ca_v_2.2 (N-type), Ca_v_2.3 (R-type), Ca_v_3.2 and Ca_v_3.3 (T-type) channels in the SPN together with [Ca]_NMDA_ in the diffusion model. Indeed, non-selective blockade of voltage-gated calcium channels with mibefradil has shown that they are also necessary for inducing LTP in SPNs (***Fino et al., 2010)***. Mibefradil has been shown to block T-type, L-type, R-type, N-type and P/Q-type calcium channels in different experimental preparations, so, apart from the L-type channels, we included all of them in the diffusion model (***Randall and Tsien***, ***1997; Viana et al., 1997; Aczél et al., 1998; Eller et al., 2000; Leuranguer et al., 2001)***. What the role of each of these channels is in LTP in SPNs requires further study with selective blockers.

**Table 2.**
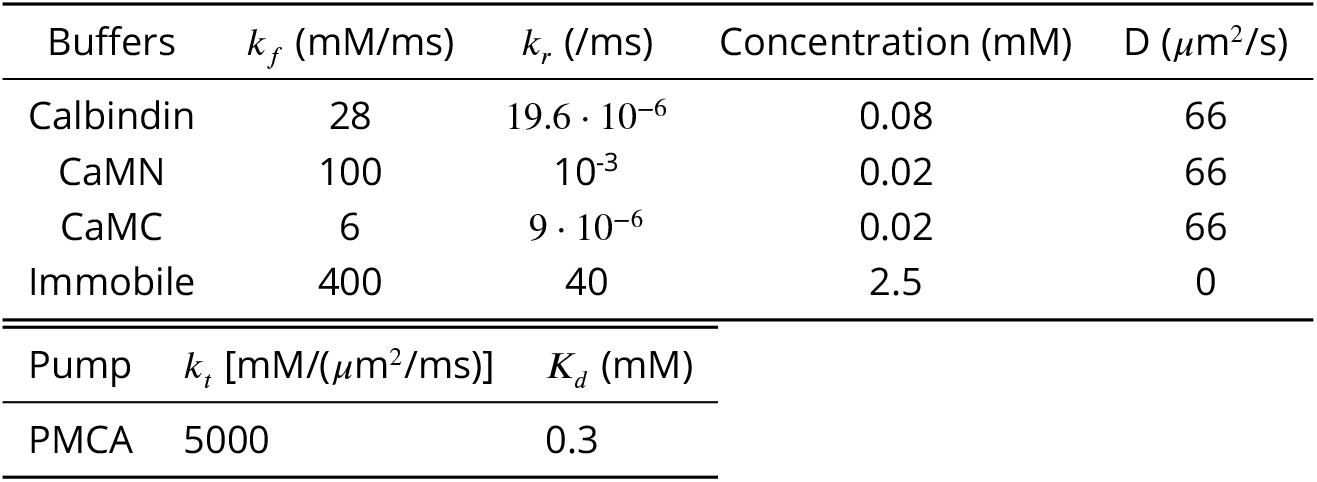
The parameters for the axial calcium diffusion model. Compared to ***Dorman et al. (2018)***, we have reduced the concentration of the immobile calcium buffer from 2.5 mM to 0.15 mM and changed the catalytic rate of the PMCA. The diffusion coefficient of Ca^2+^ is 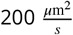 (same as in ***Dorman et al. (2018)***).

We also show how diffusion of only [Ca]_NMDA_ looks like in the SPN in Figure 13–figure supplement 1 (and Fig. 13C). The difference with the diffusion model used throughout the study, which includes VGCCs, is minor (Figure 13–figure supplement 1, cf. Figs. 2C, D). The qualitative and quantitative shapes of the voltage and calcium elevations are very similar (Figure 13–figure supplement 1A, cf. Fig. 2C). Regarding the amplitudes of these quantities, there is a 1 mV increase in the evoked somatic amplitude after glutamate spillover when only [Ca]_NMDA_ diffuses (Figure 13– figure supplement 1B_1_), and an obvious difference in the amplitude of [Ca]_NMDA_ (Figure 13–figure supplement 1B_3_). Nevertheless, the threshold nonlinearity in the [Ca]_NMDA_ amplitude is still present, and the learning rule would work the same provided the upper plasticity threshold Θ_LTP_ is placed in the region of the nonlinearity.

Even though mibefradil also affects L-type channels, we did not include them in the axial diffusion model together with [Ca]_NMDA_ because L-type calcium channels frequently form calcium microdomains, where calcium entry and diffusion is restricted within a very small, micrometer range (***Berridge***, ***2006; Chen and Sabatini***, ***2012; Parekh***, ***2008)***. In the microdomain, the L-type channels can be physically coupled in supramolecular protein complexes, further contributing to the local effect of the calcium influx in activating the relevant signaling networks, and this is also the case for Ca_v_1.3 channels in SPNs (***Olson et al., 2005; Stanika et al., 2015)***. Because of this, we assume that such highly localized [Ca]_L-type_ signals trigger LTD in the synapse, and calculated [Ca]_L-type_ with a separate, phenomenological pool model, which describes [Ca]_L-type_ in a thin cylindrical shell under the membrane. The implementation follows that of ***Wolf et al. (2005)***, according to Eq. 6. In this model [Ca]_L-type_ decays with a time constant *τ*_Ca_, modeling Ca^2+^ diffusion only phenomenologically, i.e no Ca^2+^ ions move between dendritic compartments. Compared to the very simplest pool models, this pool model also includes a Ca^2+^ pump that extrudes intracellular Ca^2+^ to the extracellular space. *F* is the Faraday constant, *k*_*t*_ is the catalytic activity of the pump, *K*_*d*_ is the pump dissociation constant for Ca^2+^, *d* is the depth of the submembrane shell where Ca^2+^ accumulates, and *k* and *p* are phenomenological parameters used in ***Wolf et al. (2005)*** used to balance Ca^2+^ influx and efflux. The values for the parameters in Eq. 6 are given in Table 3.

**Table 3.**
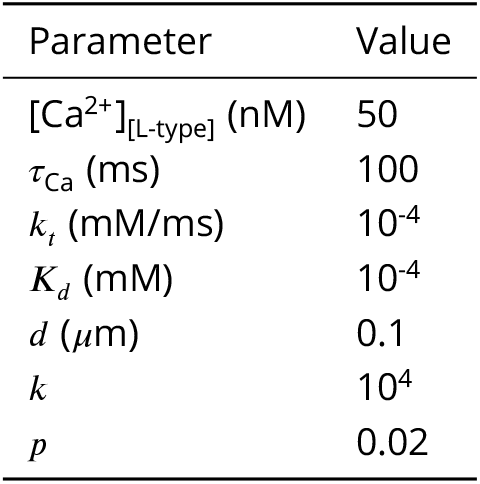
The parameters for the pool model for [Ca]_L-type_.

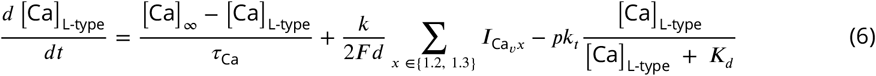

In this pool model we included calcium from both Ca_v_1.2 and Ca_v_1.3 channels, since there are no pharmacological agents that selectively block only one of the channels, i.e. the compounds used in the experiments which have determined Ca_v_1.3 as necessary for LTD in cortico-striatal synapses would also block Ca_v_1.2 channels in SPNs (***Berger and Bartsch*** (***2014)***; nimodipine in ***Wang et al. (2006)*** and ***Shindou et al. (2011)***; also, other non-selective L-type channel blockers are used to block LTD in the following studies: nifedipine in ***Calabresi et al. (1994)***; nitrendipine in ***Kreitzer and Malenka*** (***2005)***; mibefradil in ***Fino et al. (2010)***; finally, experiments in ***Plotkin et al. (2013)*** have suggested that other VGCCs than Ca_v_1.3 may be involved in activating calcium-induced calcium release, which is necessary for LTD).

### Synaptic inputs

There are two different models of synapses used in this study. One model – the dual exponential model – is used in describing the background noise, which consists of glutamatergic and GABAergic synapses, distributed across the dendrites according to the distribution measured in ***Cheng et al. (1997)***. These synapses are fixed (they have no plasticity) because they are activated at a low rate of 1 Hz. Compared to the 30 ms activation window in Fig. 3, this rate is low enough so that we decided to neglect any background noise synapses that might be co-active in the same time window and thus become eligible for plasticity together with the feature-carrying synapses. Because there are many synapses representing the background noise, this assumption also lowers the computational cost – if they were plastic, the learning rule would need to be computed for each of them.

The other model, used to describe the synapses carrying the features, is a saturating synapse model. We chose a saturating synapse model because we activate these synapses with 3 spikes each within a 30 ms window, and wanted to avoid any unrealistically high conductance values that could otherwise arise in non-saturating models.

#### Dual exponential model

The dual exponential synaptic model (a difference of two exponential functions) for a spike arriving at time 0 is represented by:

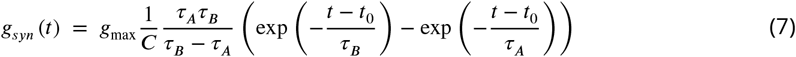

where *g*_max_ is the maximal synaptic conductance, and *τ*_*A*_ and *τ*_*B*_ are the rise and decay time constants. *C* is a normalization factor given with:

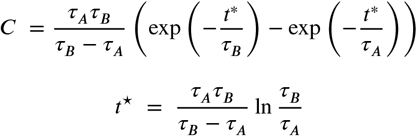

where *t*^⋆^ is the position of the maximum of the dual exponential function. This normalization ensures that the time-varying part of the dual exponential reaches a peak value of 1, which is then scaled by the maximal conductance *g*_max_. (The implementation in NEURON contains a synaptic weight parameter, as well. Since in the dual exponential model we have fixed the weight to 1, it is omitted from Eq. 7.)

The synaptic current arising from the conductance *g*_*syn*_ is:

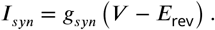

Excitatory synapses have both AMPA and NMDA components, and NMDA synapses in addition have a Mg^2+^ block described by a sigmoidal function:

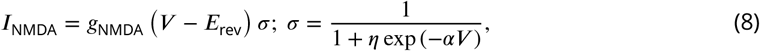

with parameters *α* = 0.062 and *η* = 0.38. The parameters of the dual exponential model for the glutamatergic AMPA and NMDA synapses, and the GABAergic synapses are given in Table 4.

**Table 4.**
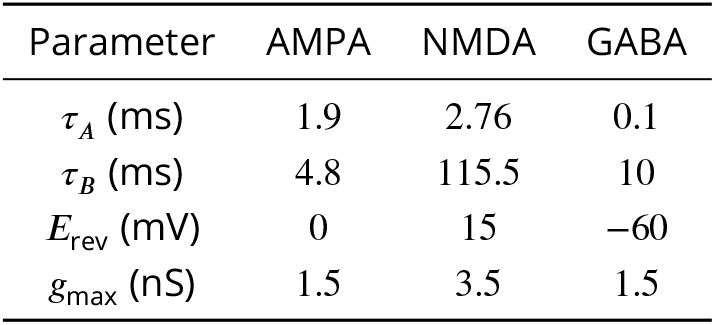
The parameters in the dual exponential synaptic model.

Finally, because the SPN model contains less electrical compartments than the number of synapses reported in ***Cheng et al. (1997)***, this implies that several background noise synapses will arrive in a single compartment. As such, their synaptic inputs will be integrated in the same voltage variable. Because of this, we have represented such multiple synapses arriving in one compartment with a single synapse instead, whose input frequency is scaled by the number of synapses it represents. This is why a non-saturating synapse model, such as the dual exponential model, was necessary for the background noise synapses. This also greatly increases simulation speed.

#### Saturating synapse model

The saturating synapse models are taken from ***Gao et al. (2021)***, which are a variation of the saturating synapse models in ***Destexhe et al. (1994)***. They are kinetic models which operate according to two different kinetic schemes depending on the presence or absence of neurotransmitters. When neurotransmitters arrive at the postsynaptic site, receptor dynamics are described by a kinetic scheme that models switching between closed (C) and open (O) states with rates of *α* and *β*:

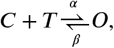

where [*T*] is the transmitter concentration. After neurotransmitter has been cleared from the synaptic cleft, receptor dynamics evolves according to a kinetic scheme that describes just the closing of the receptor with a rate *β*:

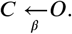

In this model, when a presynaptic spike arrives, neurotransmitter levels always reach a fixed saturating concentration in the synaptic cleft, [*T*]_max_, which lasts for a short time *T*_dur_ (a short glutamate lifetime in the synaptic cleft of cultured hippocampal synapses has been reported in ***Clements et al. (1992)***). If another presynaptic spike arrives while the neurotransmitter pulse is still “on”, the pulse duration is lengthened by *T*_dur_.

The synaptic conductance *g*_*syn*_ in this model is represented by the fraction of glutamate receptors in the open state, [*O*]:

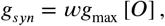

where *g*_max_ is again the maximal synaptic conductance, and *w* is the synaptic weight which is modified by the learning rule. The parameters in the saturating synapse model are given in Table 5.

**Table 5.** The parameters in the saturating synapse model.

| Parameter | AMPA | NMDA | eNMDA |
| --- | --- | --- | --- |
| $E_{\text{rev}}$ (mV) | 0 | 15 | 15 |
| $g_{\max}$ (nS) | 1.5 | 2.5 | 4.5 |
| $T_{\text{dur}}$ (ms) | 1 | 1 | $50 + 200w$ |
| $[T]_{\max}$ (mM) | 1 | 1 | 0.02 |
| $\alpha$ [/(ms mM)] | 12.5 | 4 | 4 |
| $\beta$ (/ms) | 0.25 | 0.01 | 0.01 |
| $K_d$ ($\mu\text{M}$) | 20 | 2.5 | 2.5 |

The fraction of NMDAR current that contributes to calcium influx is 17.5%, between the values of 15.9% for receptors with GluN1-GluN2A subunit composition (Table 18 in ***Traynelis et al. (2010)***) and 18.5% (***Schneggenburger***, ***1996)***.

### Glutamate spillover

To produce plateau potentials which have an all-or-none jump in the voltage, i.e. a threshold nonlinearity (as suggested by existing *in vitro* experiments), we included glutamate spillover. A detailed study of glutamate spillover and its role in triggering plateau potentials is given in ***Trpevski et al. (2023)***, and here we briefly describe the model. Glutamate spillover is a phenomenon that occurs during repeated synaptic stimulation, when glutamate reuptake by the surrounding astrocytes is saturated and when the astrocytes may even reverse function from reuptake to glutamate releas (***Malarkey and Parpura***, ***2008)***. When this happens in a synaptic cluster, glutamate spills over from the synaptic cleft into the extrasynaptic space, activating extrasynaptic NMDARs (eNMDARs). To model glutamate spillover, we place eNMDARs in the dendritic shaft, at the base of each spine in a cluster. Glutamate spillover is activated when a threshold level of clustered synaptic stimulation is reached – the “glutamate threshold”. In this study, the glutamate threshold is reached when 20 synapses with a weight *w* > 0.5 are activated (with 3 spikes each). Note that this threshold is equivalent to activating 10 synapses with a weight *w* > 1.0, or 40 synapses with a weight *w* > 0.25, i. e. the summed weight of all activated clustered synapses should be greater than 10.

When glutamate spills over, i) an activating spike to the eNMDARs is delivered 1.5 ms after the glutamate threshold is reached, ii) it stays much longer around the eNMDARs than in the synaptic cleft (*T*_dur_ is much longer and depends on the synaptic weight *w*), and iii) its concentration around the eNMDARs is much lower than in the synaptic cleft ([*T*]_max_ is much lower) (Table 5 and ***Trpevski et al. (2023)***). Also, the eNMDARs represent about 64% of the total NMDA conductance in this study (Table 5).

### Stimulation protocol

The stimulation protocol in Fig. 3A for presenting the patterns (in random order), which are followed by dopamine feedback, contains several additional details that we list here. First of all, when the distributed or clustered synapses are activated within the 30 ms stimulation window, 3 input spikes arrive to each synapse within this window (as explained in the caption of Fig. 3). This is because at physiological extracellular calcium concentrations, single spikes do not induce plasticity (at least in the spike-timing-dependent protocols in ***Inglebert et al. (2020)***). Furthermore, the backward rate constants from ***Dorman et al. (2018)*** for calcium unbinding from the buffers are much lower than experimentally measured values for calmodulin and calbindin, making full calcium unbinding and equilibration of the calcium dynamics very long (***Faas et al., 2011; Nägerl et al., 2000)***. To avoid long simulation times on the computing cluster used for this study, 600 ms after a pattern is presented, we reset the values of the buffers, their bound forms to calcium, and the intra- and extracellular calcium concentrations back to their initial values before the first pattern is presented, thus “equilibrating” the system faster and shortening simulation time. As can be seen from Fig. 2C, by 600 ms the plateaus are over, and both [Ca]_NMDA_ and [Ca]_L-type_ are back to their baseline levels, so this abrupt reset of the system does not affect learning and synaptic plasticity. And as can be seen from Fig. 3C, it neither affects the baseline voltage in the soma and dendrites before the arrival of a future pattern.

### Performance scores on the FBP and NFBP

In the FBP, a score of 50% means that the SPN is correct only half of the time. This happens if the SPN is silent for both the relevant and irrelevant pattern (correct output only for the irrelevant pattern), or if it spikes for both patterns (correct output only for the relevant pattern). A score of 100% indicates it always spikes for the relevant pattern, and is always silent for the irrelevant pattern, and the FBP is perfectly solved. The threshold score for solving the FBP is 75%, exemplified with the situation where the SPN is always silent for the irrelevant pattern, and spikes at least half of the time for the relevant pattern. A score of 0% would indicate that the SPN’s output is incorrect all the time, being silent for the relevant pattern, and spiking for the irrelevant pattern (this does not happen in the simulations).

In the NFBP, a score of 50% also means that the SPN output is correct only half of the time. An example of such a score is when the SPN is silent for all four patterns, or spikes for all four patterns. Similarly, a score of 100% indicates it always spikes for the relevant patterns, and is always silent for the irrelevant patterns. A score of 75% is exemplified with the situation where the SPN is silent for the two irrelevant patterns, and spikes for only one of the relevant patterns. The threshold score for solving the NFBP is 87.5%, where the SPN would also spike for the other relevant pattern at least half of the time. Alternatively, a score of 75% can be achieved by the SPN spiking for both relevant patterns, but being silent for only one irrelevant pattern. Similarly, the threshold score of 87.5% can be reached by being silent for the other irrelevant pattern at least half of the time.

## Acknowledgments

DT thanks Ana Kalajdjieva for illustrating the mouse brain. Simulations were performed on resources provided by the National Academic Infrastructure for Supercomputing in Sweden (NAISS) at PDC KTH.

---

**Figure 1—figure supplement 1.**
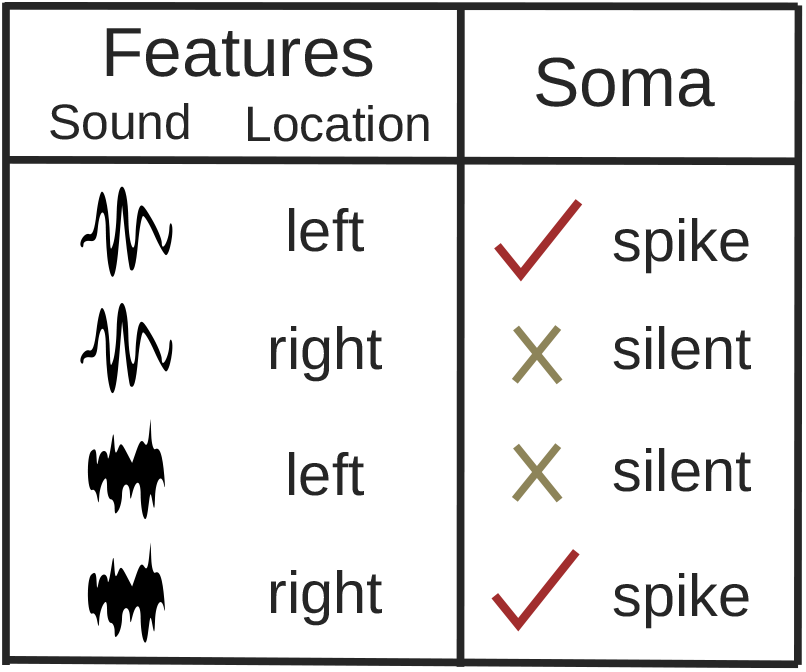
An example of the NFBP with features relevant for the striatum. The example is inspired by the tasks used in ***Bernklau et al. (2024)***, and consists of pairing a sound frequency with a location where the sound originates from. Two sound frequencies (low and high) and two locations (left and right) were used, with a reward being provided to only two of the feature combinations (low sound to the left and high sound to the right).

**Figure 3—figure supplement 1.**
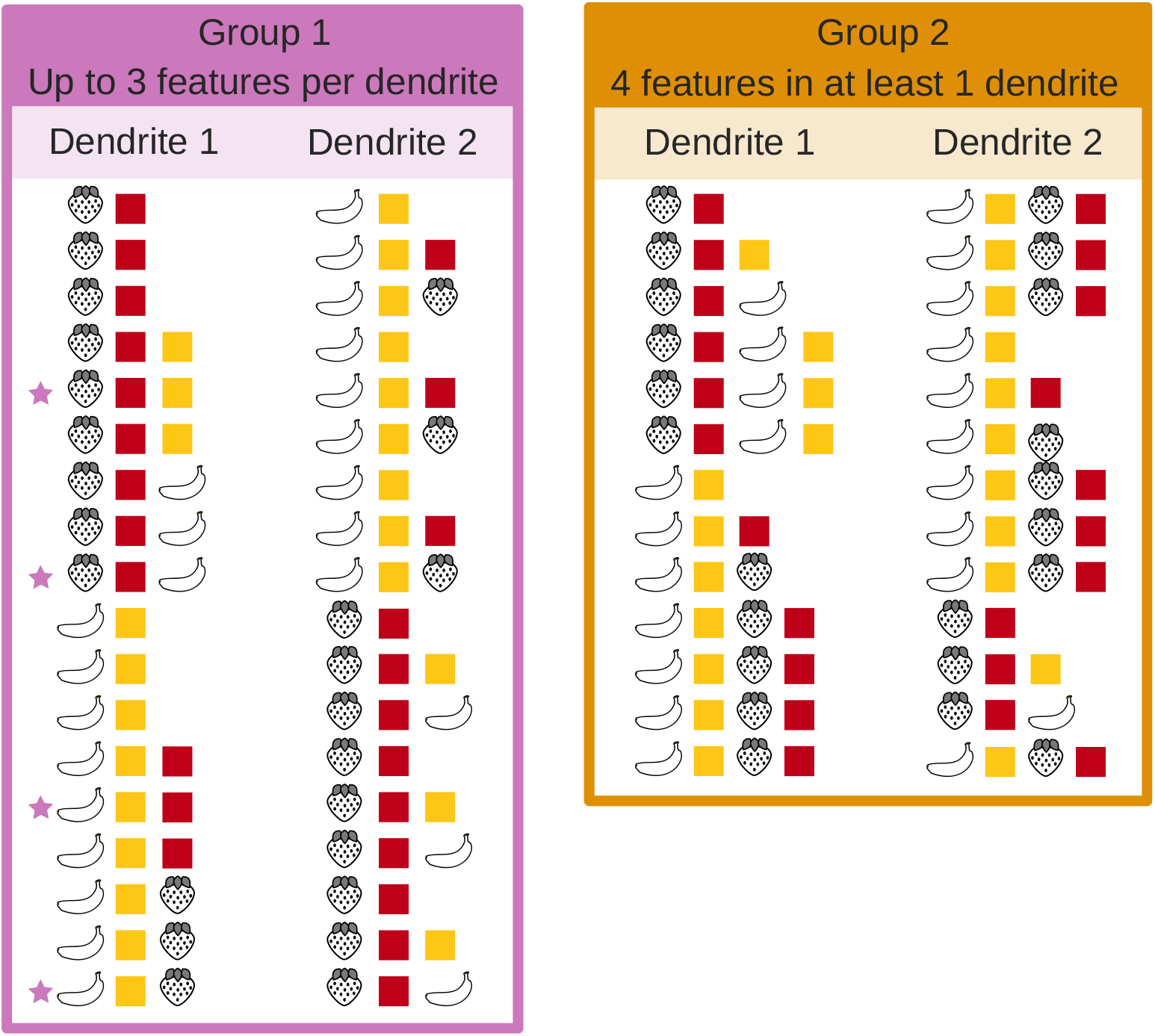
All possible input configurations of four features on two dendrites that allow the NFBP to be learned. They are divided in two groups: those with up to 3 features per dendrite (group 1), and those with 4 features in at least 1 dendrite (group 2). We have made this division because when a dendrite is innervated with all four features (group 2), learning to solve the NFBP depends on the order in which the patterns arrive. In this case, often only one of the relevant patterns is stored by the neuron (the same pattern in both dendrites), and solving the NFBP requires additional mechanisms, such as branch plasticity or inhibition (***Legenstein and Maass, 2011; Trpevski et al., 2026)***.

**Figure 4—figure supplement 1.**
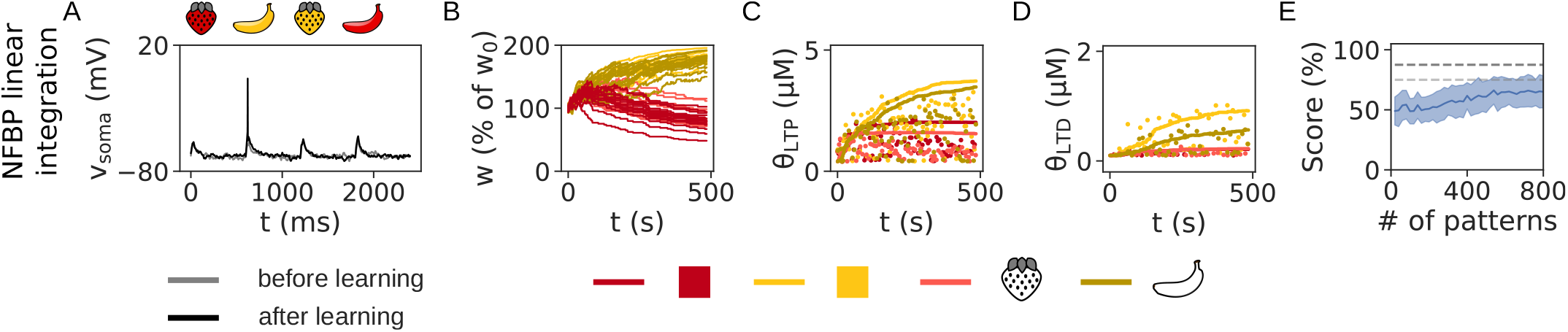
. The NFBP cannot be solved with linear integration at the soma. Each feature is represented by 15 distributed synapses across the dendrites. Only one relevant pattern is learned by the neuron. (A) The somatic voltage before and after learning. No plateaus are evoked as in Fig. 3B_3_ because synapses are distributed across the dendrites. After learning, the neuron spikes for only one of the relevant patterns. (B–D) The evolution of synaptic weights (B), LTP thresholds (C) and LTD thresholds (D). In (B) synaptic weights for all synapses are shown, while in (C) and (D) only one synapse per feature is chosen to show its calcium threshold. Dots represent the amplitudes of [Ca]_NMDA_ and [Ca]_L-type_, respectively, during pattern presentation. In this simulation, only the “yellow banana” is stored by the neuron’s distributed synapses (B). Because no plateaus are evoked, [Ca]_NMDA_ never reaches the upper threshold for LTP, Θ_LTP_ (C), and as a result, synapses are strengthened close to the maximal value *w*_max_, as in the FBP with linear integration (B, cf. Fig. 4B_1_). (D) Performance on the NFBP averaged over 50 trials. The dashed lines mark the threshold scores of 75% and 87.5%.

**Figure 4—figure supplement 2.**
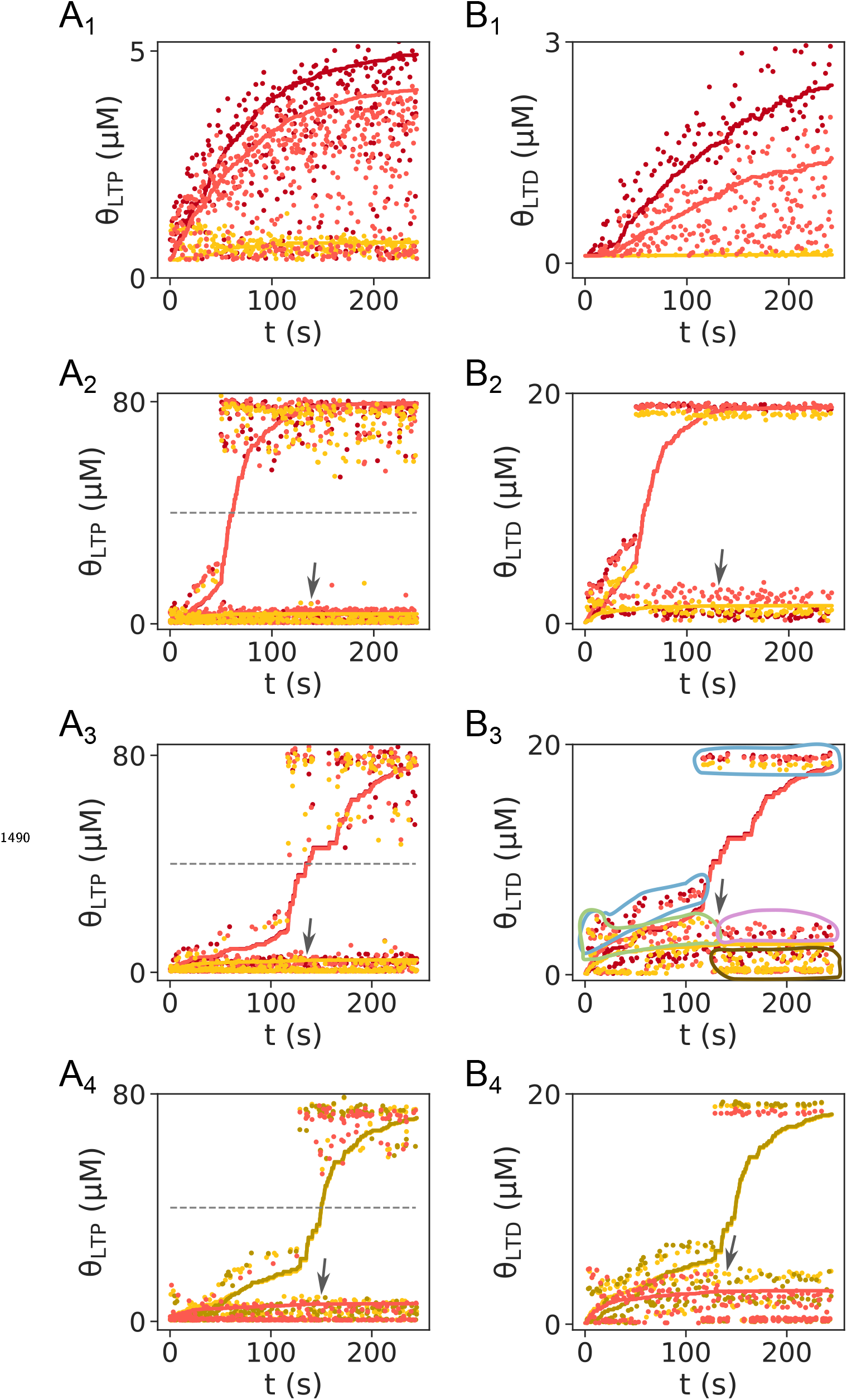
The calcium thresholds and calcium amplitudes from Fig. 4 shown in larger panels with greater detail. (A) LTP thresholds for the FBP with distributed (A_1_) and clustered (A_2_) synapses, and for the NFBP (A_3_, A_4_). (B) LTD thresholds for the FBP with distributed (B_1_) and clustered (B_2_) synapses, and for the NFBP (B_3_, B_4_). In (B_3_), regions with calcium amplitudes of interest are highlighted (the main text refers to these regions when describing the dynamics of the weights, thresholds, and calcium concentrations). The same regions exist in all panels, although they are not highlighted.

**Figure 4—figure supplement 3.**
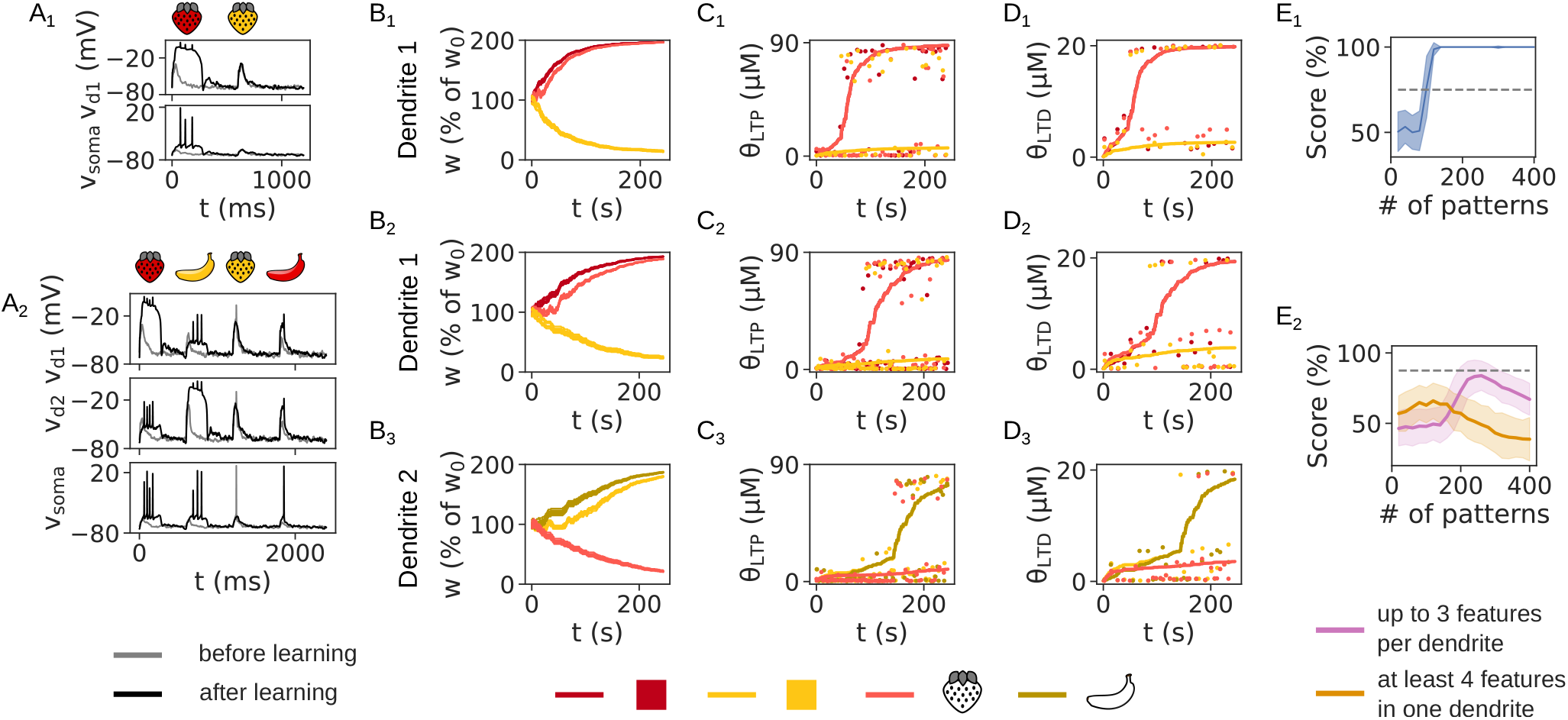
The effect of no upper threshold Θ_LTP_ on supralinear integration. (A) The somatic and dendritic voltages before and after learning in the FBP (A_1_) and NFBP (A_2_). Not having an upper threshold causes somatic spiking for the irrelevant patterns after learning in the NFBP, but not the FBP. (B-D) The evolution of synaptic weights (B), LTP thresholds (C) and LTD thresholds (D) in the FBP (B_1_-D_1_) and the NFBP (B_2_-D_2_ and B_3_-D_3_). Strengthened synapses saturate at their maximal levels, also driving more weakening in the weakened synapses. Having no upper threshold does not affect the thresholds of the strengthened synapses, but causes the thresholds of the weakened synapses to stabilize at higher levels. (E) Performance on the FBP (E_1_) and NFBP (E_2_). The same number of trials as in Fig. 3D is used.

**Figure 4—figure supplement 4.**
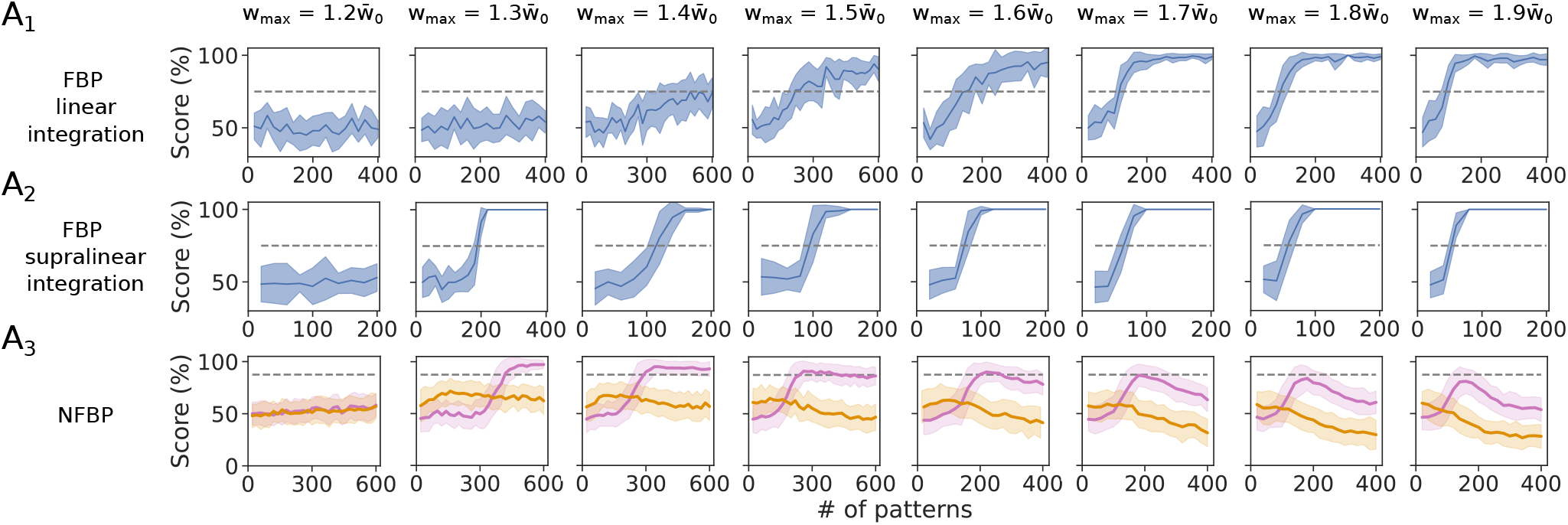
A parameter scan of *w*_max_ using the rule without an upper LTP threshold. (A) The performance on the FBP with linear (A) and supralinear integration (B) and on the NFBP (C) is shown when the maximal weight is varied from 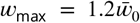 to 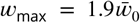. (For the FBP, the performance is averaged over 10 trials, and for the NFBP over 4 trials per input configuration, divided into the two groups in Figure 3–figure supplement 1, with in 72 and 52 trials per group, respectively.) There is no single *w*_max_ value that solves all three tasks. Only 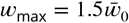 comes close to solving all three of them (with the score on the NFBP being at the border for solving the task). This suggests that if a range of values for *w*_max_ that solves all three tasks exists, it is very narrow, and that such a solution of the three tasks without an upper LTP threshold is not robust to variations in *w*_max_.

**Figure 4—figure supplement 5.**
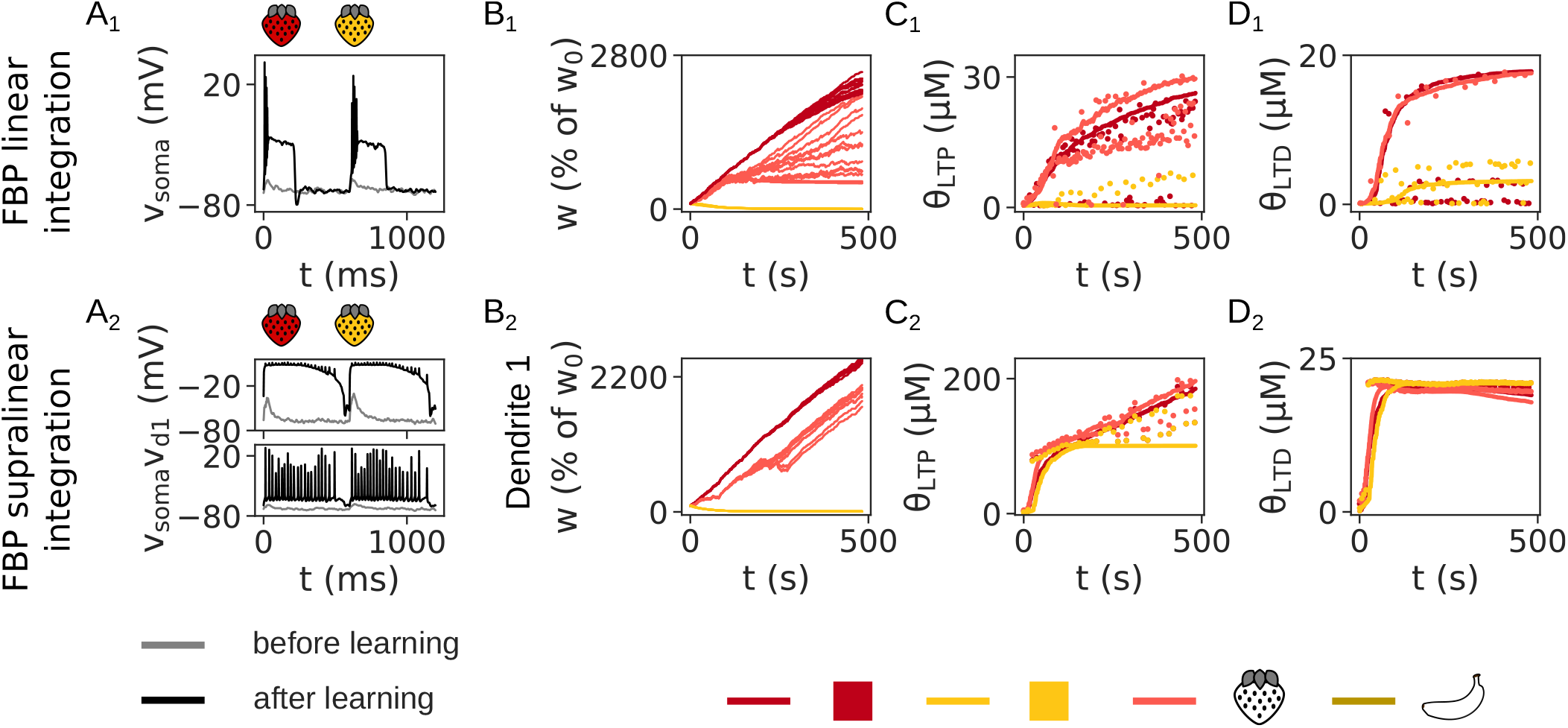
The maximal weight *w*_max_ prevents unbounded weight growth. In these simulations, no *w*_max_ was used, causing the weights to increase until calcium concentration saturates in the spines. (A) The somatic and dendritic voltages in the FBP with linear (A_1_) and supralinear (A_2_) integration before and after learning (gray and black traces, respectively), elicited by the relevant and irrelevant pattern. *w*_max_ ensures firing rate stability in the FBP with linear integration – without it, the neuron enters into a depolarization block after learning. In the FBP with supralinear integration, the plateaus ensure firing rate stability – regardless of how high the synaptic weights are, a plateau’s voltage is limited by the NMDAR reversal potential, providing dynamic range compression (see also Figs. 3E, 4D and 12 in ***Oikonomou et al. (2012)***). (B) The evolution of the weights in the FBP with linear (B_1_) and supralinear (B_2_) integration. (C, D) The LTP (C) and LTD threshold (D) shown for a single synapse per feature in the FBP with linear (C_1_, D_1_) and supralinear (C_2_, D_2_) integration. No upper threshold for LTP was used in the FBP with supralinear integration in order to showcase the effect of not having a maximal weight cap, *w*_max_. A modified version of the spillover model was used here in which the synaptic weight does not prolong the plateaus’ duration (otherwise, an increasingly long inter-pattern interval would have been needed to allow the plateau to finish, significantly prolonging the simulation.)

**Figure 4—figure supplement 6.**
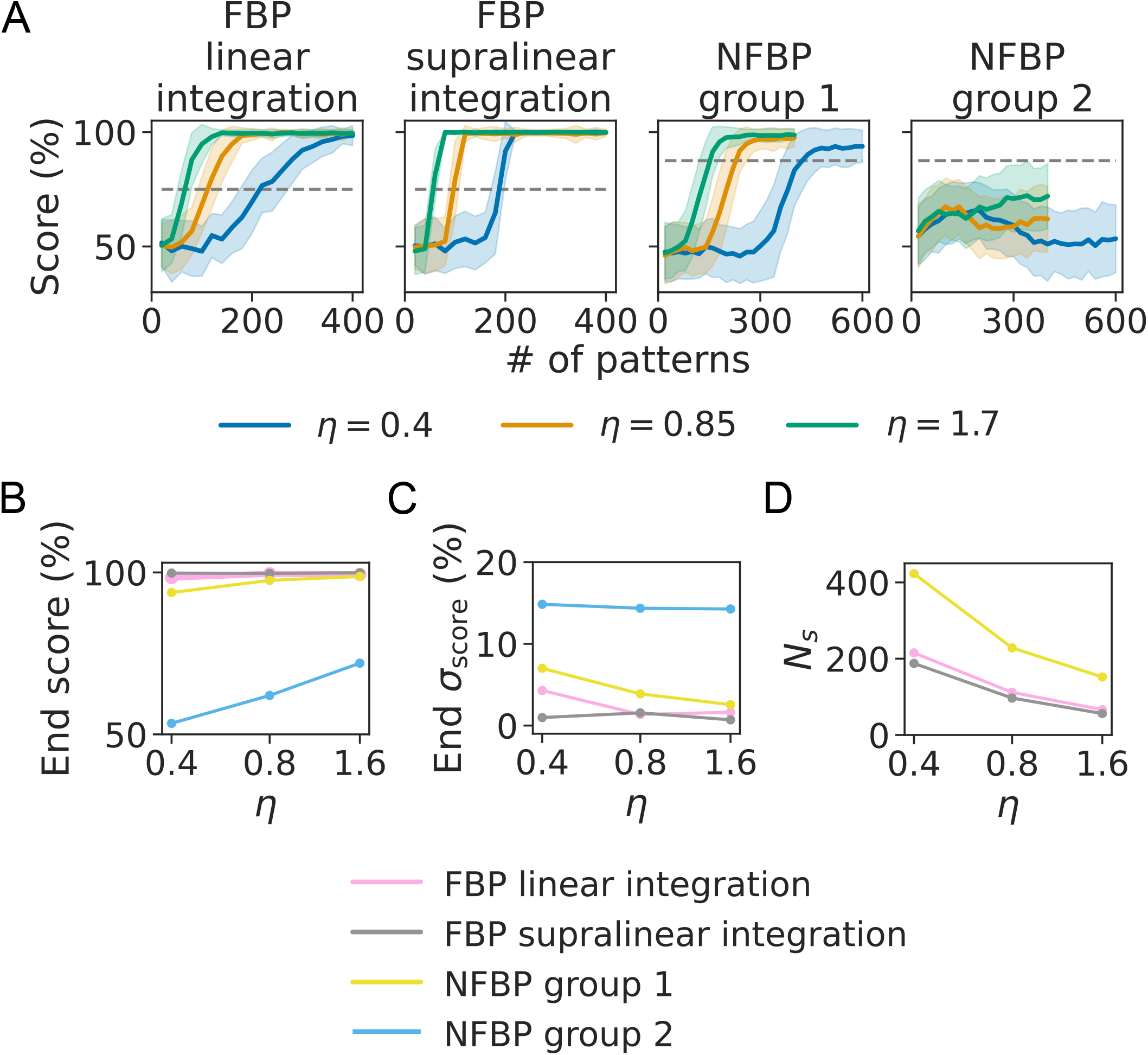
The effect of the learning rate *η* on learning. (A) The average score on the FBP and NFBP for three values of the learning rate, *η* ∈ {0.4, 0.85, 1.7} (the number of trials is the same as in Fig. 3D). Increasing the learning rate causes faster learning of the tasks (less pattern presentations are needed). (B) The score at the end of learning varies little with *η*, except in group 2 for the NFBP, where a higher learning rate increases performance, so that one of the relevant patterns is remembered. (C) The standard deviation at the end of learning also varies little with *η* (there is a 5% decrease for the NFBP, group 1 only). (D) The speed of learning measured by the number of patterns, *N*_*s*_, needed to reach the score threshold for solving each task (the dashed line in (A)). The speed of learning to solve the tasks is affected by the learning rate *η*, as is also seen in (A). Increasing the learning rate speeds up learning.

**Figure 4—figure supplement 7.**
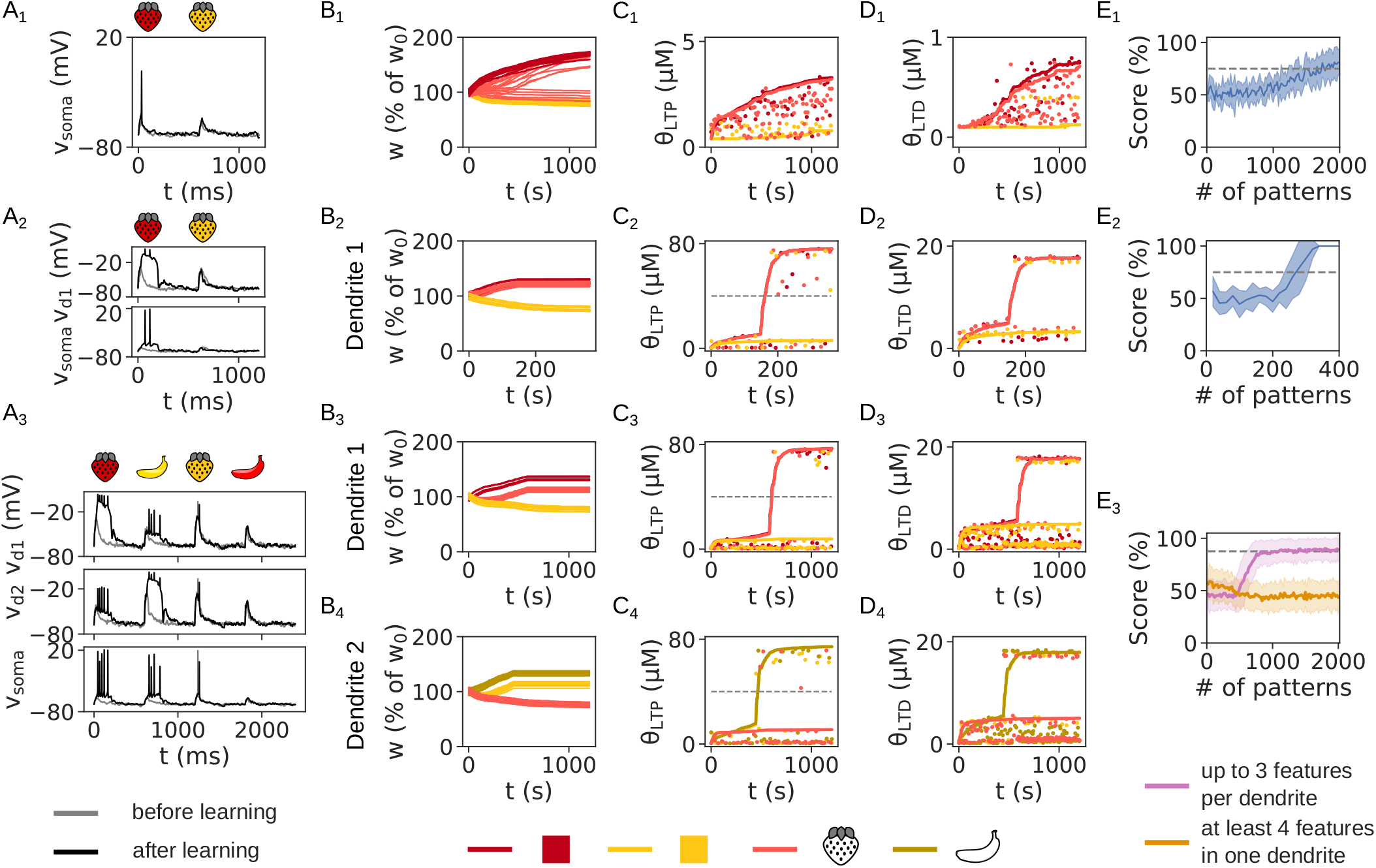
The effect of a very low learning rate on learning (*η* = 0.1). (A) The somatic and dendritic voltages in the FBP with linear (A_1_) and supralinear (A_2_) integration, and the NFBP (A_3_), before and after learning (gray and black traces, respectively), elicited by the relevant and irrelevant pattern(s). (B-D) The evolution of synaptic weights (B), LTP thresholds (C) and LTD thresholds (D) in the FBP with linear (B_1_-D_1_) and supralinear (B_2_-D_2_) integration, and the NFBP (B_3_-D_3_ and B_4_-D_4_). The low value of *η* cuses small weight updates, resulting in less weakening than is necessary to solve the NFBP (E_3_), and significantly prolonging the learning of the FBP with linear integration (E_1_). In (B) all synaptic weights for the features are shown, while in (C) and (D) only one synapse per feature is chosen to show its calcium threshold (solid lines). Dots in (C) and (D) represent the amplitudes of [Ca]_NMDA_ and [Ca]_L-type_, respectively, during pattern presentation. Dashed lines in (C_2_–C_4_) show the upper threshold for LTP, Θ_LTP_. (E) The performance on the FBP with linear (E_1_) and supralinear (E_2_) integration, and on the NFBP (E_3_). In (E_1_) the score is averaged over 10 trials and (E_2_) the score is averaged over 50 trials. In (E_3_) the score is averaged over 4 trials per input configuration, divided into two groups (Figure 3–figure supplement 1), resulting in 72 and 52 trials per group, respectively.

**Figure 4—figure supplement 8.**
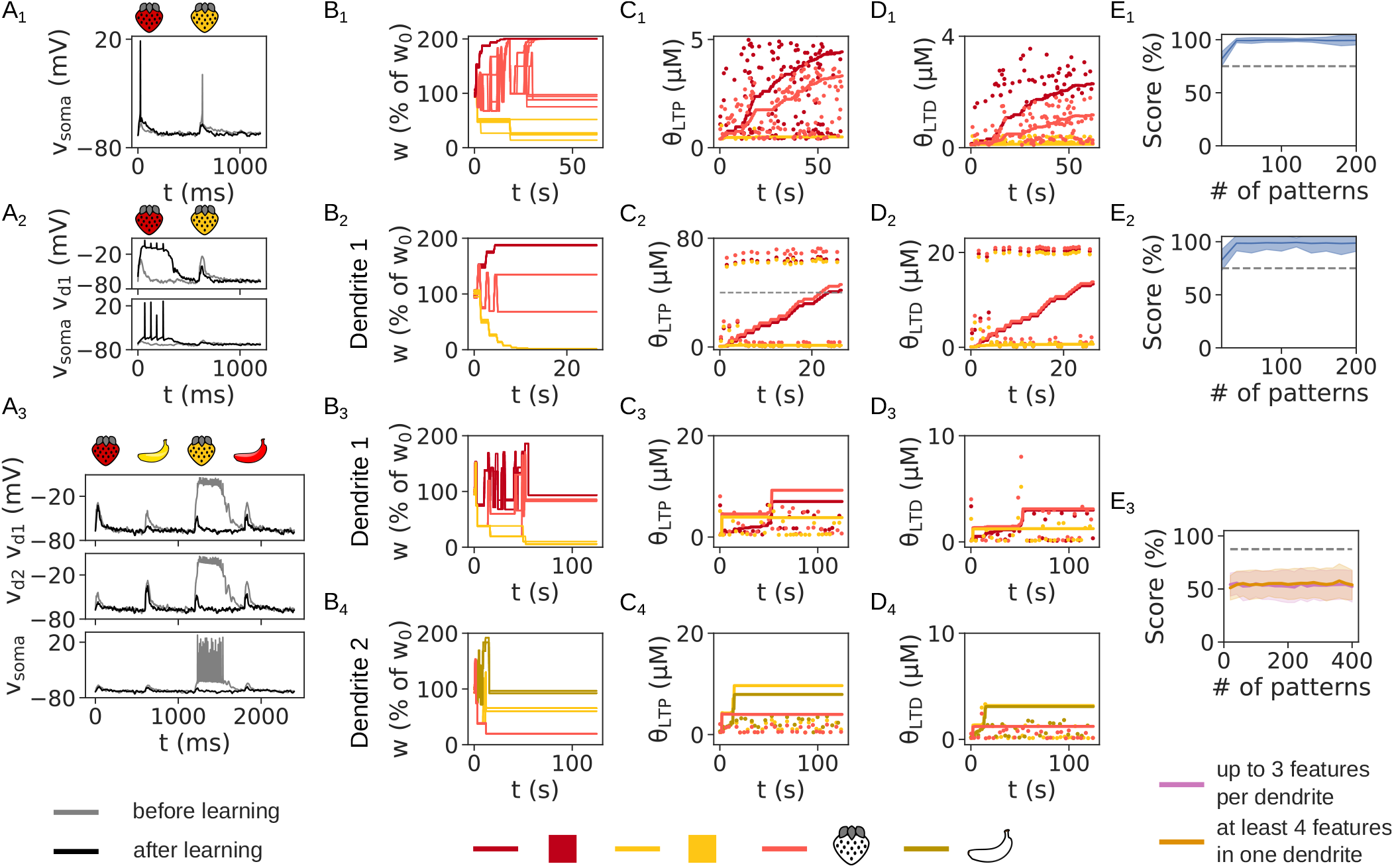
The effect of a very high learning rate on learning (*η* = 20). (A) The somatic and dendritic voltages in the FBP with linear (A_1_) and supralinear (A_2_) integration, and the NFBP (A_3_), before and after learning (gray and black traces, respectively), elicited by the relevant and irrelevant pattern(s). (B-D) The evolution of synaptic weights (B), LTP thresholds (C) and LTD thresholds (D) in the FBP with linear (B_1_-D_1_) and supralinear (B_2_-D_2_) integration, and the NFBP (B_3_-D_3_ and B_4_-D_4_). Large updates occur in the weights due to the high value of *η*, which in the NFBP cause calcium to fall below the thresholds and prematurely stop learning, without storing any pattern. The large weight updates even cause a plateau very early during learning, for the first irrelevant pattern (“yellow strawberry” in A_3_). In (B) all synaptic weights for the features are shown, while in (C) and (D) only one synapse per feature is chosen to show its calcium threshold (solid lines). Dots in (C) and (D) represent the amplitudes of [Ca]_NMDA_ and [Ca]_L-type_, respectively, during pattern presentation. Dashed lines in (C_2_–C_4_) show the upper threshold for LTP, Θ_LTP_. (E) The performance on the FBP with linear (E_1_) and supralinear (E_2_) integration, and on the NFBP (E_3_). In (E_1_) and (E_2_) the score is averaged over 10 trials. In (E_3_) the score is averaged over 8 trials per input configuration, divided into two groups (Figure 3–figure supplement 1), resulting in 144 and 104 trials per group, respectively.

**Figure 4—figure supplement 9.**
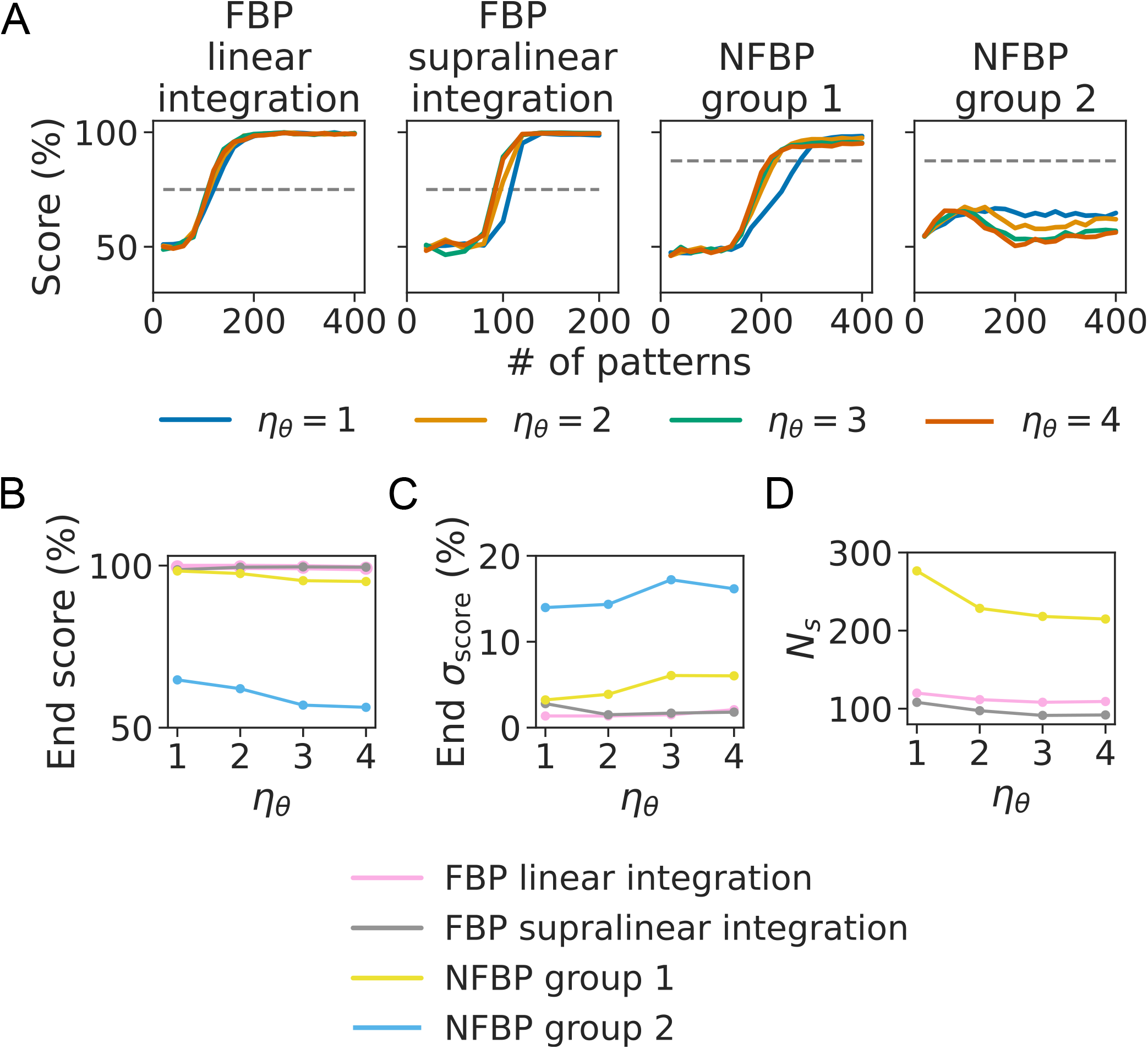
The effect of the metaplasticity rate *η*_*θ*_ on learning. (A) The average score on the FBP and NFBP for four values of the metaplasticity rate, *η*_*θ*_ ∈ {1, 2, 3, 4} (the standard deviation is ommited for figure clarity, but the number of trials is the same as in Fig. 3D). There is little effect of the metaplasticity rate on the final score (B), the final standard deviation of the score (C) and the speed of learning (D). (B) The score at the end of learning varies little with *η*_*θ*_ (only slightly for the NFBP). (C) The standard deviation at the end of learning also varies little with *η*_*θ*_ (up to 5% increase for the NFBP only). (D) The speed of learning measured by the number of patterns, *N*_*s*_, needed to reach the score threshold for solving each task (the dashed line in (A)). The speed of learning to solve the FBP varies very little with *η*_*θ*_, while that for solving the NFBP only shows a large increase between *η*_*θ*_ = 1 and *η*_*θ*_ = 2.

**Figure 4—figure supplement 10.**
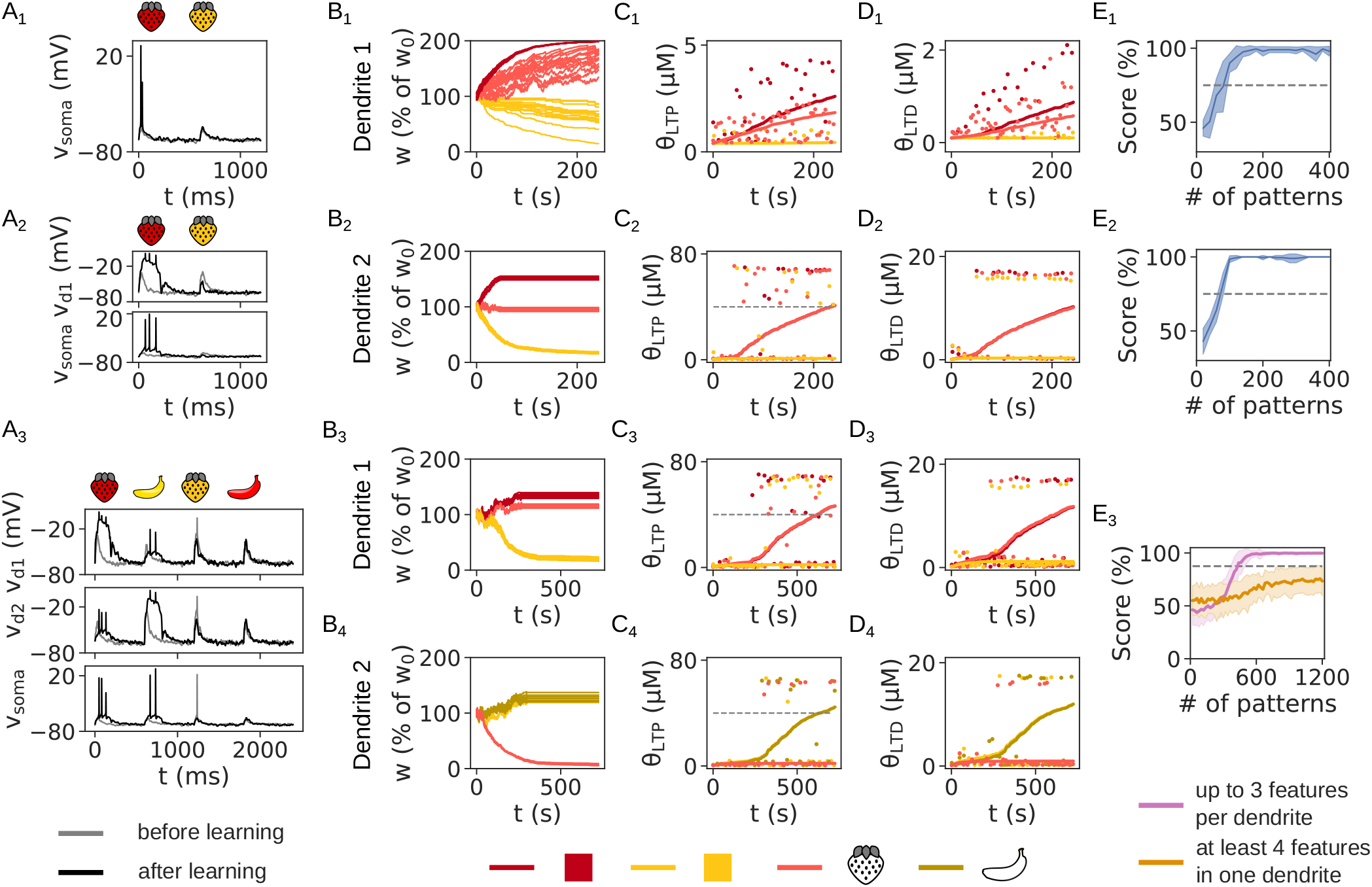
The effect of a low metaplasticity rate on learning (*η*_*θ*_ = 0.2). (A) The somatic and dendritic voltages in the FBP with linear (A_1_) and supralinear (A_2_) integration, and the NFBP (A_3_), before and after learning (gray and black traces, respectively), elicited by the relevant and irrelevant pattern(s). (B) The evolution of the weights in the FBP with linear (B_1_) and supralinear (B_2_) integration, and the NFBP (B_3_, B_4_). When *η*_*θ*_ is very low, the weights take a longer time to stabilize (note that the simulation on the NFBP is run three times longer). (C, D) The LTP (C) and LTD threshold (D) shown for a single synapse per feature in the FBP with linear (C_1_, D_1_) and supralinear (C_2_, D_2_) integration, and the NFBP (C_3,4_, D_3,4_). Because of the low metaplasticity rate, none of the thresholds reach the maximal calcium levels within the length of these simulations. (E) The performance on the FBP with linear (E_1_) and supralinear (E_2_) integration, and on the NFBP (E_3_). The low value of *η*_*θ*_ does not affect learning in the FBP (cf. Fig. 3D_1_, D_2_) but prolongs learning in the NFBP (cf. Fig. 3D_3_). In (E_1_) and (E_2_) the performance is averaged over 10 trials, and in (E_3_) over 4 trials per input configuration, which are divided into the two groups shown in Figure 3–figure supplement 1 (with 72 and 52 trials in the two groups, respectively).

**Figure 4—figure supplement 11.**
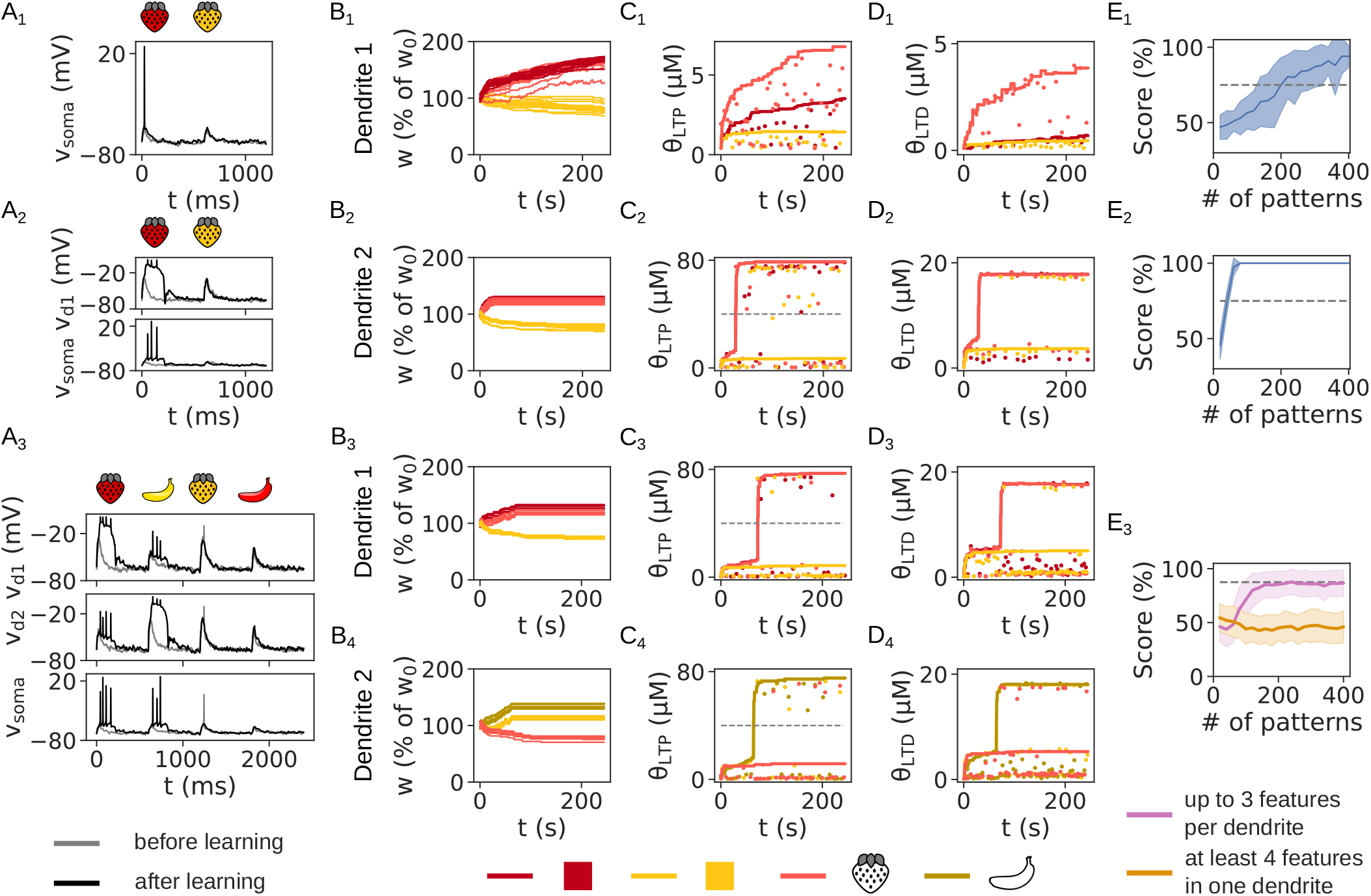
The effect of a low metaplasticity rate on learning (*η*_*θ*_ = 20). (A) The somatic and dendritic voltages in the FBP with linear (A_1_) and supralinear (A_2_) integration, and the NFBP (A_3_), before and after learning (gray and black traces, respectively), elicited by the relevant and irrelevant pattern(s). (B) The evolution of the weights in the FBP with linear (B_1_) and supralinear (B_2_) integration, and the NFBP (B_3_, B_4_). (C, D) The LTP (C) and LTD threshold (D) shown for a single synapse per feature in the FBP with linear (C_1_, D_1_) and supralinear (C_2_, D_2_) integration, and the NFBP (C_3,4_, D_3,4_). When *η*_*θ*_ is very high, the thresholds quickly reach their respective calcium levels, and stabilize the weights more quickly. As a result, the weights are modified less with respect to their initial values. (E) The performance on the FBP with linear (E_1_) and supralinear (E_2_) integration, and on the NFBP (E_3_). The high value of *η*_*θ*_ causes more variability in the performance for the FBP with linear integration because weights are not strengthened enough. It also lowers the performance on the NFBP because weakened clusters are not weakened enough. In (E_1_) and (E_2_) the performance is averaged over 10 trials, and in (E_3_) over 4 trials per input configuration, which are divided into the two groups shown in Figure 3–figure supplement 1 (with 72 and 52 trials in the two groups, respectively).

**Figure 7—figure supplement 1.**
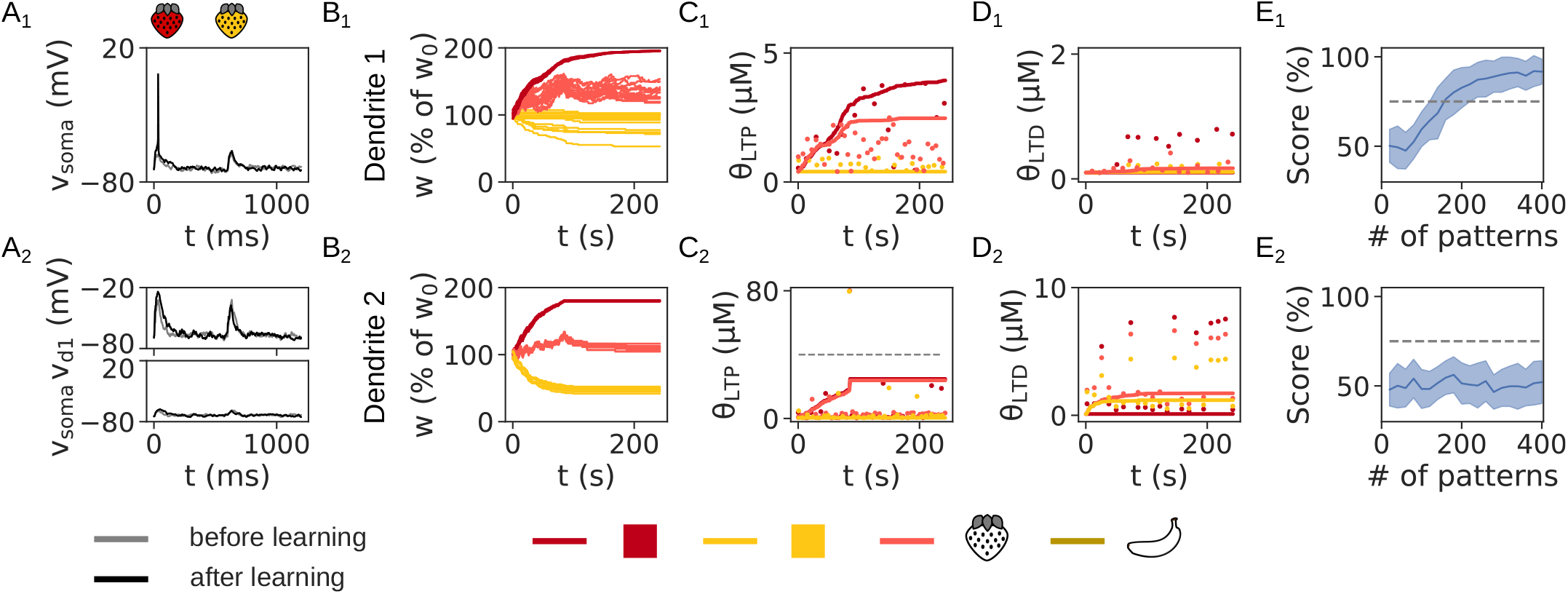
Learning outcome on the FBP with linear (A_1_-E_1_) and supralinear integration (A_2_-E_2_) with partial metaplasticity. (A) The somatic and dendritic voltage evoked by both patterns before and after learning. (B-D) The evolution of synaptic weights (B), LTP thresholds (C) and LTD thresholds (D) during learning. In (B) all synaptic weights are shown, and in (C, D) the thresholds for only one synapse per feature are shown. (E) The performance on the FBP, averaged over 50 trials.

**Figure 8—figure supplement 1.**
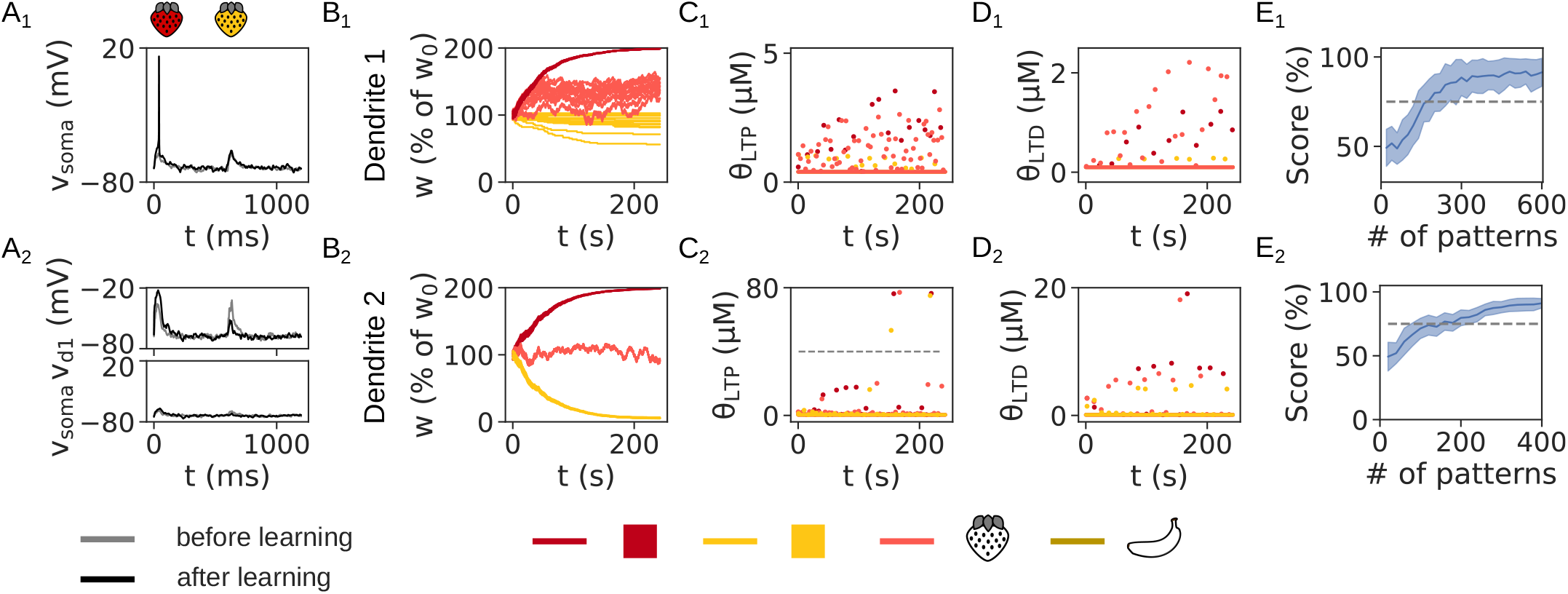
Learning outcome on the FBP with linear (A_1_-E_1_) and supralinear integration (A_2_-E_2_) without metaplasticity. (A) The somatic and dendritic voltage evoked by both patterns before and after learning. (B) The evolution of synaptic weights during learning. (C, D) The LTP (C) and LTD thresholds (D) are fixed. In (B) all synaptic weights are shown, and in (C, D) the thresholds for only one synapse per feature are shown. (E) The performance on the FBP, averaged over 50 trials.

**Figure 9—figure supplement 1.**
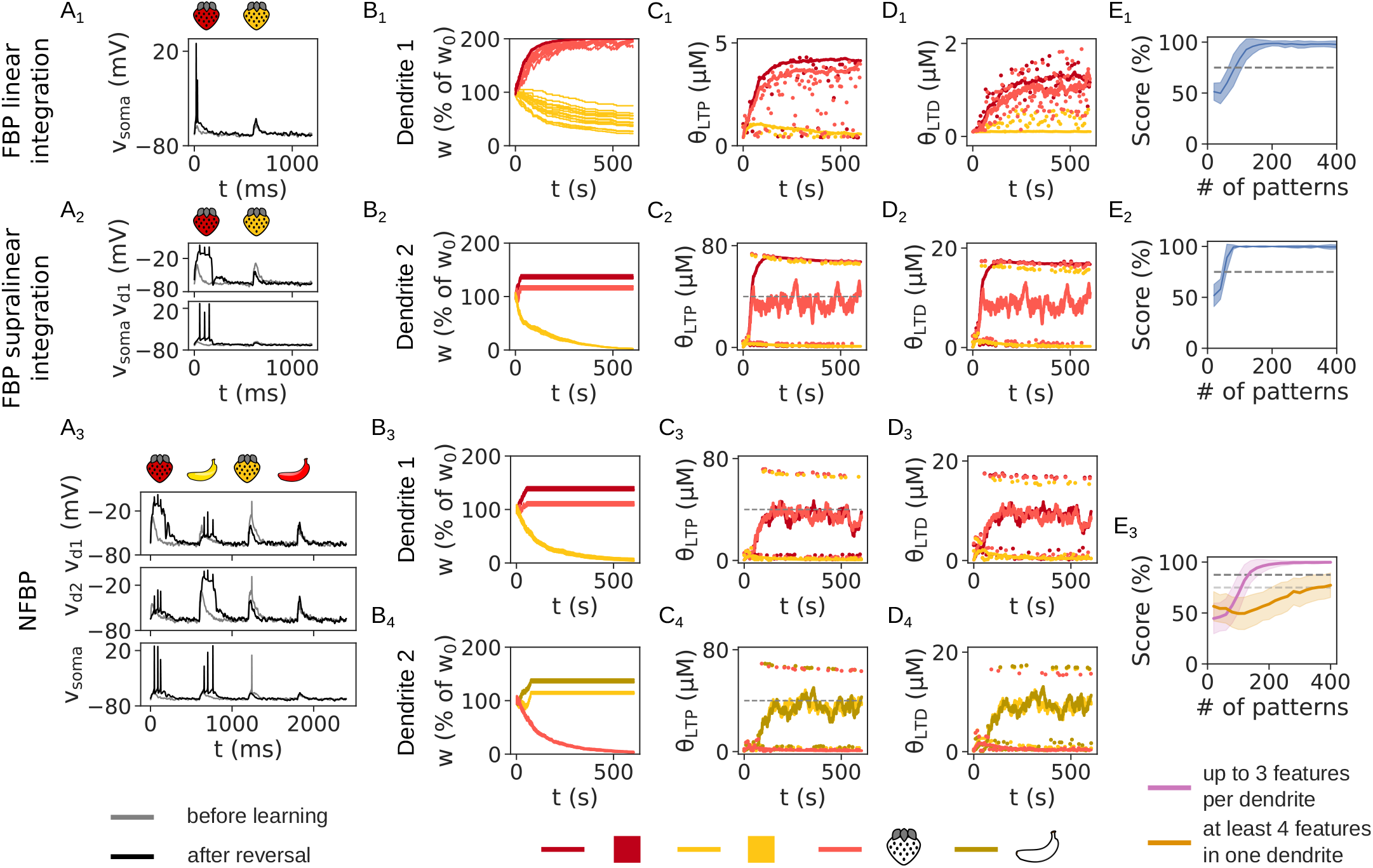
Relaxed metaplasticity also solves the FBP and NFBP. (A) The somatic and dendritic voltage before (gray traces) and after learning (black traces) for the FBP with linear (A_1_) and supralinear integration (A_2_) and the NFBP (A_3_).(B–D) The evolution of synaptic weights (B), LTP thresholds (C) and LTD thresholds (D) during learning. The thresholds for the shared features that need to be strengthened show large fluctuations in the cases of supralinear integration (C_2_–C_4_ and D_2_–D_4_). However, this does not influence the performance on the tasks. In (B) all synaptic weights are shown, and in (C, D) the thresholds for only one synapse per feature are shown. (E) The performance on the FBP with linear (E_1_) and supralinear integration (E_2_) and the NFBP (E_3_). Performance on the FBP is averaged over 50 trials, and in the NFBP over 12 trials per input configuration (divided into the two groups shown in Figure 3–figure supplement 1, with 216 and 156 trials, respectively).

**Figure 9—figure supplement 2.**
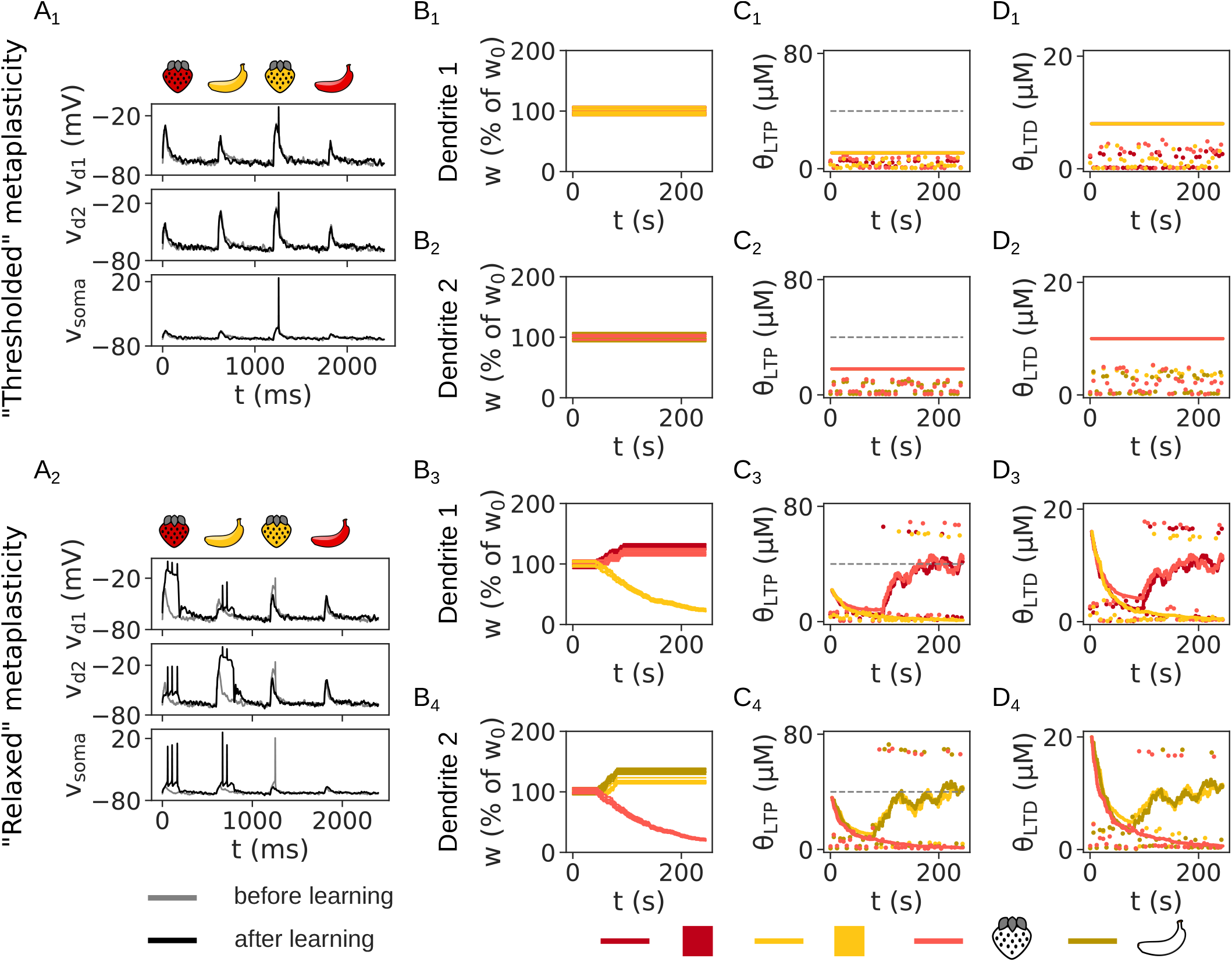
“Relaxed” metaplasticity can overcome initially high calcium thresholds, unlocking synapses and allowing learning. Only the NFBP is used as a demonstration. The somatic and dendritic voltage before (gray traces) and after learning (black traces) for “thresholded” (A_1_) and “relaxed” metaplasticity (A_2_). (B–D) The evolution of synaptic weights (B), LTP thresholds (C) and LTD thresholds (D) during learning. In (B) all synaptic weights are shown, and in (C, D) the thresholds for only one synapse per feature are shown. Dots in (C, D) represent calcium amplitudes evoked by the patterns. With “thresholded” metaplastiity, initializing thresholds to high values leaves the synapses locked (B_1_, B_2_), as calcium amplitudes always remain below the thresholds (C_1,2_, D_1,2_). The NFBP cannot be solved in this case (A_1_). On the other hand, with “relaxed” metaplasticity, the initially high thresholds adapt (C_3,4_, D_3,4_), thus unlocking the synapses and allowing learning (B_3_, B_4_). The NFBP is solved in this case (A_2_).

**Figure 10—figure supplement 1.**
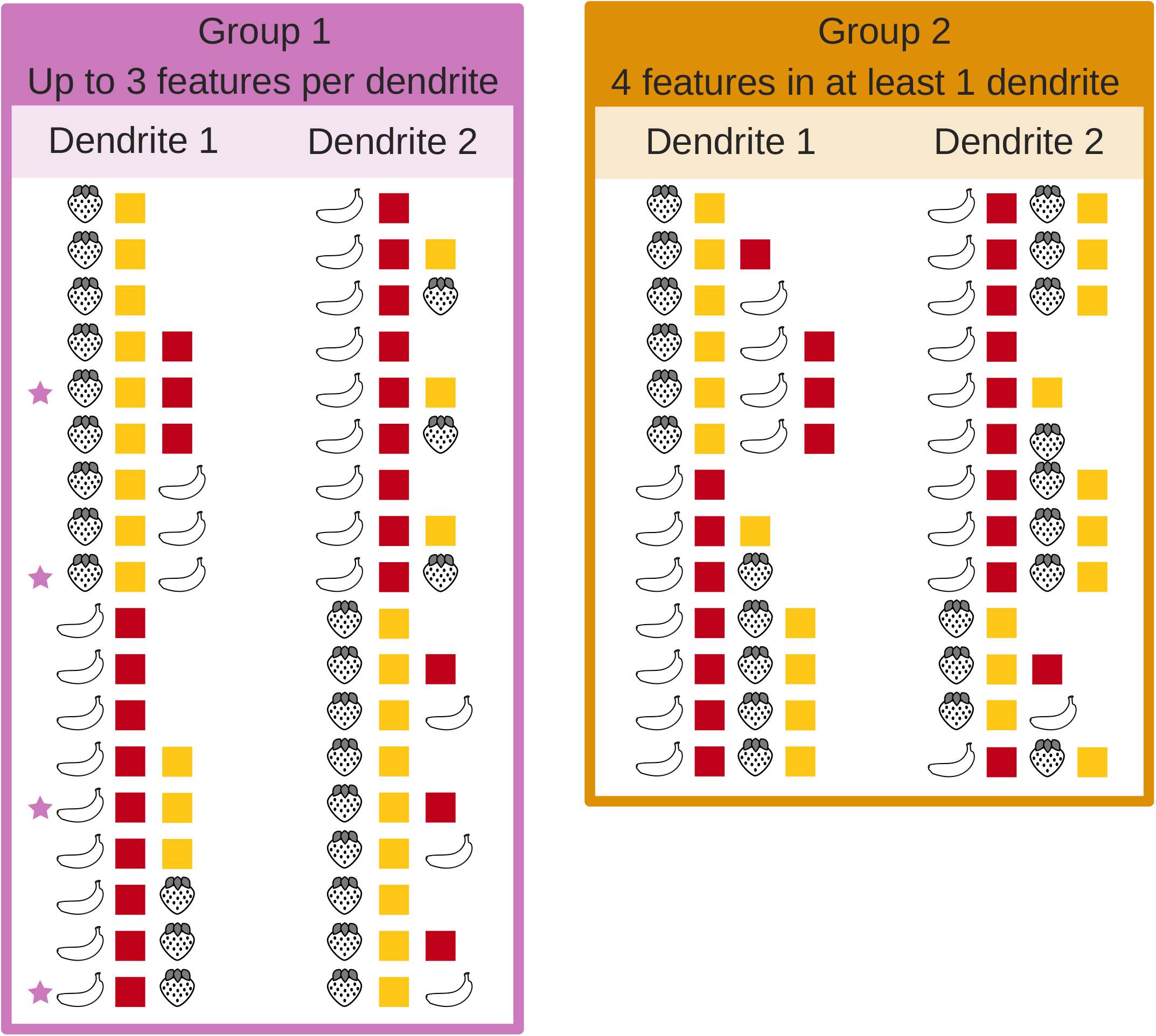
All possible input configurations of four features on two dendrites where one initially relevant pattern and both irrelevant patterns can be stored through in the simulations with reversal learning. They are divided in two groups: those with up to 3 features per dendrite (group 1), and those with 4 features in at least 1 dendrite (group 2). This division is made because when a dendrite is innervated with all four features (group 2), learning to solve the NFBP depends on the order in which the patterns arrive and solving the NFBP requires additional mechanisms, such as branch plasticity or inhibition (***Legenstein and Maass, 2011; Trpevski et al., 2026)***.

**Figure 10—figure supplement 2.**
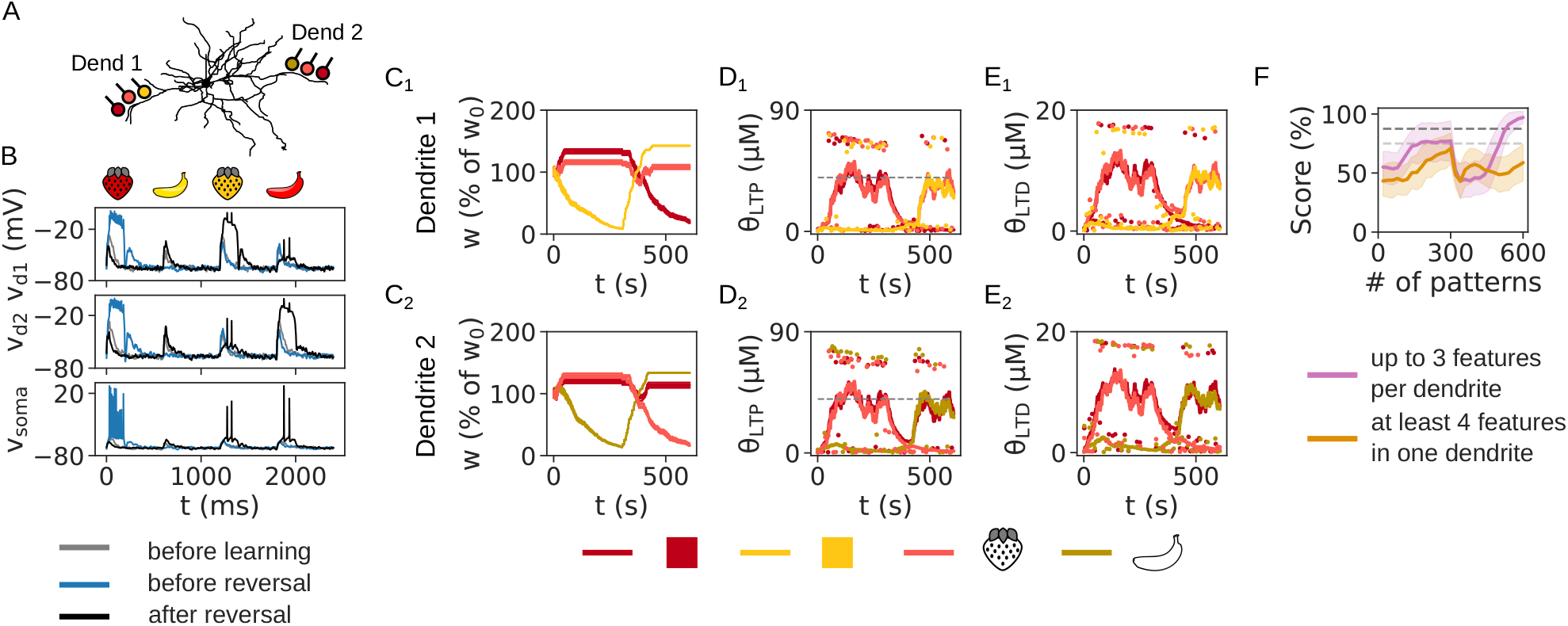
Relaxed metaplasticity also solves the NFBP in input configurations that can store both (initially) irrelevant patterns. (A) The dendritic and somatic voltage before learning (gray traces), before reversal (blue traces) and after reversal (black traces). (B–D) The evolution of synaptic weights (B), LTP thresholds (C) and LTD thresholds (D) during learning. In (B) all synaptic weights are shown, and in (C, D) the thresholds for only one synapse per feature are shown. (E) The performance on the NFBP in the input configurations that can store one relevant pattern and both irrelevant patterns (shown in Figure 10–figure supplement 1). 12 trials per input configuration were used, divided into the two groups shown in Figure 3–figure supplement 1, with 216 and 156 trials, respectively).

**Figure 10—figure supplement 3.**
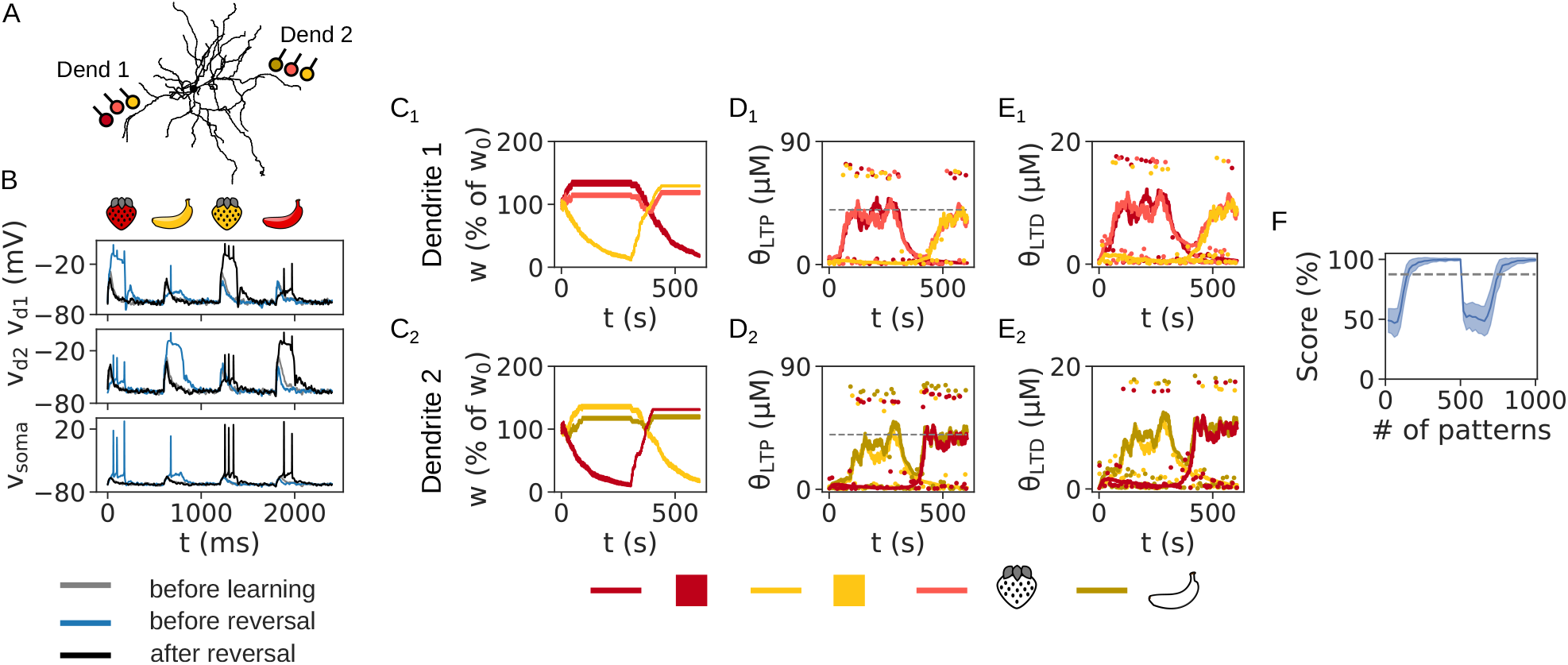
Reversal learning in the NFBP on one input configuration that can store both relevant and both irrelevant patterns. (A) An input configuration that allows the initially relevant patterns ‘red strawberry’ and ‘yellow banana’ to be stored in dendrite 1 and 2, respectively, as well as, the initially irrelevant patterns ‘yellow strawberry’ and ‘red banana’ to be stored in dendrites 1 and 2 after reversal of the reward policy. (B) The somatic and dendritic voltage before learning (gray traces), before reward policy reversal (blue traces) and after reversal (black traces).(C–E) The evolution of synaptic weights (C), LTP thresholds (D) and LTD thresholds (E) during learning. In (C) all synaptic weights are shown, and in (D, E) the thresholds for only one synapse per feature are shown. (F) The performance on the NFBP in all four input configurations in Figure 3– figure supplement 1 that can store both relevant and irrelevant patterns. 32 trials were performed for each input configuration, for a total of 128 trials.

**Figure 12—figure supplement 1.**
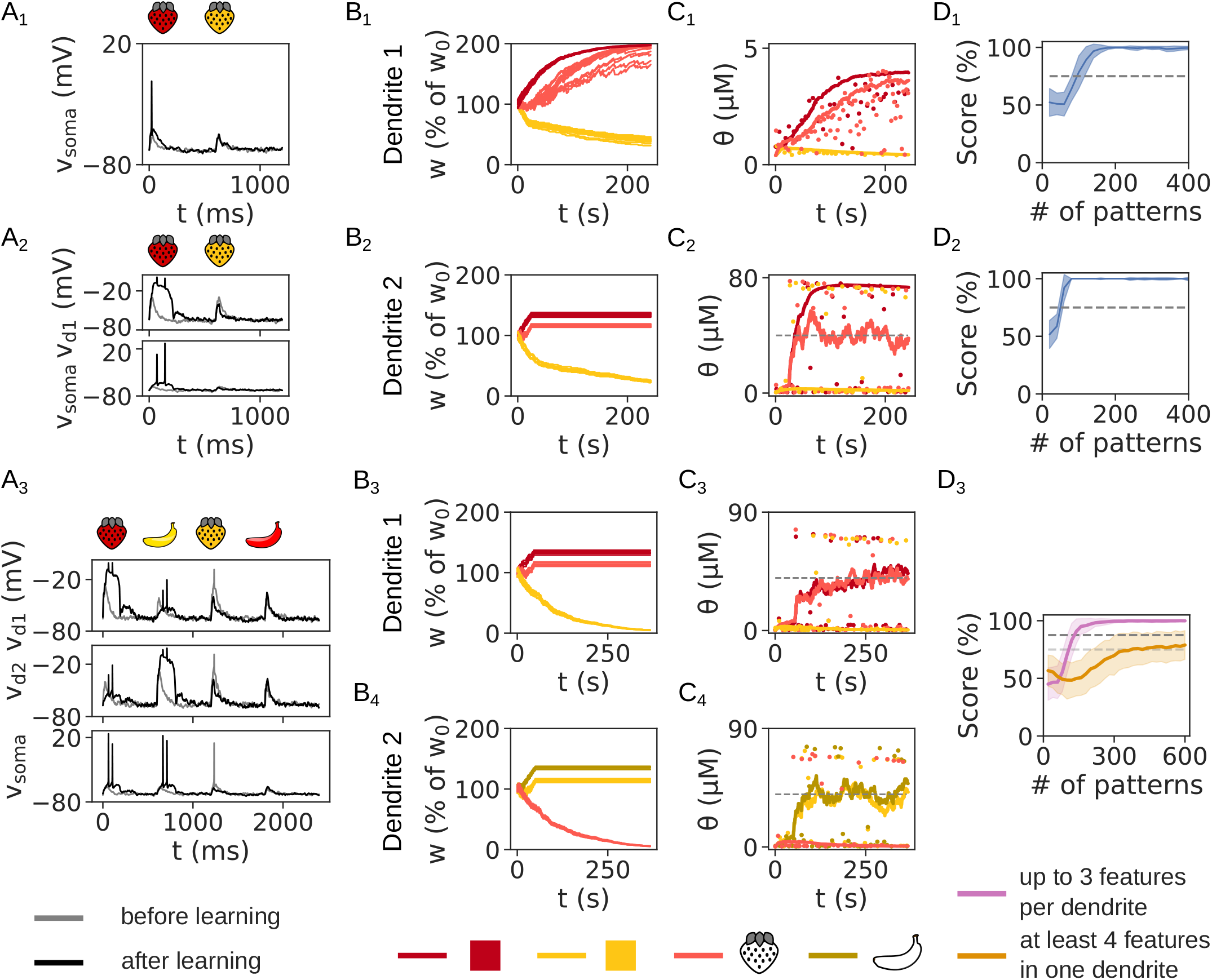
A single plasticity threshold is sufficient to solve the FBP and NFBP when the conditions for metaplasticity are relaxed. (A) The dendritic and somatic voltage before (gray traces) and after learning (black traces) for the FBP with linear (A_1_) and supralinear integration (A_2_) and the NFBP (A_3_). (B, C) The evolution of synaptic weights (B) and the single calcium threshold (C) for the synapses in each dendrite. Dashed lines in (C) show the upper LTP threshold, Θ_LTP_. In (B) all synapses are shown, and in (C) only one synapse per feature is used to show its calcium threshold. Dots represent the [Ca] amplitudes during pattern presentation, omitting some patterns for clarity. (E) The performance on the FBP with linear (E_1_) and supralinear integration (E_2_) and the NFBP (E_3_). Performance on the FBP is averaged over 50 trials, and in the NFBP over 12 trials per input configuration (divided into the two groups shown in Figure 3–figure supplement 1, with 216 and 156 trials, respectively).

**Figure 12—figure supplement 2.**
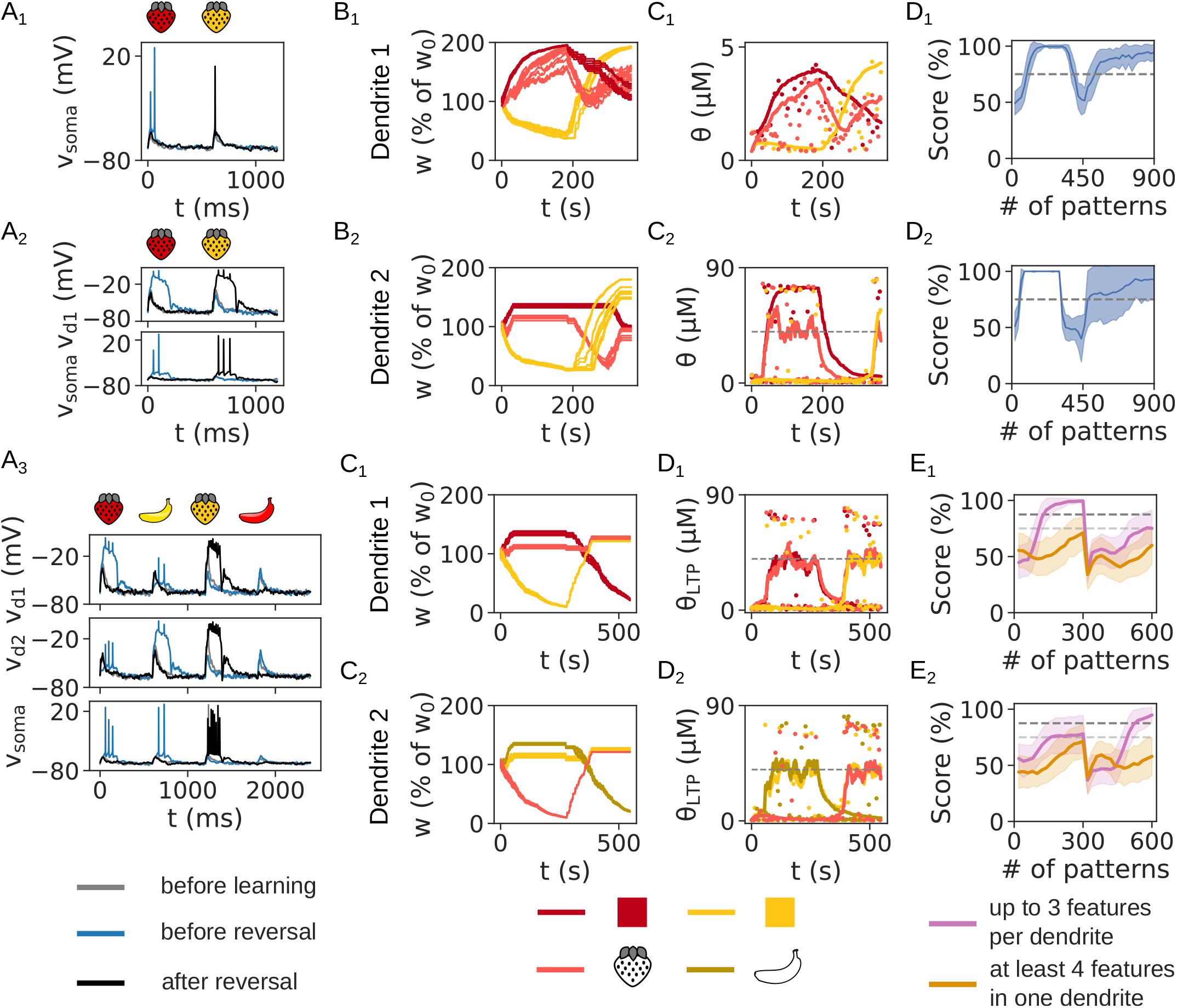
A single plasticity threshold is sufficient to solve reversal learning when the conditions for metaplasticity are relaxed. (A) The dendritic and somatic voltage before learning (gray traces), before reward policy reversal (blue traces) and after reversal (black traces) for the FBP with linear (A_1_) and supralinear integration (A_2_) and the NFBP (A_3_). (B, C) The evolution of synaptic weights (B) and the single calcium threshold (C) for the synapses in each dendrite. Dashed lines in (C) show the upper LTP threshold, Θ_LTP_. In (B) all synapses are shown, and in (C) only one synapse per feature is used to show its calcium threshold. Dots represent the [Ca] amplitudes during pattern presentation, omitting some patterns for clarity. (D) The performance on the FBP with linear (E_1_) and supralinear integration (E_2_) and the NFBP (E_3_, E_4_). Performance on the FBP is averaged over 50 trials, and in the NFBP over 12 trials per input configuration (divided into the two groups shown in Figure 3–figure supplement 1, with 216 and 156 trials, respectively).

**Figure12—figure supplement 3.**
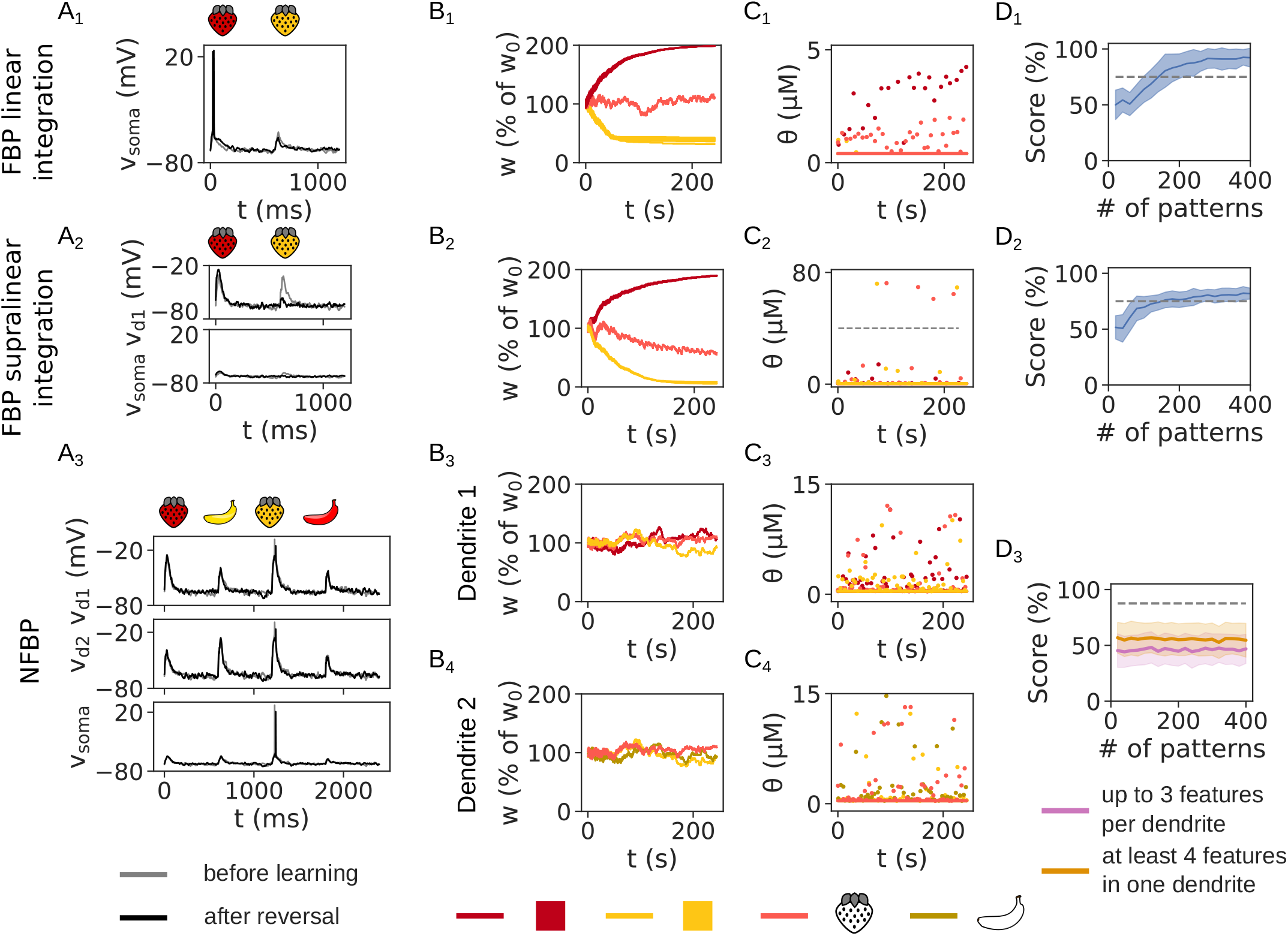
Learning outcome without metaplasticity in the single threshold scenario (the threshold is fixed). Similarly to when two thresholds are used, only the FBP is solved with reduced performance. (A) The dendritic and somatic voltage before (gray) and after learning (black) for the FBP with linear (A_1_) and supralinear integration (A_2_) and the NFBP (A_3_). (B) The evolution of synaptic weights. (C) The single calcium threshold is fixed. Dashed lines show the upper LTP threshold, Θ_LTP_, and only one synapse per feature is used to show its calcium threshold. Dots represent the [Ca] amplitudes during pattern presentation, omitting some patterns for clarity. (D) The performance on the FBP with linear (D_1_) and supralinear integration (D_2_) and the NFBP (D_3_, E_4_). Performance on the FBP is averaged over 10 trials, and in the NFBP over 5 trials per input configuration (divided into the two groups shown in Figure 3–figure supplement 1, with 216 and 156 trials, respectively).

**Figure 12—figure supplement 4.**
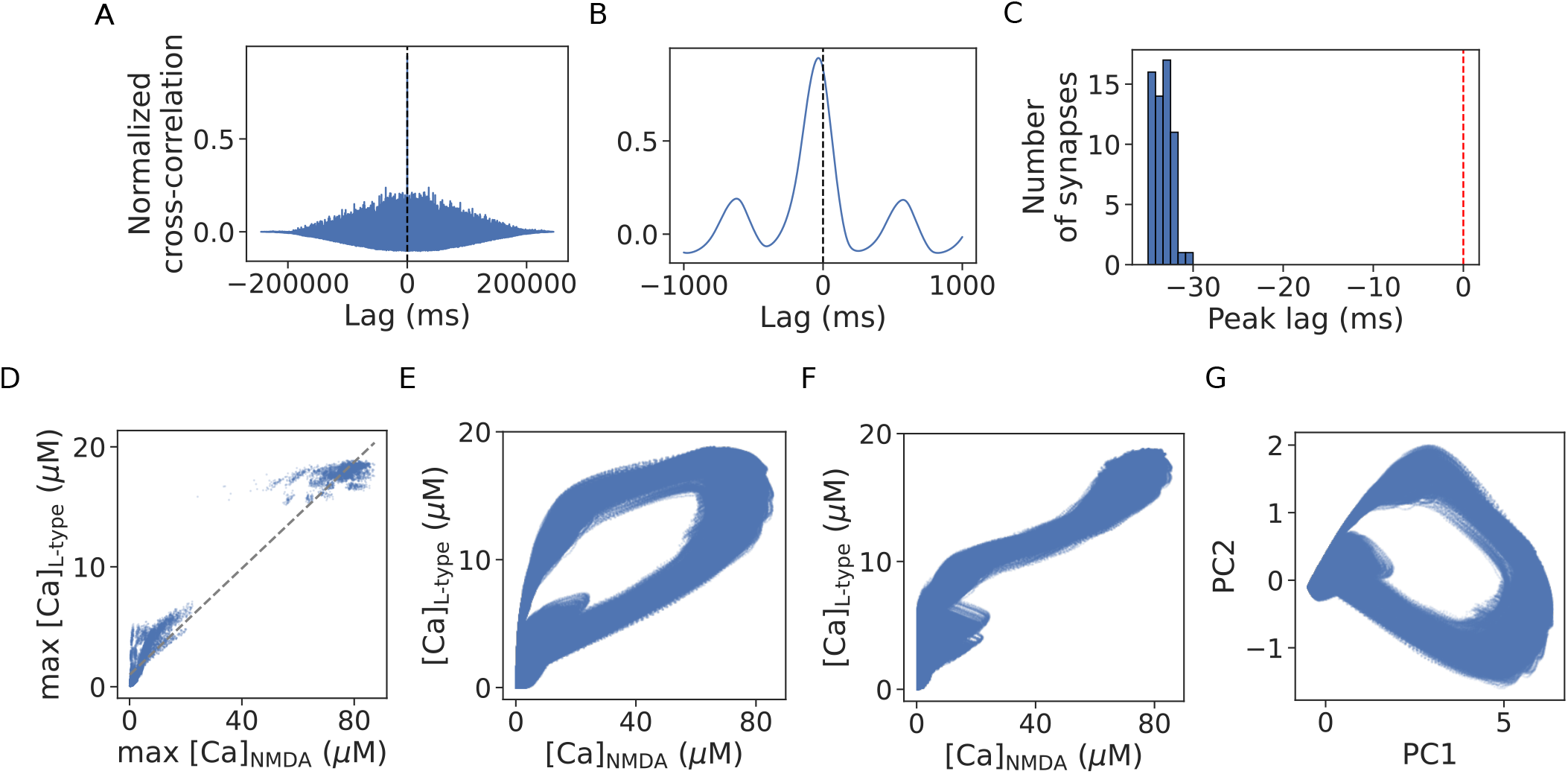
[Ca]_NMDA_ and [Ca]_L-type_ are highly correlated and carry essentially the same information. The analysis is done on calcium signals from one learning simulation of the NFBP with “thresholded” metaplasticity. (A) Cross-correlation of the normalized [Ca]_NMDA_ and [Ca]_L-type_ in a single synapse, averaged over all clustered synapses. (B) Zoom-in on the crosscorrelation function shows that the peak is centered sligthly off 0, at around -35 ms. (C) Histogram of the peak cross-correlation lag suggests that [Ca]_NMDA_ precedes [Ca]_L-type_ by about 35 ms. (D) A scatter plot of the amplitudes of [Ca]_NMDA_ and [Ca]_L-type_ (whose values are used to check if the calcium thresholds are crossed in the learning rule), with a fitted linear regression model. The coefficient of determination is *R*^2^ ≈ 0.98141, so in the linear regression model, max[Ca]_NMDA_ explains 98.14% of the variance in max[Ca]_L-type_. Also, Pearson’s correlation coefficient for the calcium amplitudes is *r* ≈ 0.99066, indicating that they are highly correlated. (Note that in this case, *r*^2^ = *R*^2^). (E) A scatter plot of the entire [Ca]_NMDA_ and [Ca]_L-type_ time-courses in all clustered synapses. It shows a circular structure, in agreement with the fact that [Ca]_NMDA_ precedes [Ca]_L-type_. (F) A scatter plot of the entire [Ca]_NMDA_ and [Ca]_L-type_ time-courses in all clustered synapses where [Ca]_L-type_ is shifted by 35 ms in order to be aligned with [Ca]_NMDA_. The time-courses also have a pronounced linear relationship. (G) Principal component analysis of the normalized [Ca]_NMDA_ and [Ca]_L-type_ time-courses ([Ca]_L-type_ was not shifted by any amount). PC1 explains around 94.26 % of the variance in the two calcium signals, suggesting that the joint calcium dynamics are strongly dominated by a onedimensional variable (the membrane voltage). (PC2 explains 5.74 % of the variance.)

**Figure 13—figure supplement 1.**
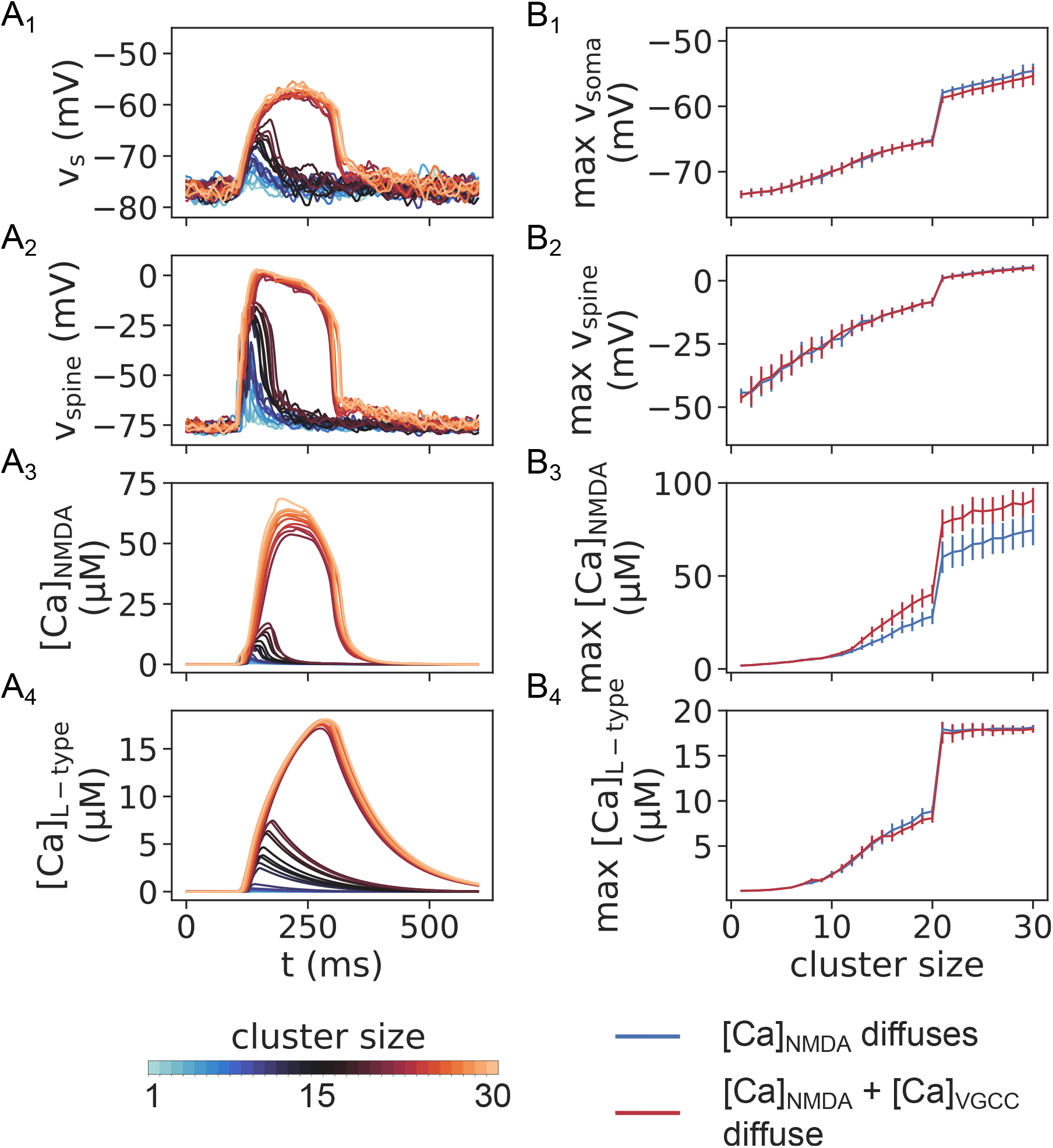
Voltage and calcium elevations when only [Ca]_NMDA_ diffuses axially through the SPN. (A_1_-C_4_) Somatic voltage, spine voltage, spine [Ca]_NMDA_ and spine [Ca]_L-type_ evoked in the supralinear integration scenario, to be compared with Fig. 2C (synaptic cluster placed on one dendrite, in a 20-micrometer region approximately 120 micrometers away from the soma). Line color represents the size of the synaptic cluster, which is varied from 1 to 30 synapses. (B_1_-B_4_) The amplitude of the somatic voltage, spine voltage, spine [Ca]_NMDA_ and [Ca]_L-type_ compared between the cases of only [Ca]_NMDA_ diffusing and [Ca]_NMDA_ + [Ca]_VGCC_ diffusing (the latter is replfrom Fig. 2D). Results are averages over 20 trials, and in the case of synaptic clusters, over 8 different dendrites (clusters located in a 20-micrometer in a dendritic region starting at approximately between 120 micrometers from the soma).

